# Linking the gut microbiome to host DNA methylation by a discovery and replication epigenome-wide association study

**DOI:** 10.1101/2023.11.03.565420

**Authors:** Ayşe Demirkan, Jenny van Dongen, Casey T. Finnicum, Harm-Jan Westra, Soesma Jankipersadsing, Gonneke Willemsen, Richard G. Ijzerman, Dorret I. Boomsma, Erik A. Ehli, Marc Jan Bonder, Jingyuan Fu, Lude Franke, Cisca Wijmenga, Eco J.C. de Geus, Alexander Kurilshikov, Alexandra Zhernakova

## Abstract

Both gene methylation and the gut microbiome are partially determined by host genetics and partially by environment. We investigated the relations between gene methylation in blood and the abundance of common gut bacteria profiled by 16s rRNA gene sequencing in two population-based Dutch cohorts: LifeLines-Deep (LLD, n = 616, discovery) and the Netherlands Twin Register (NTR, n = 296, replication). In LLD, we also explored microbiome composition using data generated by shotgun metagenomic sequencing (n = 683). We then investigated if genetic and environmental factors can explain the methylation–microbiota associations in a set of 78 associated CpG–taxa pairs from the EWAS meta-analysis. In both cohorts, blood and stool samples were collected within 2 weeks of each other. Methylation was profiled in blood samples using the Illumina 450K array. Methylation and microbiome analysis pipelines were harmonized across cohorts. Epigenome-wide association study (EWAS) of microbial features were analysed using linear regression with adjustment for technical covariates.

Discovery and replication analysis using 16s data identified two independent CpGs associated with the genus *Eggerthella*: cg16586104 (P_meta-analysis_ = 3.21 × 10^-11^) and cg12234533 (P_meta-analysis_ = 4.29 × 10^-10^). While we did not find human genetic variants that could explain the associated CpG–taxa/pathway pairs, we show that microbiome can mediate the effect of environmental factors on epigenetics.

In this first association study linking epigenome to microbiome, we found and replicated the associations of two CpGs to the abundance of genus *Eggerthella* and identified microbiome as a mediator of the exposome.

## Introduction

The gut microbiome is now widely accepted to be a modifiable factor that is associated with a wide range of host health outcomes. It has been repeatedly shown to express its effects not only within the intestinal system, e.g. in colorectal cancer^1^ (CRC) and inflammatory bowel disease^2^ (IBD), but also at the systemic level, for instance in metabolic disorders such as type 2 diabetes^3^ or neuropsychiatric conditions^4^ such as Parkinson’s disease^5^ and mood disorders. Causal roles for the gut microbiome have been proven for some of these associations, and the underlying mechanisms include short-chain fatty acid production by bacteria^6^, stimulation of the vagus nerve^7^ or via the enteroendocrine cells^8^, or microbial production of triggers for inflammatory pathways, such as lipopolysaccharides^9^. The gut microbiome is assumed to be shaped primarily by the exposome and secondarily by host genetics^10^, but the contribution of the host epigenome and its relation between environmental factors (such as diet) and microbiome, has never been studied in human cohorts at large scale. Thus the mechanistic links for the recently suggested “Microbiota ↔ Nutrient Metabolism ↔ Host Epigenome” model are far from being understood^11^.

Host epigenetics is mostly studied by capturing DNA methylation at CpG dinucleotides over the human genome. Similar to the microbiome, methylation can be modified by both host genetics and environment^12^. Modification of CpG markers can be responsible for switching gene expression genes on or off during development^13^ and throughout life. In addition to genetic and environmental control for this modification, it can also be influenced by early life events^14,15^ and maternal factors^16,17^. Multiple epigenome-wide association studies (EWAS) have identified links between differential CpG methylation in blood cells and cardiovascular^18^, metabolic^19,20^, psychiatric^21^ outcomes and cancer^22^. In line with this, several EWAS have also linked CpG methylation to environmental exposures, e.g. air quality^23^, stress^24^, occupational exposure to chemicals^25,26^, prescribed medications^27^, dietary habits such as coffee and alcohol consumption^28,29^ and supplement intake^30^.

Here, we performed an EWAS of the gut microbiome to test whether gut microbial abundances associate to differentially methylated CpGs measured in blood. We performed this study in a discovery and replication setting in two cohorts from the Netherlands: LifeLines-DEEP (LLD, n = 616) and the Netherlands Twin Register (NTR, n = 296). Additionally, using metagenomic shotgun sequencing (MGS) data generated in the LLD cohort, we obtained the abundances of bacterial metabolic pathways and microbial species and linked these with host DNA methylation (n = 683). Finally, as gut microbial abundances are, in part, under the control of host genetics^10^, environment^31^, diet^32^ and medication use^33^, we elucidated whether the observed associations can be explained by any of these factors and searched for a mediating role for the gut microbiome in these associations.

## Methods

### Cohorts

LLD is a subcohort of Lifelines. Lifelines is a multi-disciplinary prospective population-based cohort study examining, in a unique three-generation design, the health and health-related behaviors of 167,729 persons living in the North of the Netherlands. It employs a broad range of investigative procedures to assess the biomedical, socio-demographic, behavioral, physical and psychological factors that contribute to the health and disease of the general population, with a special focus on multi-morbidity and complex genetics. Blood and fecal samples of LLD participants were collected between April and August 2013. Fecal DNA was extracted using the Qiagen AllPrep kit with a bead-beating step. Sequencing of the bacterial 16s rRNA gene, domain V4, was performed at the Broad Institute (Boston, USA) using the Illumina MiSeq platform, as described in^10^. Metagenomics sequencing of the same DNA samples was performed at the Broad Institute, as described in_31._

The NTR collects longitudinal data in twin families^34^. Biological samples are collected in the NTR-Biobank^35,36^. The NTR samples included in our microbiome EWAS were collected for two separate studies. The first focused on the association between obesity and the gut microbiome^37^ and the second collected samples from family members and spouses^38^. Fecal DNA was extracted using the Qiagen PowerSoil kit with the addition of the heating step from the protocol of the Qiagen PowerFecal kit. The sequencing of the V4 domain of the 16S gene was performed using the Illumina MiSeq platform, as described in ^10^. DNA extraction and sequencing were performed at the Avera Institute for Human Genetics (Sioux Falls, SD, USA).

### Analysis of 16s data

In both cohorts, a standardized 16s processing pipeline (https://github.com/alexa-kur/miQTL_cookbook) was used to characterize the V4 region using the RDP classifier^39^ over the SILVA128 database^40^. We used the 16s rRNA–based microbiome profiling from both cohorts, the data of which were harmonized earlier as part of a large meta-analysis^10^. MGS was only performed in LLD and was analyzed using the Metaphlan v.2 and Humann2 algorithms with the aim of yielding higher taxonomic resolution and functional insights^31^.

### Methylation

In both cohorts, genome-wide DNA methylation in whole blood was analyzed using the Infinium HumanMethylation450 BeadChip Kit^41^. Genomic DNA (500ll!ng) from whole blood was bisulfite-treated using the ZymoResearch EZ DNA Methylation kit (Zymo Research Corp, Irvine, CA, USA), following the standard protocol for Illumina 450K micro-arrays, at the Department of Molecular Epidemiology of Leiden University Medical Center, the Netherlands. In short, subsequent steps (sample hybridization, staining and scanning) were performed by the Human Genomics Facility (HuGe-F) at the Erasmus Medical Center, Rotterdam, the Netherlands. The resulting data were processed in accordance with BIOS consortium guidelines, as detailed earlier^17,42^. Sample-level quality control was performed using MethylAid^43^. Probes were set to be missing in a sample if they had an intensity value of exactly zero, a detection P-value > .01 or a bead count < 3. After these steps, the probes that failed the above criteria in >5% of the samples were excluded from all samples, leaving only the probes with a success rate ≥0.95. Probes were also excluded from all samples if they were mapped to multiple locations in the genome or had a single nucleotide polymorphism (SNP) within the CpG site (at the C or G position), irrespective of the minor allele frequency in the Dutch population^41^. Only autosomal methylation sites were analyzed in the EWAS. The methylation data were normalized with functional normalization, as implemented in *minfi*^44^. In both cohorts, blood and stool samples were collected within 2 weeks of each other.

### Statistical analysis

We calculated methylation M-values as the logarithm of methylated over unmethylated probe intensity ratio. As M-values were the outcome in our regression models, we further excluded outliers, which we defined as values lying outside of the ± 3.5 interquartile range for each methylation probe. We performed association tests in LLD (which has no related individuals) using linear regression models adjusted for age, gender, sample plate, position on plate, blood cell counts, smoking and the first three genetic principal components. Genome-wide inflation in each EWAS was corrected using the R package “*Bacon*”. In the replication cohort NTR, generalized estimation equation (GEE) models were fitted using the R package “*gee*” to enable inclusion of both the twins in the analysis, controlling for kinship. Covariates in the GEE models in NTR include age, sex, sample plate, array row, blood cell counts and smoking. We set a minimum number of 50 available data points (samples with both CpG and microbial trait abundance data available) as a criterium to be included in the EWAS analysis, and 223 taxa and 410K CpG markers were ultimately included in the analysis. For the EWAS of the 16s rRNA gene data, the final analytical set was 616 individuals for LLD and 296 individuals for NTR.

In the EWAS of the LLD MGS data (683 individuals), we analyzed 209 bacterial taxa and 326 bacterial pathways. In the single cohort analysis the experiment wide p-value was considered significant at 1.08 × 10^-9^ for 16S, 7.36 × 10^-10^ for bacterial pathways and 1.15 × 10^-9^ for MGS-derived taxa abundances. These numbers are based on the EWAS-sign threshold of 2.4 × 10^-7^ for a single experiment performed on 450k array ^45^ and according to the Bonferroni procedure for multiple testing correction. For meta-analysis, only discovery set associations with P < 1 × 10^-4^ were included in the meta-analysis.

We searched EWAS datahub^46^ and EWAS atlas^47^ for existing epidemiological evidence for the identified CpGs. We used the BIOS meQTL/eQTM atlas (http://bbmri.researchlumc.nl/atlas/#data, files: “Cis-meQTLs independent top effects” and “Cis-eQTMs independent top effects”) to identify genetic determinants of CpGs of interest^48,49^ and their correlated gene expression. Genetic determinants for bacterial abundance were extracted from MiBioGen GWAS results^10^. Phenome-wide associations for selected meQTLs were extracted from GWASATLAS^50^. Exposome data was initially available for 1135 individuals in the larger LLD cohort and included information about diet and medication. Association to these exposome exposures was performed in the LLD dataset, including 60 dietary preferences and 22 prescription medications (medications with >10 users) in 870 people for 16s, in 1124 people for MGS and in 689 people for host gene-methylation data.

## Results

### EWAS of gut microbiome profiled by 16s gene sequencing, discovery and replication

Figure 1 shows a schematic presentation of the study design. In the discovery EWAS of 16s-derived taxonomy performed in 602 participants of the LLD cohort, we identified 3520 CpG– taxa pairs with a P_discovery_ < 1.00 × 10^-4^ and tested their association in the replication set of 296 samples from NTR. This revealed that two independent CpGs, cg16586104 (P_replication_ = 1.25 × 10^-11^, P_meta-analysis_ = 3.21 × 10^-11^) and cg12234533 (P_replication_ = 2.87 × 10^-6^, P_meta-analysis_ = 4.29 × 10^-10^) were associated with the abundance of genus *Eggerthella* (Table 1). The methylation M-value of cg16586104, located ∼600kb from the closest gene*RWDD3 (chr1p21.3)*, was associated with increased abundance of genus *Eggerthella* (methylation level positively correlated with abundance). The M-value of cg12234533, located inside *ULK4* (unc-51-like serine/threonine kinase, *chr3p22.1*), was associated with decreased abundance of *Eggerthella*. Additionally, we selected 76 CpG–taxa pairs with a suggestive P_meta-analysis_ < 1.00 × 10^-4^ for further investigation in relation to exposome, so that 78 CpG– taxa pairs from the 16s EWAS were followed-up (TableS1, FigureS1 and S2).

**Figure 1:**
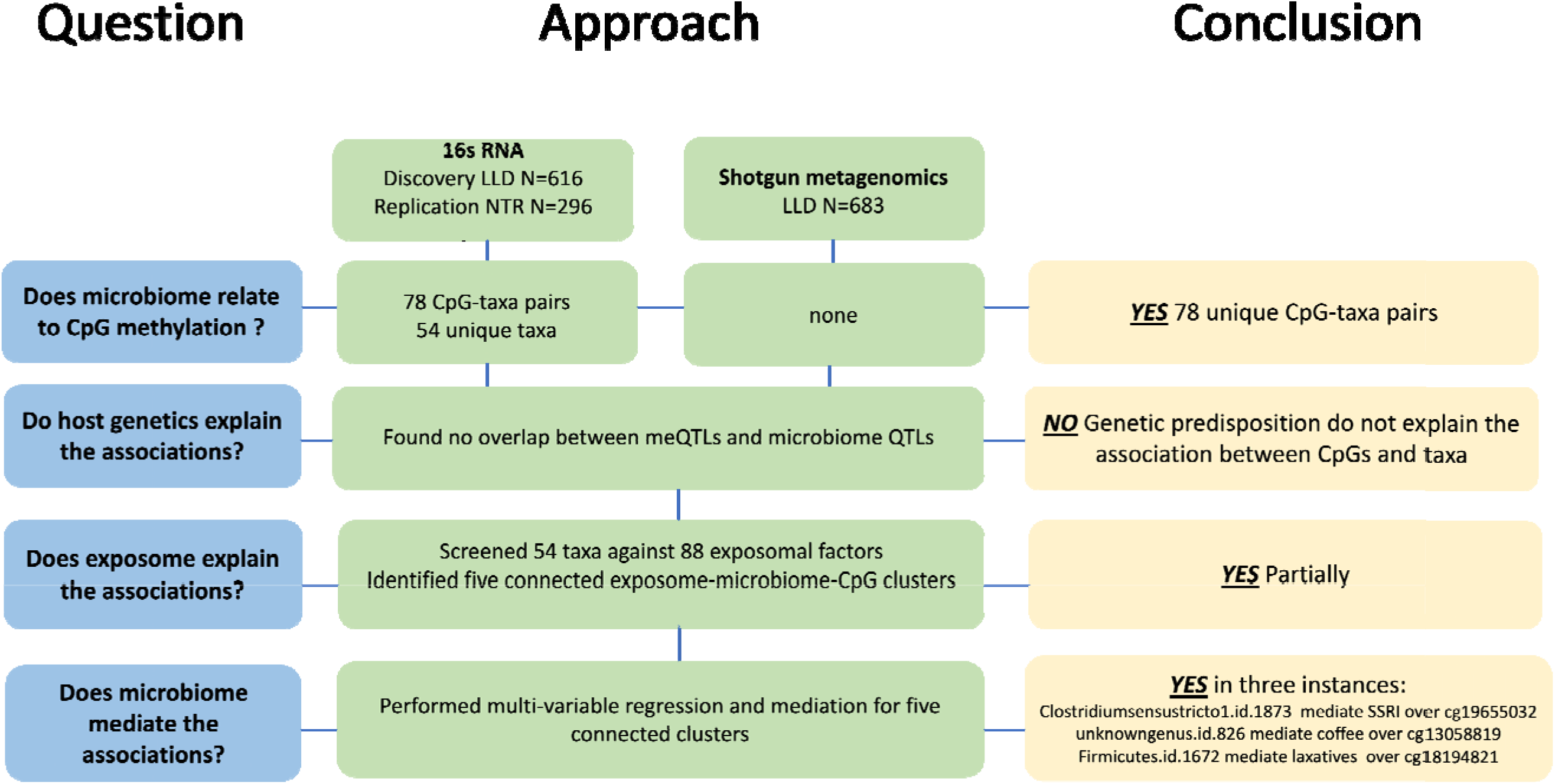
Schematic representation of the study.

**Figure 2.**
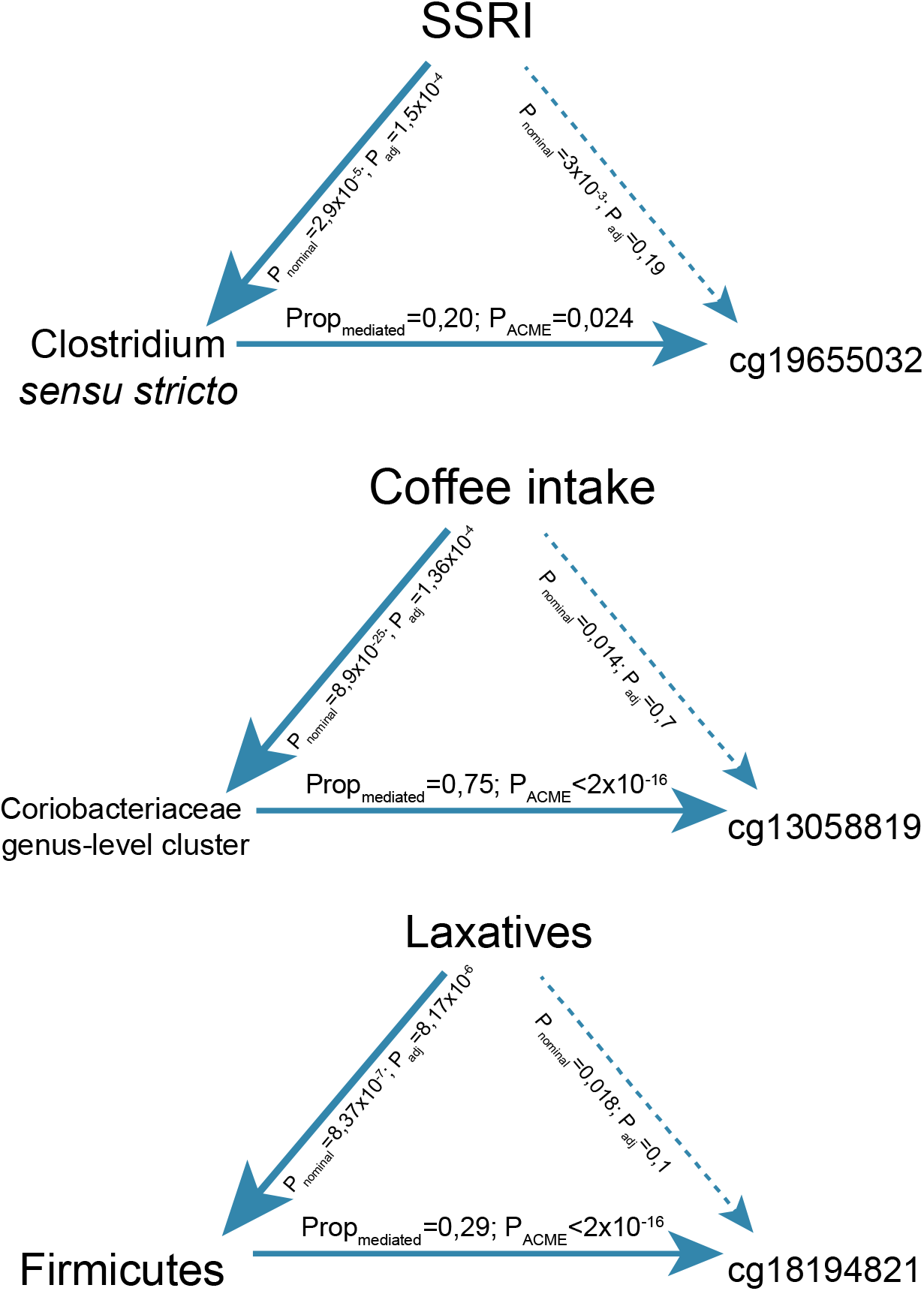
Mediation analysis of environmental exposures, microbiome and CpG methylation. Figure 2. Mediation analysis of environmental exposures, microbiome and CpG methylation. P_nominal_ represents the p-value of association of environmental exposure to microbial trait and CpG. P_adj_ represents the conditional association of environmental exposure to both microbiome and methylation traits, adjusted for each other. Prop_mediated_ represents the proportion of environmental effect on CpG methylation which is mediated by microbial trait. P_ACME_ represents the significance of Average Causal Mediation Effects (ACME).

**Table 1.**
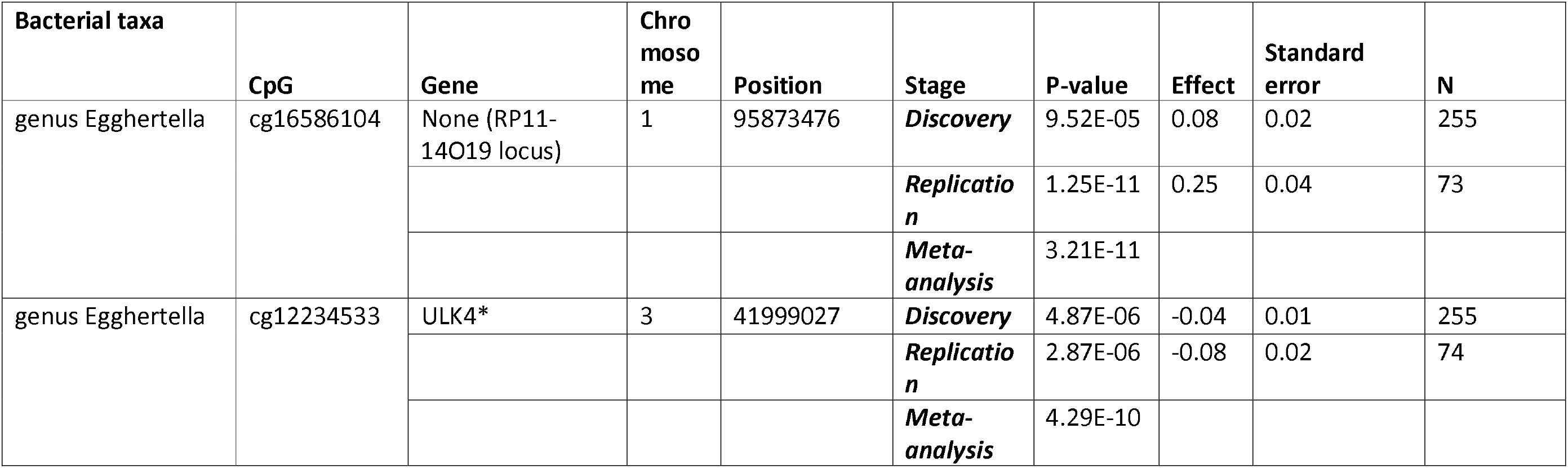
CpGs associated with bacterial taxa. Table 1 shows CpGs associated with bacterial taxa. Two CpGs that were selected from the 16S discovery EWAS (LLD cohort) with P_discovery_ <10^-4^ and replicated in the NTR with P_replication_<1.42 × 10^-5^. The CpGs were not associated with any disease or human traits in the EWAS datahub (https://ngdc.cncb.ac.cn/ewas/datahub access date: 07/01/2022). *Other CpGs in ULK4 gene have been associated with smoking, preterm birth, glucocorticoid exposure, down syndrome, systemic lupus erythematosus. P-value: two sided type 1 error rate of null hypothesis assuming effect estimate of bacterial abundance equals zero in the regression model where methylation M-value is the outcome and bacterial abundance, age, sex and smoking and technical covariates included. N: Number of observations included in the analyses.

### EWAS of metagenomics-derived abundance and pathway profiles

In contrast to 16s rRNA gene sequencing, MGS allows for accurate identification of bacterial species and analysis of bacterial pathways. MGS was available for one cohort, LLD (683 individuals). None of the CpG-species/pathways pairs were associated at EWAS-wide significant levels. We further selected the suggesting signals for the follow-up and enrichment analyses. Twenty-three MGS-derived taxa that we found associated with host DNA methylation in the discovery-only EWAS, with suggestive P-value significance level (2.4 × 10^-7^ > P > 1.25 × 10^-9^). (Table S2). Ten of these 23 CpGs had been linked to other phenotypic outcomes by previous EWAS (Table S2). We also found eleven independent CpG–bacterial pathway associations that reached suggestive significance (2.4 × 10^-7^ > P > 7.34 × 10^-10^). (Table S3, Figure S3 and S4).

### Genetic effects shared between host DNA methylation and gut microbiome

We next examined genetic control over microbial abundance and gene methylation. We searched for the genetic determinants of 78 *unique* CpGs from the 16s rRNA EWAS and their associated bacterial taxa (n = 54). Based on an earlier meQTL study from the BIOS consortium ^48^, we identified 68 unique meQTLs that act as genetic regulators of 27 of the 78 CpGs we investigated (TableS3). Seventeen of these meQTLS had earlier been found to be associated with several traits and diseases (TableS4). Three of them, for which the methylation was associated with an unknown genus from the *Coriobacteriaceae* family, were also associated to gastrointestinal diseases in several studies: rs11576137 (meQTL of cg13058819, located in *SLC35F3*), rs1736020 (meQTL of cg08706567, located in *MPL*) and rs1736135 (cg08706567, *MPL*) were associated with diverticular disease^55,56^, IBD^57,58^, ulcerative colitis^57^ and Crohn’s disease^59^ ^58^ (TableS4). As none of the associated bacterial taxa were shown to be under genetic control in the previously published meta-analysis^10^, we did not find any shared genetic loci simultaneously controlling host methylation and microbiome, indicating that the correlation we observe is not due to host genetics.

### Shared effects of environmental exposures on host DNA methylation and gut microbiome

We next tested whether shared exposure effects could explain the associations between host DNA methylation and abundance of bacterial taxa. We first tested the effect of the exposome (82 environmental variables: 60 on dietary intake and frequencies and 22 on medication intake) on the microbiome for the 54 unique microbial abundances that we selected by 16S meta-analysis. EWAS. In total, 52 exposure–taxa pairs passed the 5% FDR cutoff (Benjamini-Hochberg procedure, TableS6). For five of them, the related CpG was also associated with the exposure (P_nominal_ < 0.05, Table 2). In particular, we identified that use of SSRI-type antidepressants was negatively associated with abundances of both genus *Clostridium sensu stricto 1* and family *Peptostreptococcaceae* (P = 2.91 × 10^-5^ and P = 1.84 × 10^-5^, respectively), as well as with their related CpGs: cg19655032 (P = 1.35 × 10^-2^) and cg06372145 (P = 1.54 × 10^-2^). For dietary factors, shared microbiome/epigenetic associations were observed for dairy and coffee consumption: Genus *Ruminococcus gauvreauii group* and cg20400838 were associated with dairy consumption (P = 5.99 × 10^-4^ and P = 1.45 × 10^-2^, respectively). A genus-level cluster from *Coriobacteriaceae* family and its associated CpG cg13058819 were simultaneously associated with coffee consumption (P = 8.93 × 10^-25^ and P = 1.42 × 10^-2^, respectively). In conclusion, for 5 of the 52 exposure–taxa pairs selected, we identified environmental traits that are associated to both the microbiome marker and its linked CpG site, suggesting that the microbiome may have a mediator effect.

**Table 2.** Diet and medication use factors that associate simultaneously with bacterial taxa/pathways and host DNA methylation, forming seven exposure-microbiome-methylation clusters. Table 2 shows the results from initial exposome analysis. In these analysis either bacterial abundance/taxa or CpG M values were included as outcomes, and exposure as predictors in regression models, together with technical covariates. We first selected the BH-signifci ant microbiome-exposure pairs, and then tested whether they are significantly associated with CpGs, and selected a total of FIVE exposure-microbiome-methylation clusters. Table shows the results from two different tests. P-value: two sided type 1 error rate of null hypothesis assuming effect estimate of bacterial abundance equals zero in the regression model where methylation M-value is the outcome and bacterial abundance, age, sex and smoking and technical covariates included. N: Number of observations included in the analyses.

| Exposure | Effect of exposure on microbial taxa/pathways |  |  |  |  | Effect of exposure on host gene methylation |  |  |  |
| --- | --- | --- | --- | --- | --- | --- | --- | --- | --- |
|  | Taxa(outcome) | Effect | P-value | N total (N users/non users) | BH P-value | CpG (outcome) | Effect | P-value | N total (N users/non users) |
| laxatives | phylum.Firmicutes.id.1672 | -0.17 | 8.37E-07 | 875(18/857) | 6.86E-05 | cg18194821 | 0.19 | 0.018 | 689(13/676) |
| SSRI | genus.Clostridiumsensustricto1.id.1873 | -1.64 | 2.91E-05 | 829(22/807) | 3.02E-03 | cg19655032 | -0.15 | 1.35E-02 | 689(21/68) |
| coffee_log | genus.unknowngenus.id.826 | 0.51 | 8.93E-25 | 812 | 9.28E-23 | cg13058819 | 0.04 | 1.42E-02 | 689 |
| dairy_log | genus..Ruminococcusgavreaii group.id.11342 | 0.32 | 5.99E-04 | 784 | 3.11E-02 | cg20400838 | -0.09 | 1.45E-02 | 689 |
| SSRI | family.Peptostreptococcaceae.id.2042 | -1.38 | 1.84E-05 | 867(18/766) | 9.57E-04 | cg06372145 | -0.34 | 1.54E-02 | 689(21/68) |

### Mediator effect of the microbiome

We next tested whether the associations of microbiome/exposure to methylation CpGs are independent of each other. Table 2 shows the results from association models where CpG methylation is included as the outcome and microbial traits and environmental exposures are both included as predictors, along with technical covariates, age and gender. For three taxa–CpG–exposure clusters, SSRI-type antidepressant use (N = 17 users in the statistical model), laxative use (N = 12 users) and the amount of coffee consumption, the associations between the exposure and host gene methylation were weakened by inclusion of microbial effects in the models. For these three traits, the association with host gene methylation levels (CpGs cg19655032, cg13058819 and cg18194821, respectively) was lost when association is adjusted for microbial features, indicating that the effect of environment may be mediated by the microbiome. (Table 3).

**Table 3.**
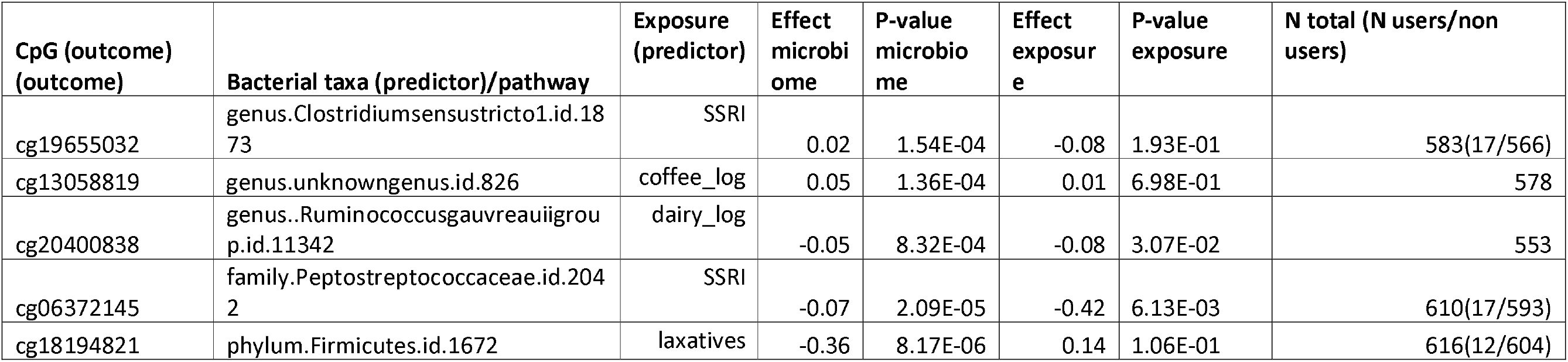
Effects of diet and medication estimated by adjusted models. Table 3 shows the results from adjusted linear regression analyses where both microbiome and exposure were included in the models together to associate with the outcome of CpG M values, along with age, sex and smoking and technical covariates.

| CpG (outcome)<br>(outcome) | Bacterial taxa (predictor)/pathway | Exposure<br>(predictor) | Effect<br>microbiome | P-value<br>microbiome | Effect<br>exposure | P-value<br>exposure | N total (N users/non users) |
| --- | --- | --- | --- | --- | --- | --- | --- |
| cg19655032 | genus.Clostridiumsensustricto1.id.1873 | SSRI | 0.02 | 1.54E-04 | -0.08 | 1.93E-01 | 583(17/566) |
| cg13058819 | genus.unknowngenus.id.826 | coffee_log | 0.05 | 1.36E-04 | 0.01 | 6.98E-01 | 578 |
| cg20400838 | genus..Ruminococcusgavreuiigroup.id.11342 | dairy_log | -0.05 | 8.32E-04 | -0.08 | 3.07E-02 | 553 |
| cg06372145 | family.Peptostreptococcaceae.id.2042 | SSRI | -0.07 | 2.09E-05 | -0.42 | 6.13E-03 | 610(17/593) |
| cg18194821 | phylum.Firmicutes.id.1672 | laxatives | -0.36 | 8.17E-06 | 0.14 | 1.06E-01 | 616(12/604) |

We next used formal mediation analysis to test the role of the microbiome in mediating the effects of the three exposures (SSRI-type antidepressants, laxative use and coffee consumption) on gene methylation and calculated the Average Causal Mediation Effects, ACME (Table 4). For all three clusters, we found significant ACME for the microbiome mediating the effects of the exposure on host gene methylation. Genus *Clostridium sensu stricto 1* mediates the effects of SSRI-type antidepressants on cg19655032 (ACME = −0.0216, P = 0.024, proportion of effect mediated = 0.20), a genus-level cluster from family *Coriobacteriaceae* mediates the effects of coffee exposure on cg13058819 (ACME = 0.02245, P < 2 × 10^-16^, proportion of effect mediated = 0.75) and phylum Firmicutes mediates the effects of laxative use on cg18194821 (ACME = 0.055, P < 2 × 10^-16^, proportion of effect mediated = 0.28). These examples show that microbiome can mediate the effect of environment on DNA methylation.

**Table 4.** Mediation analysis. Table 4 shows the results from mediation analysis for the clusters selected from adjusted analysis in Table 3. ACME: stands for Average Causal Mediation Effects.

| Outcome | Exposure | Mediator | ACME | ACME p-value | Total effect | Total effect P-value | Prop. Mediated |
| --- | --- | --- | --- | --- | --- | --- | --- |
| cg19655032 | SSRI | genus.Clostridiumsensustricto1.id.1873 | -0.0216 | 0.024 | -0.1053 | 0.056 | 0.2047 |
| cg13058819 | coffee_log.y | genus.unknowngenus.id.826 | 0.02245 | <2e-16 | 0.02988 | 0.08 | 0.75139 |
| cg18194821 | laxatives | phylum.Firmicutes.id.1672 | 0.0552 | <2e-16 | 0.1904 | 0.13 | 0.2897 |

## Discussion

We performed an association study between the gut microbiome and host epigenome in 912 individuals from two independent Dutch cohorts. For all samples, blood methylation and gut microbiome were analyzed at the same timepoint (within 2 weeks) and the same pipelines for the analysis of microbiome and methylation data were applied. We identified study-wide association of CpG methylation with one bacterial taxa,*Eggerthella, which was associated with two independent CpGs*. Exploring the effect of environmental factors on the microbiome and methylation identified that for a subset of associations the same exposome factors – coffee consumption, dairy intake and use of laxatives, PPIs and antidepressants – are associated to both the microbiome and methylation. We also show a potential role for three bacterial taxa in mediating the effects of coffee, SSRI antidepressants and laxatives on DNA methylation, although this analysis is limited by the small number of users and the cross-sectional nature of this study. We did not observe shared effect of host genetics on both microbiome and methylation, which is expected given the overall modest contribution of host genetics in regulating microbial abundance.

Genus *Eggerthella* is one of the gut microbiome genera we found to be associated with methylation in blood.*Eggerthella* is part of the normal human intestinal microbiome and has been most commonly associated with infections spreading from the gastrointestinal tract^60^, but it has also been found interacting with food intake while influencing metabolism of drugs^61^. This genus has also been shown to be more abundant in individuals with psychiatric diseases^62^ and higher grade neoplasms^63^. *E. lenta* from the same genus correlates with taurodeoxycholic acid, a bile acid metabolite, in the colon of smoke-exposed mice^64^ and has increased abundance in the presence of blood in stool, which is a marker for CRC^65^. However, we did not find any association with the related CpGs and the diseases mentioned above, although our search was limited to the EWAS studies carried out to date.

Following up on the association of a genus-level cluster under family *Coriobacteriaceae* with two independent methylation sites (cg13058819 and cg08706567), we came across*-cis* (rs11576137) and *-trans* (rs1736020 and rs1736135) meQTLs that are also genetic determinants of diverticular disease, IBD, ulcerative colitis and Crohn’s disease^50^, respectively. The *Coriobacteriaceae* family is known to have a decreased abundance in the gut of individuals with IBD^68^. In summary, we observe that these disease-associated variants influence methylation of CpG sites, associated to abundance of *Coriobacteriaceae* but we found no evidence for association between these SNPs and abundance of*Coriobacteriaceae* itself.

We did not find any genetic loci that explain the associations of microbial taxa to CpGs, but we could show that exposome factors, mainly diet and medication factors, might drive 7 of 82 (8.5%) CpG–microbiome associations. Notably, both the abundance of a genus-level cluster of the *Coriobacteriaceae* family and the methylation at cg13058819 correlated with increased coffee consumption (P = 8.93 x 10^-25^ and P = 1.42 x 10^-02^, respectively), with *Coriobacteriaceae* family abundance mediating 75% of the effect of coffee consumption on cg13058819. In earlier meQTL studies, the genetic determinant of this CpG was related to diverticular disease.

Overall, we discovered and confidently replicated two host methylation loci related to genus *Eggerthella* and identified mediating effects of gut bacteria on host gene methylation. While our research remains underpowered due to relatively small samples size and heterogeneity of microbiome and diet, these two cohorts currently form the largest dataset of simultaneous studies of microbiome, epigenetics and environment. Our results demonstrate the importance of studying microbiota and epigenetic variations concurrently when exploring the effects of diet and medication on host health.

## Acknowledgements

AZ is supported by the Netherlands Organization for Scientific Research (NWO)-VIDI grant 016.178.056, Netherlands Heart Foundation CVON grant 2018-27. AK and AZ are supported by the NWO Gravitation grant ExposomeNL 024.004.017. We thank Kateryna Onistrat for help with figure preparation.

## Data availability statement

The raw sequence data for both MGS and 16*S* rRNA gene sequencing data sets, and age and gender information per sample are available from the European genome-phenome archive (https://www.ebi.ac.uk/ega/) at accession number EGAS00001001704. Other phenotypic data can be requested from the LifeLines cohort study (https://lifelines.nl/lifelines-research/access-to-lifelines) following the standard protocol for data access. All data access to the Lifelines population cohort must follow the informed consent regulations of the Medical Ethics Review Board of the University Medical Center Groningen, which are clearly described at https://lifelines.nl/upload/file/lifelines+data+access+policy_%5B1%5D.pdf.

The pipeline for DNA methylation-array analysis developed by the Biobank-based Integrative Omics Study (BIOS) consortium are available here: https://molepi.github.io/DNAmArray_workflow/ (https://doi.org/10.5281/zenodo.3355292). The HumanMethylation450 BeadChip data from the LLD and NTR are available as part of the Biobank-based Integrative Omics Studies (BIOS) Consortium in the European Genome-phenome Archive (EGA), under the accession code EGAD00010000887, https://ega-archive.org/datasets/EGAD00010000887. The OMICs data and additional phenotype data are available upon request via the BBMRI-NL BIOS consortium (https://www.bbmri.nl/acquisition-use-analyze/bios). All NTR data can be requested by bona fida researchers (https://ntr-data-request.psy.vu.nl/).

Summary statistics of microbiome EWAS form LLD on 16s taxa, shotgun metagenomics derived taxa and pathways are deposited on https://doi.org/10.5281/zenodo.10062077

## Ethics statement

The Lifelines protocol was approved by the UMCG Medical ethical committee under number 2007/152. The Netherlands Twin Register: The study was approved by the Central Ethics Committee on Research Involving Human Subjects of the VU University Medical Centre, Amsterdam, an Institutional Review Board certified by the U.S. Office of Human Research Protections (IRB number IRB00002991 under Federal-wide Assurance-FWA00017598; IRB/institute codes, NTR 03-180).

## Author contributions

