## Supplementary figures and images for "Linking the gut microbiome to host DNA methylation by a discovery and replication epigenome-wide association study"

### Supplemental Figure 1

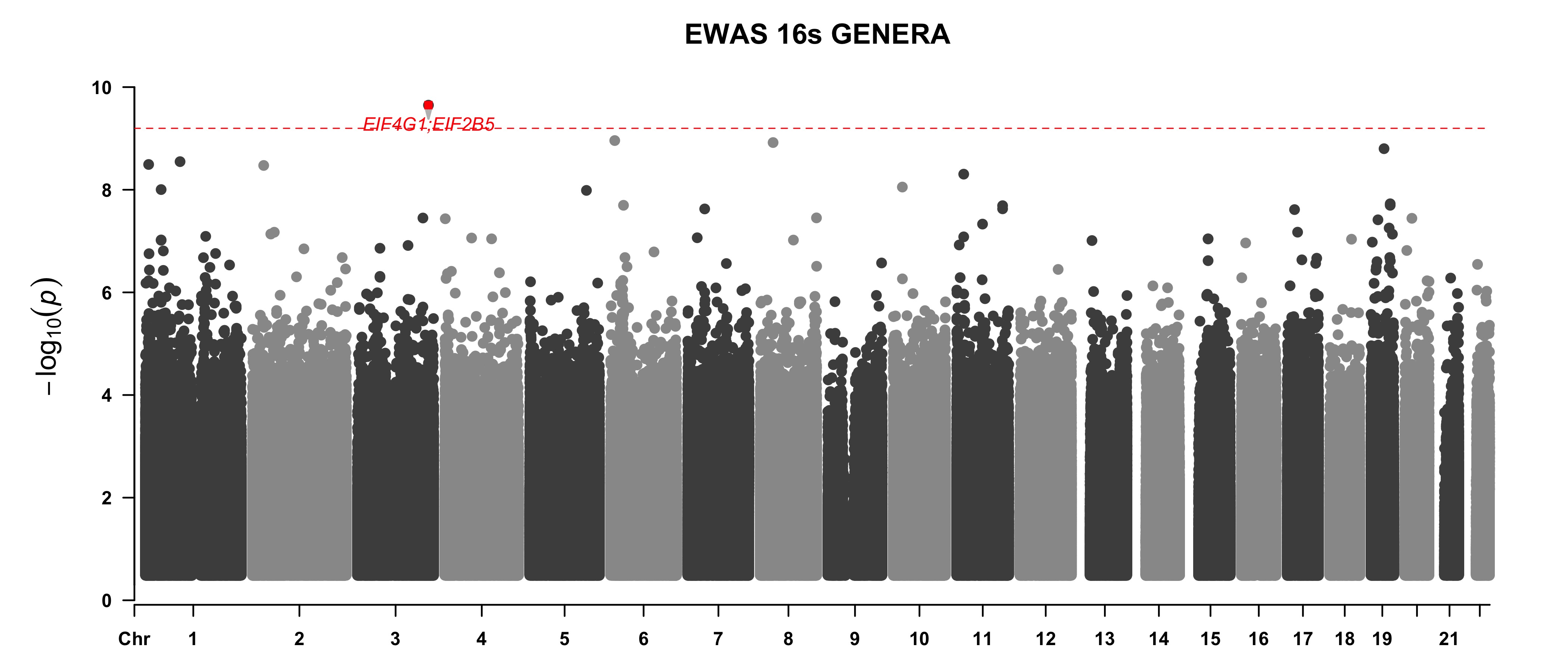

### Supplemental Figure 3

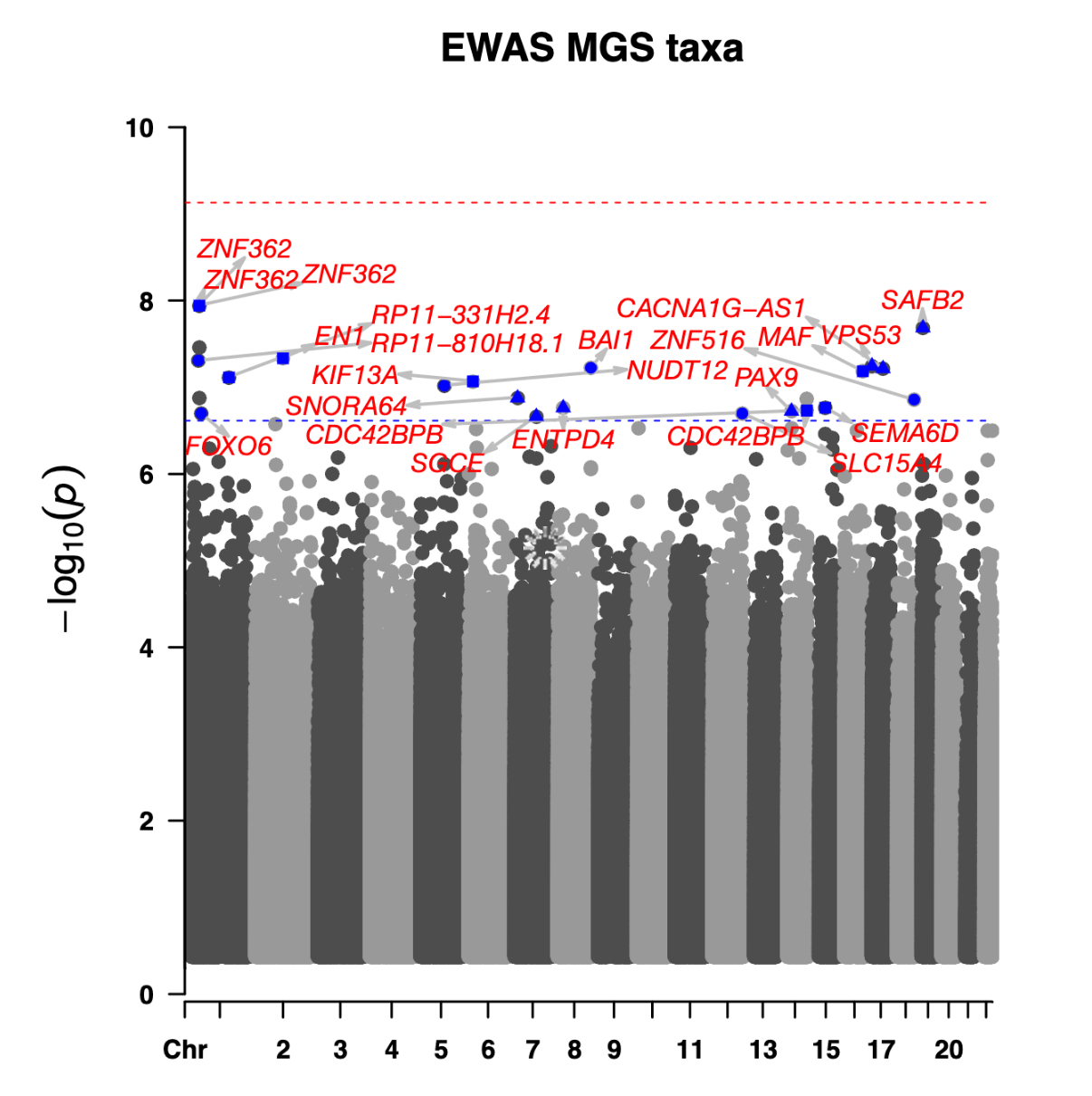

### Supplemental Figure 4

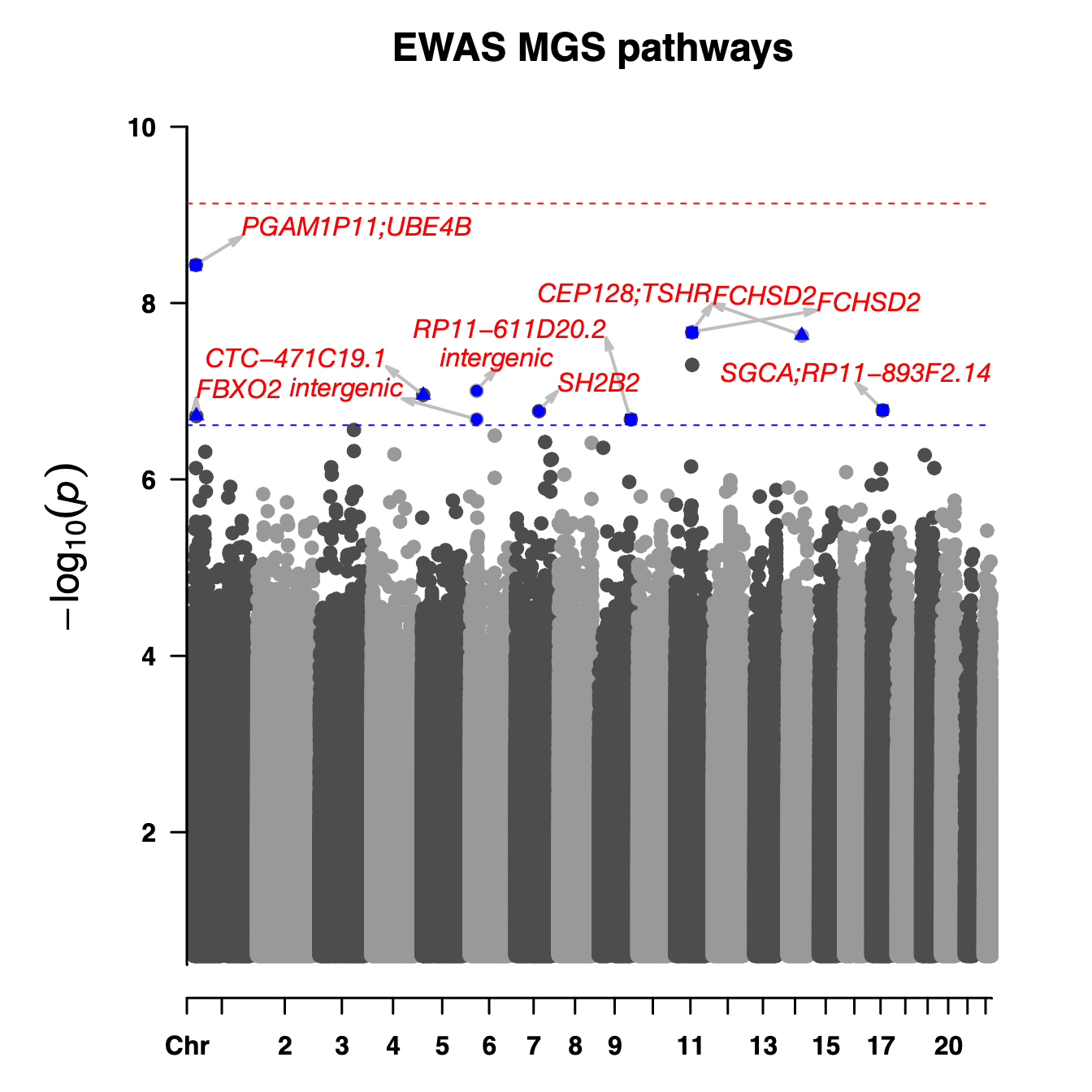
