## Supplemental Figure 2 for "Linking the gut microbiome to host DNA methylation by a discovery and replication epigenome-wide association study"

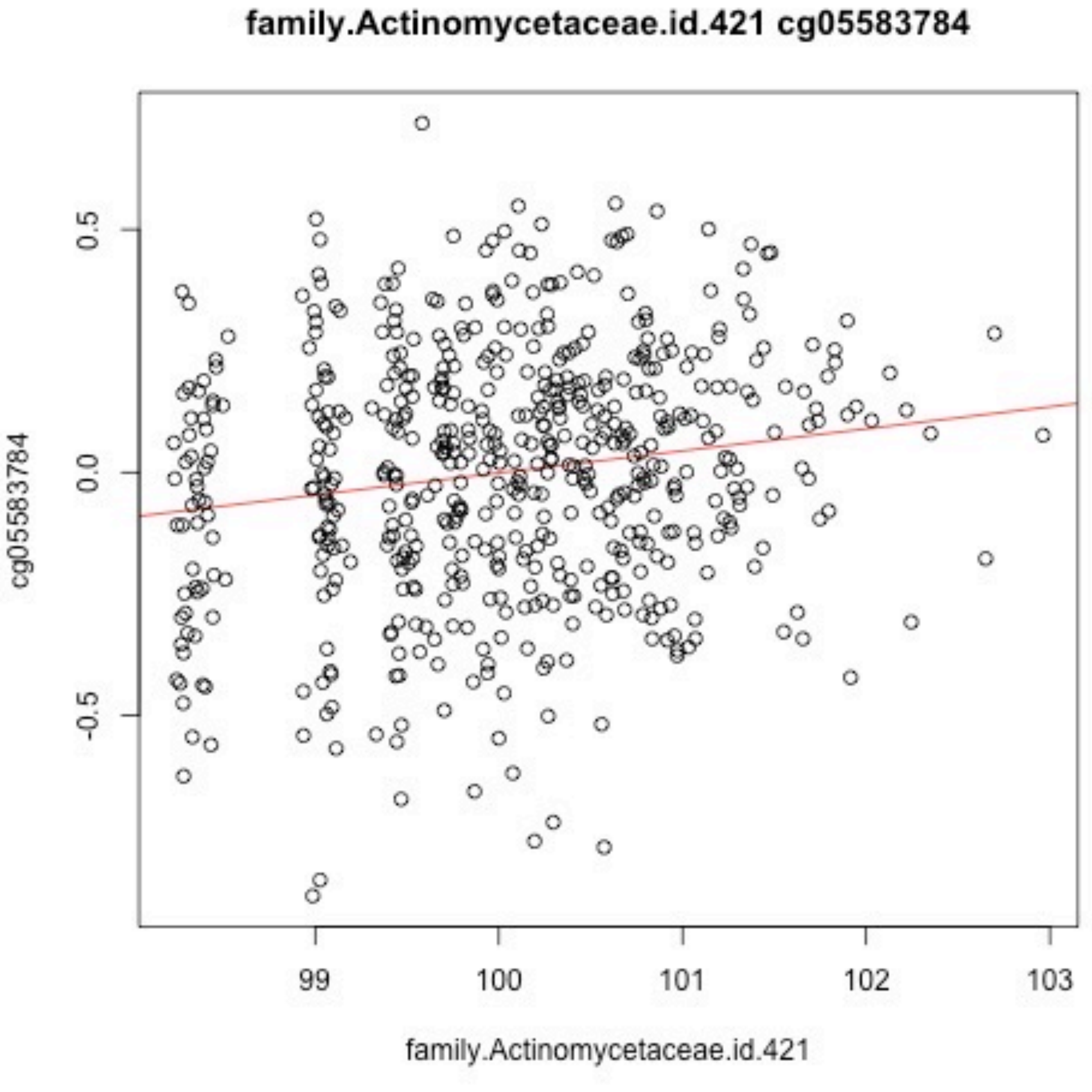
family.Actinomycetaceae.id.421 cg05583784
cg05583784
0.5
0.0
-0.5
99
100
101
102
103
family.Actinomycetaceae.id.421

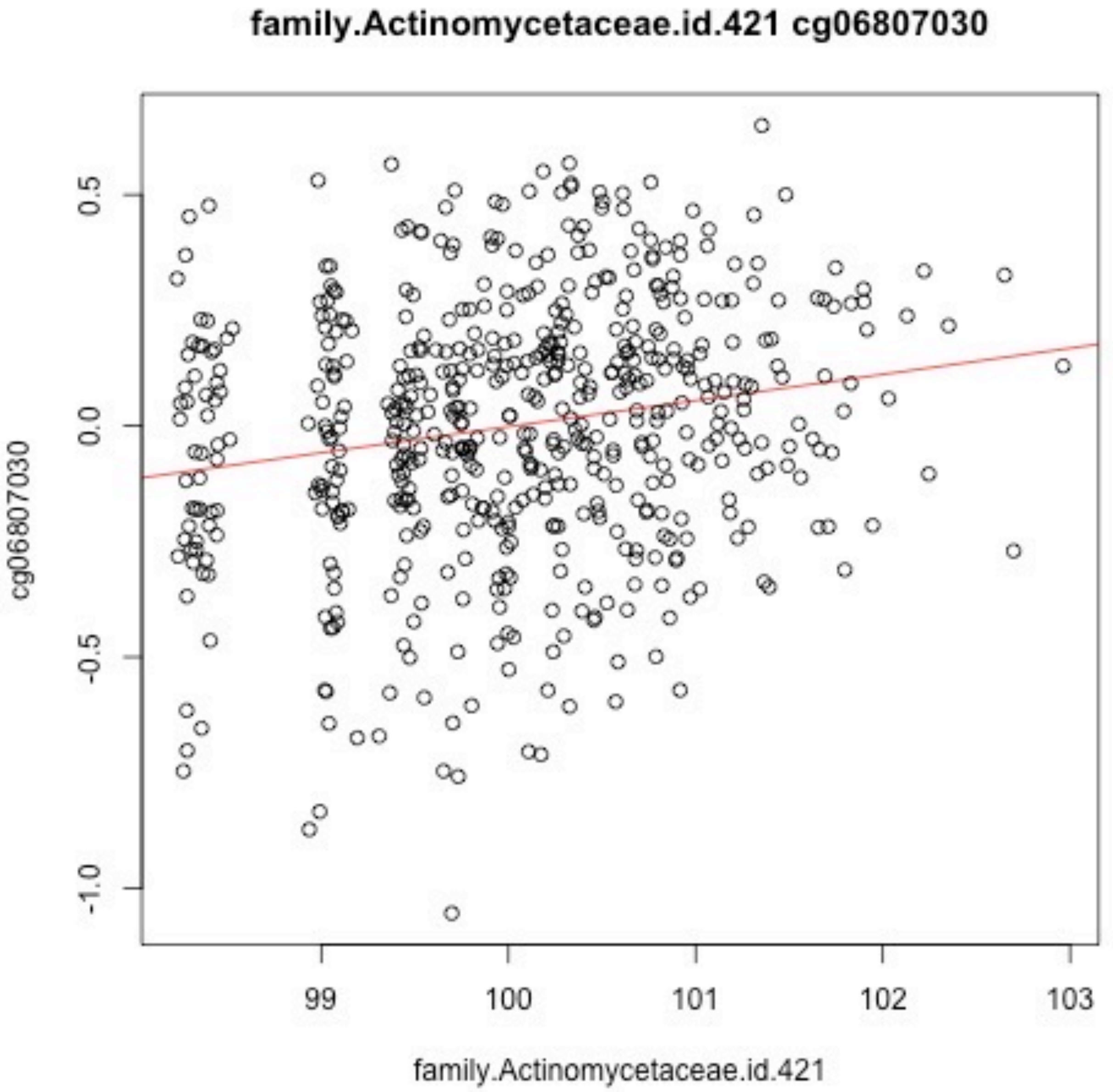
family.Actinomycetaceae.id.421 cg06807030
0.5
0.0
-0.5
-1.0
cg06807030
99
100
101
102
103
family.Actinomycetaceae.id.421

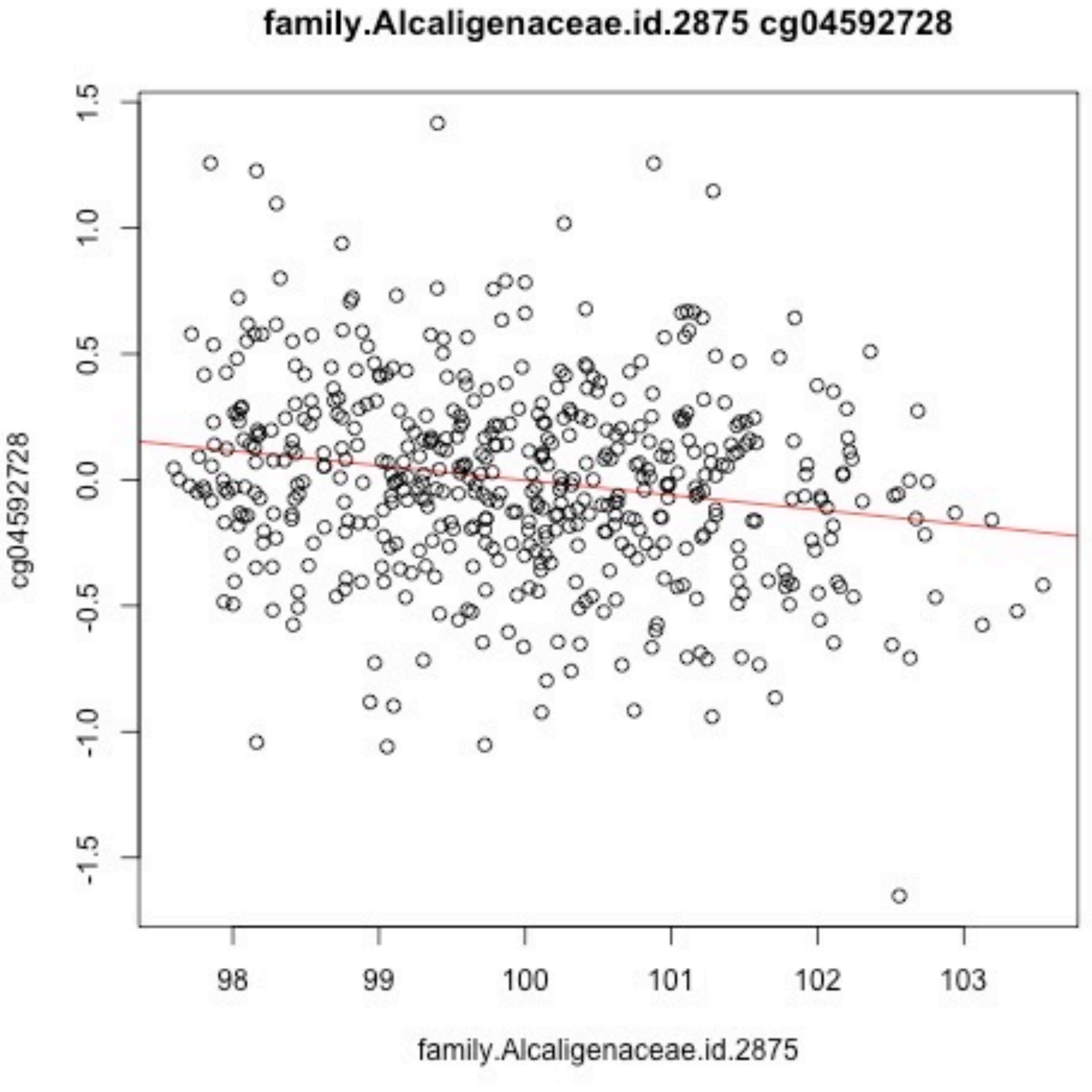
family.Alcaligenaceae.id.2875 cg04592728
cg04592728
1.5
1.0
0.5
0.0
-0.5
-1.0
-1.5
98
99
100
101
102
103
family.Alcaligenaceae.id.2875

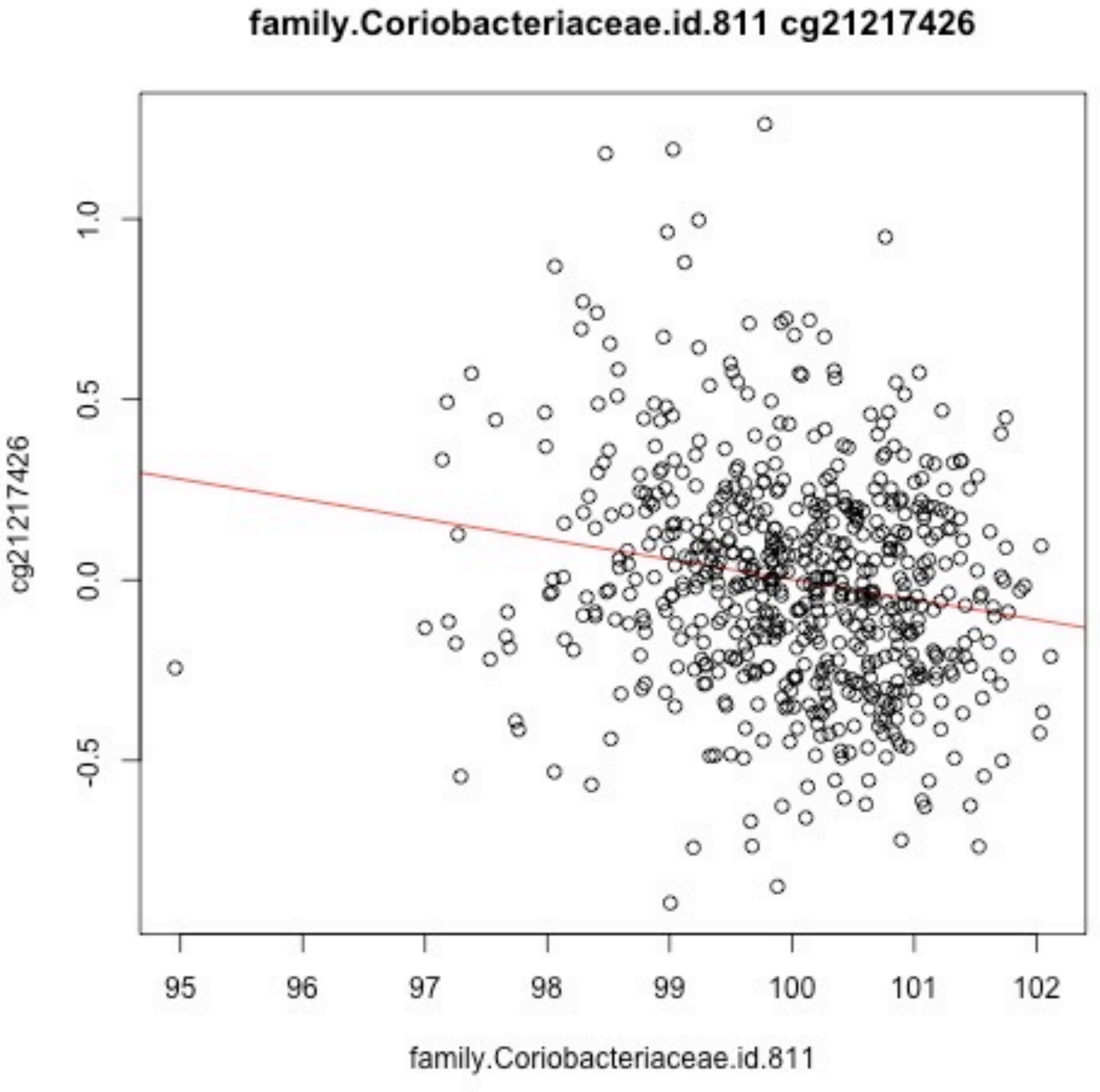
family.Coriobacteriaceae.id.811 cg21217426
cg21217426
1.0
0.5
0.0
-0.5
95
96
97
98
99
100
101
102
family.Coriobacteriaceae.id.811

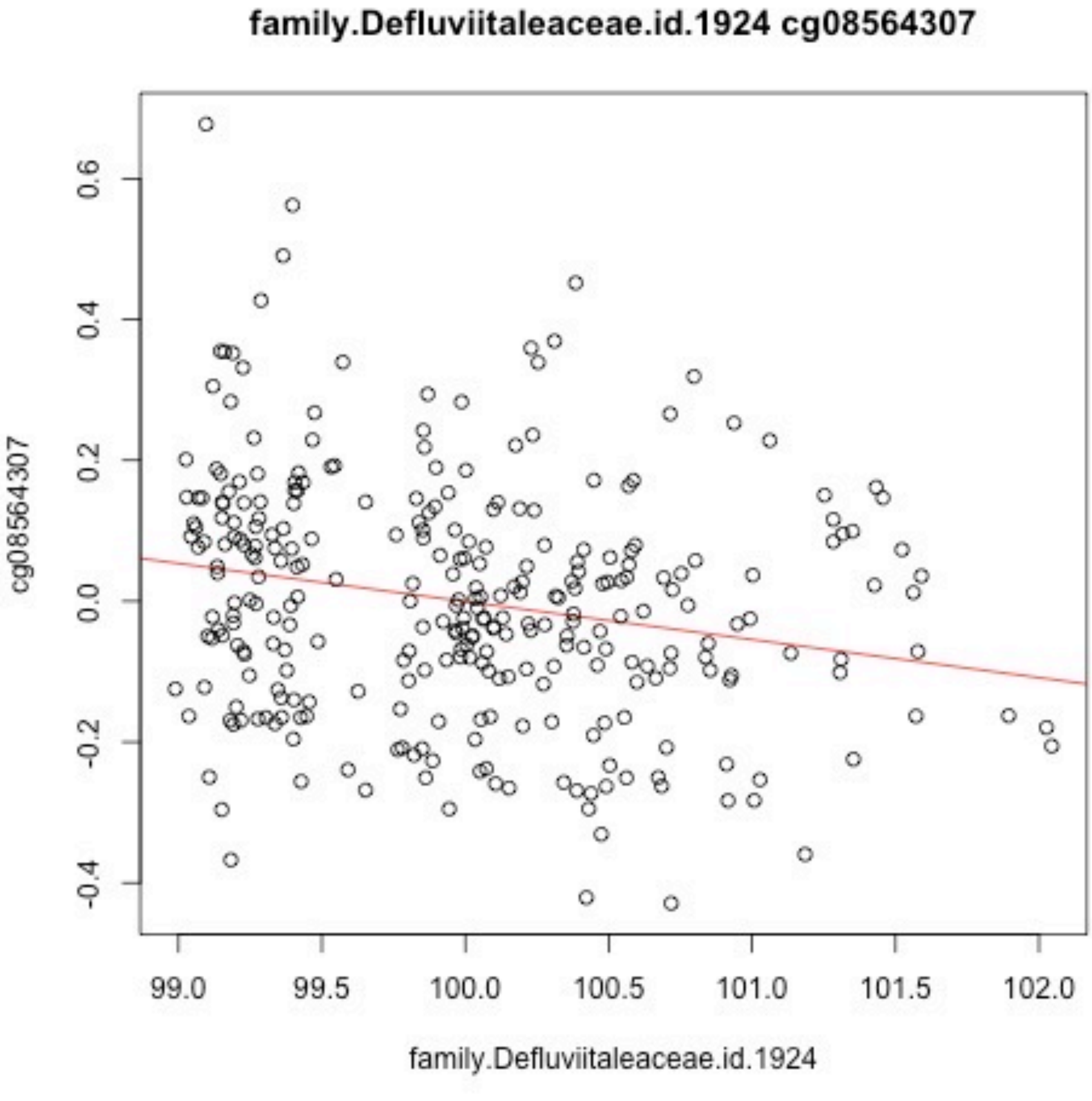
family.Defluviitaleaceae.id.1924 cg08564307
cg08564307
0.6
0.4
0.2
0.0
-0.2
-0.4
99.0
99.5
100.0
100.5
101.0
101.5
102.0
family.Defluviitaleaceae.id.1924

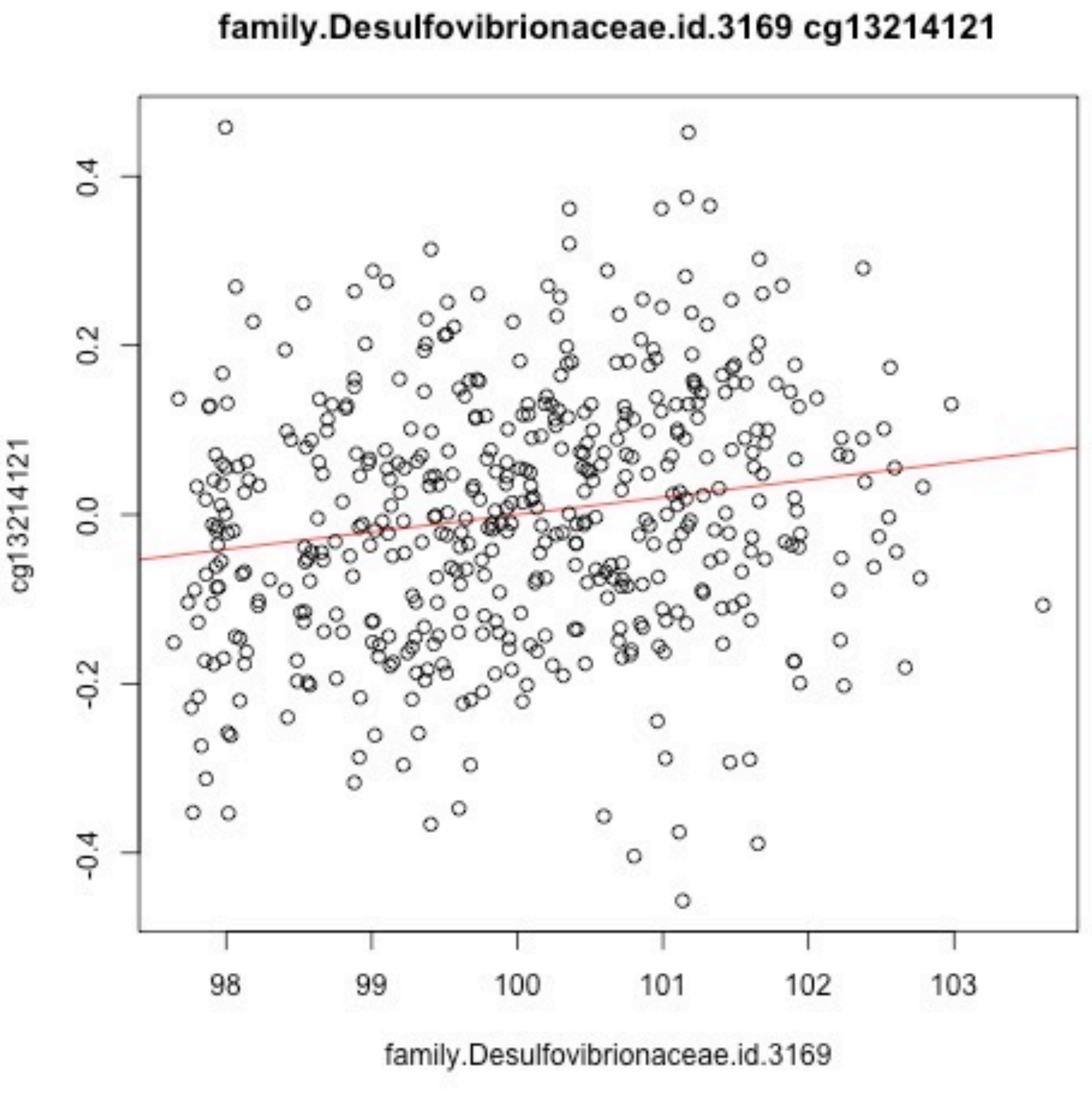
family.Desulfovibrionaceae.id.3169 cg13214121
cg13214121
0.4
0.2
0.0
-0.2
-0.4
98
99
100
101
102
103
family.Desulfovibrionaceae.id.3169

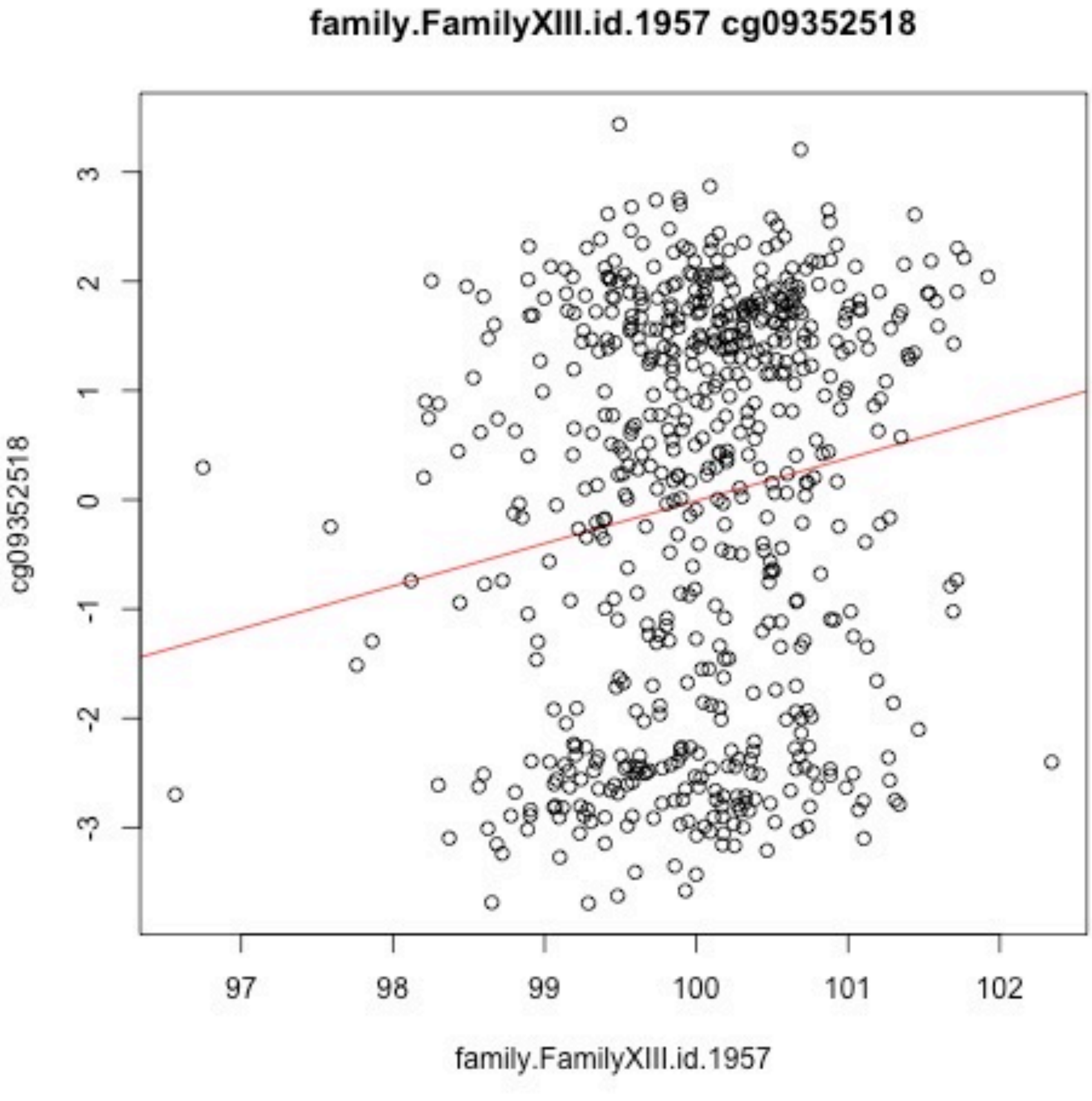
family.FamilyXIII.id.1957 cg09352518
cg09352518
3
2
1
0
-1
-2
-3
97
98
99
100
101
102
family.FamilyXIII.id.1957

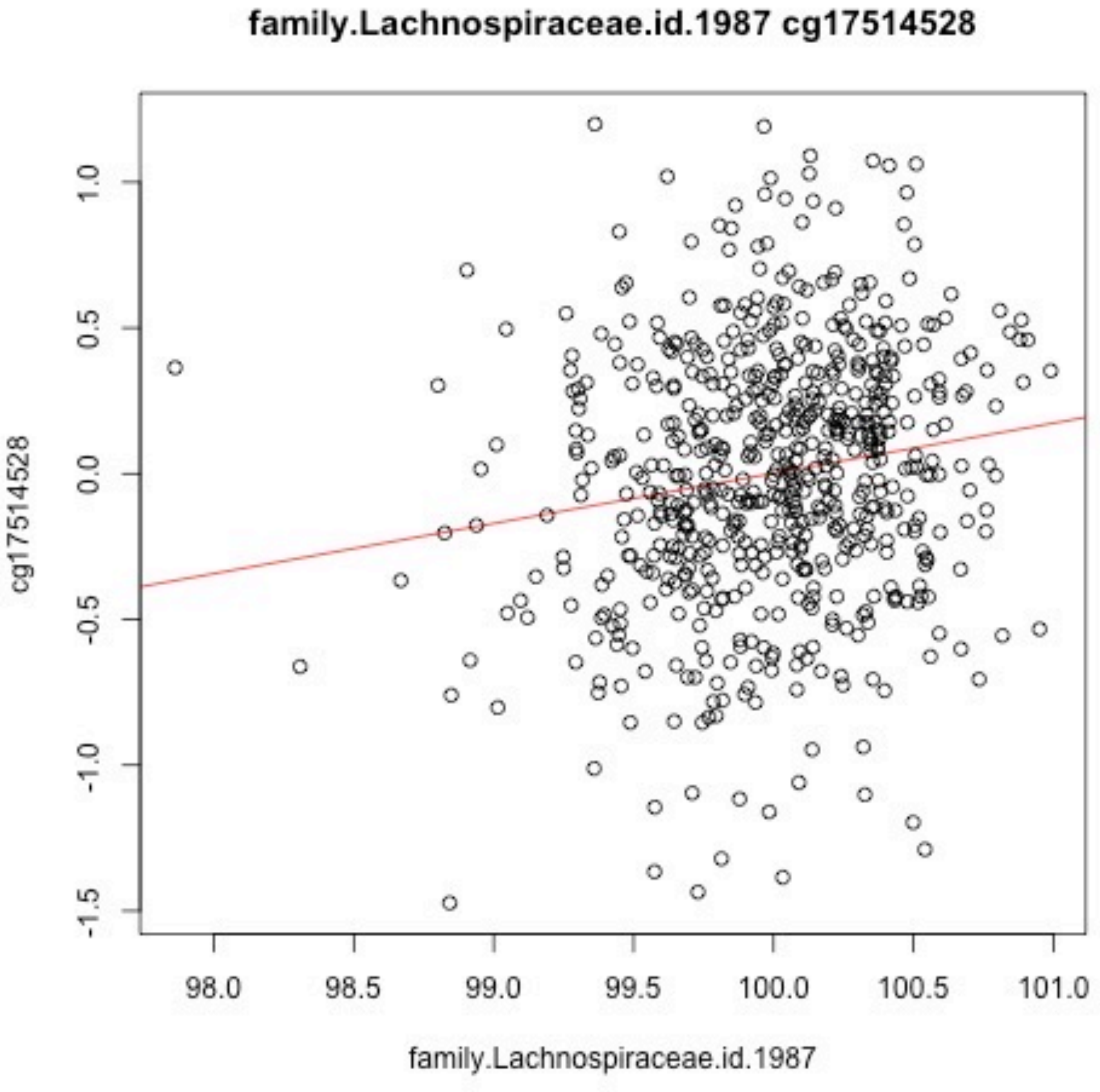
family.Lachnospiraceae.id.1987 cg17514528
cg17514528
family.Lachnospiraceae.id.1987
1.0
0.5
0.0
-0.5
-1.0
-1.5
98.0
98.5
99.0
99.5
100.0
100.5
101.0

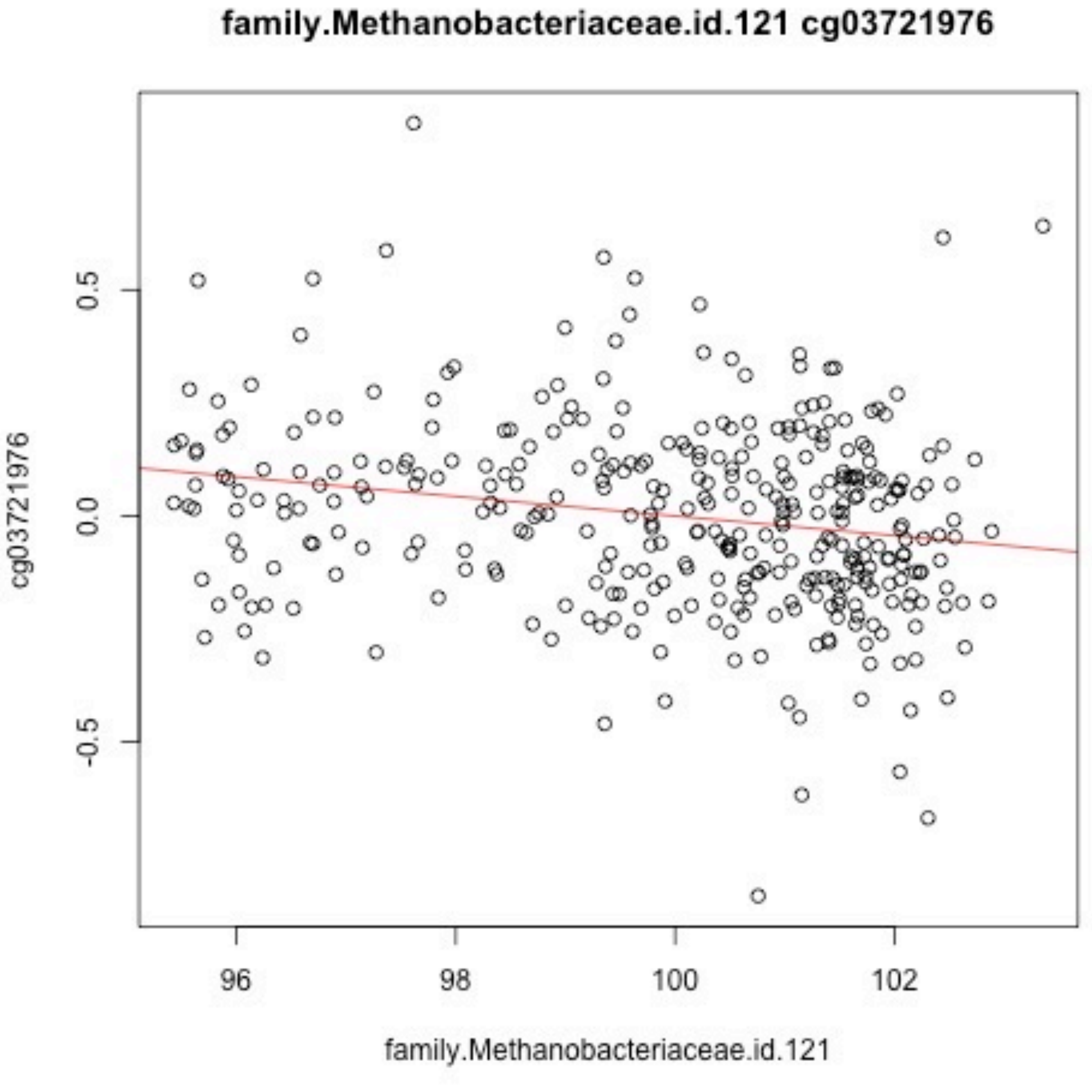
family.Methanobacteriaceae.id.121 cg03721976
0.5
0.0
-0.5
cg03721976
96
98
100
102
family.Methanobacteriaceae.id.121

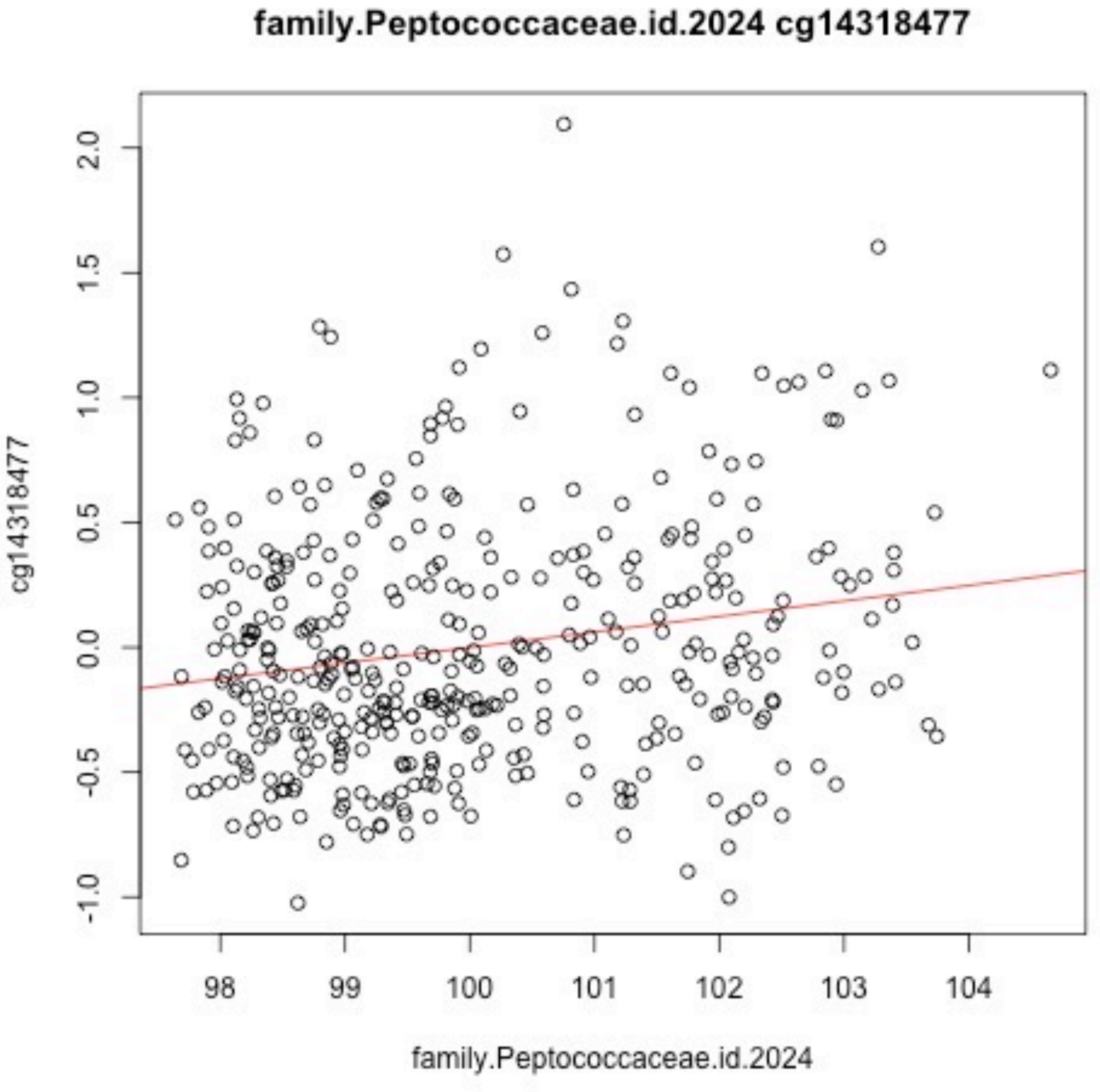
family.Peptococcaceae.id.2024 cg14318477
cg14318477
2.0
1.5
1.0
0.5
0.0
-0.5
-1.0
98
99
100
101
102
103
104
family.Peptococcaceae.id.2024

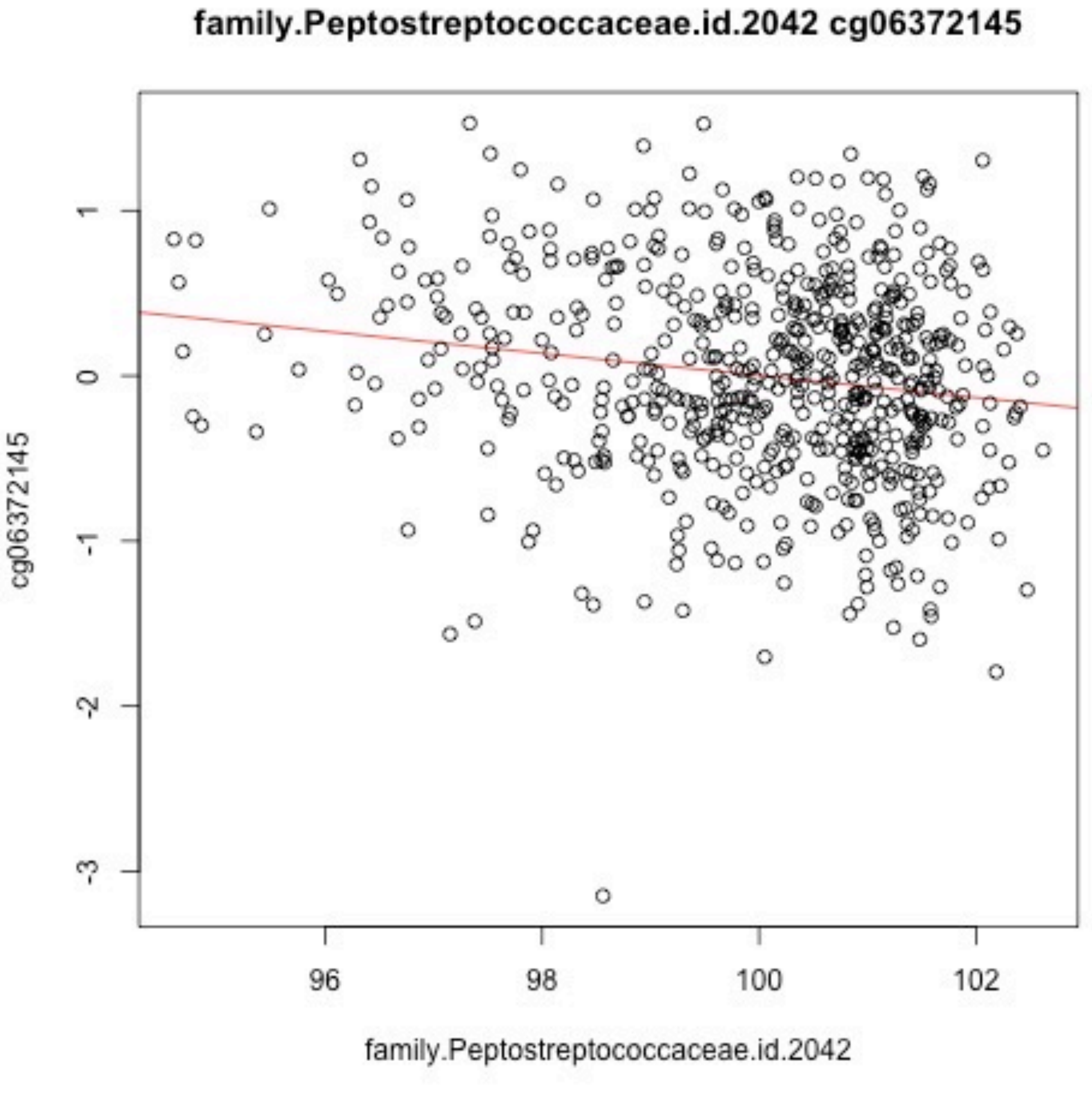

family.Peptostreptococcaceae.id.2042 cg06372145
cg06372145
1
0
-1
-2
-3
96
98
100
102
family.Peptostreptococcaceae.id.2042

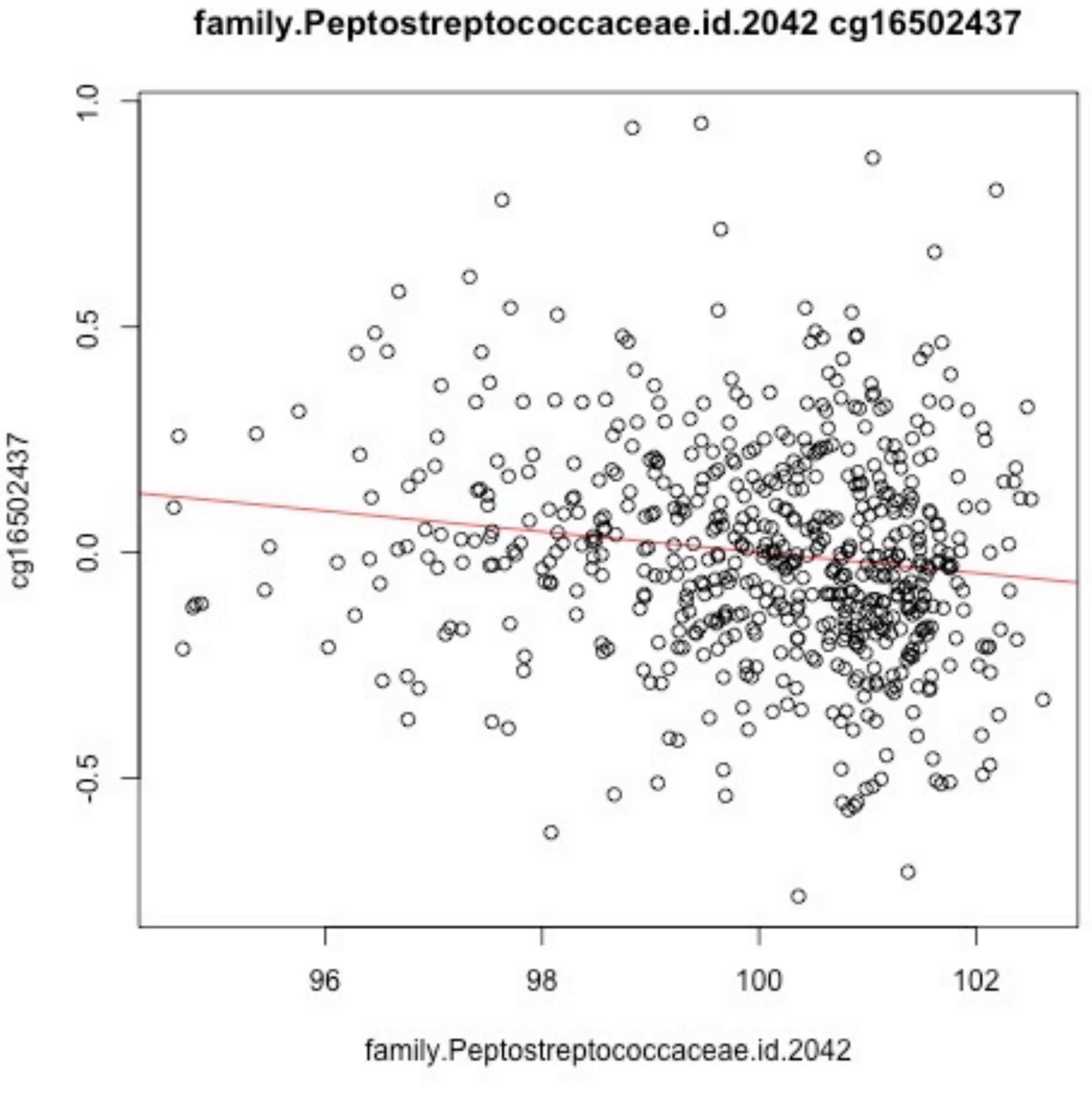

family.Peptostreptococcaceae.id.2042 cg16502437
1.0
0.5
0.0
-0.5
cg16502437
96
98
100
102
family.Peptostreptococcaceae.id.2042

family.Porphyromonadaceae.id.943 cg10542975

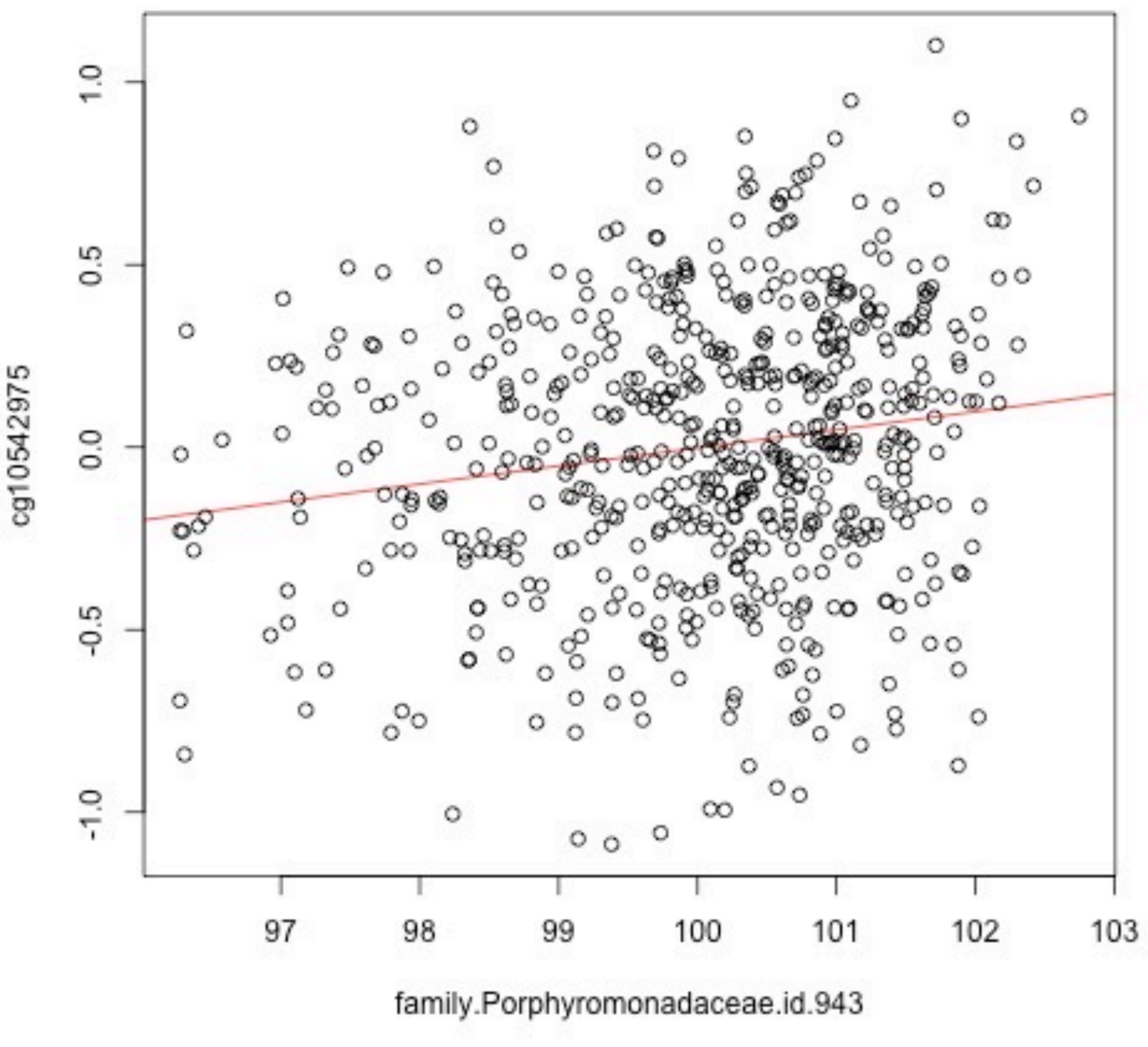

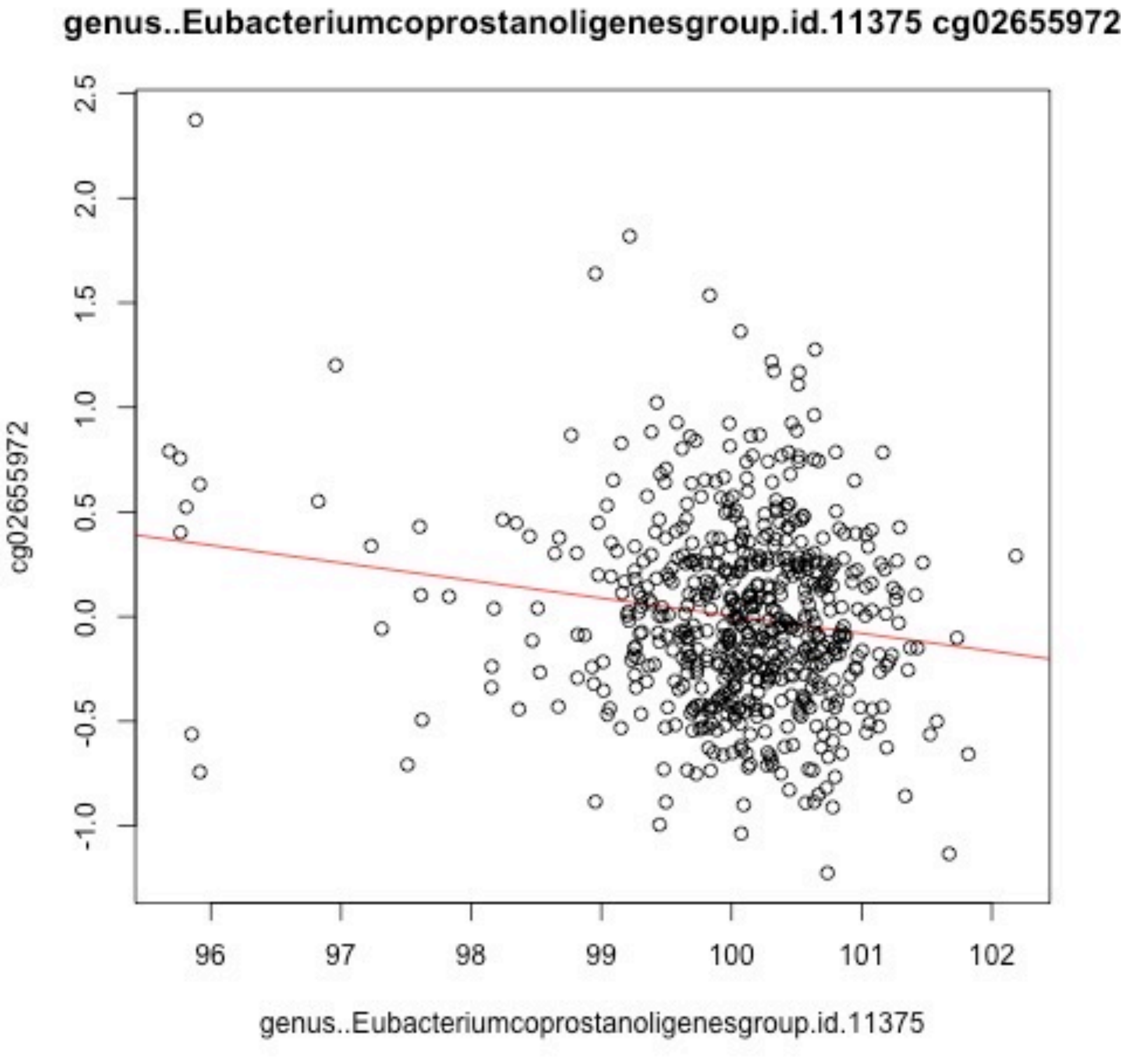
genus..Eubacteriumcoprostanoligenesgroup.id.11375 cg02655972
cg02655972
2.5
2.0
1.5
1.0
0.5
0.0
-0.5
-1.0
96
97
98
99
100
101
102
genus..Eubacteriumcoprostanoligenesgroup.id.11375

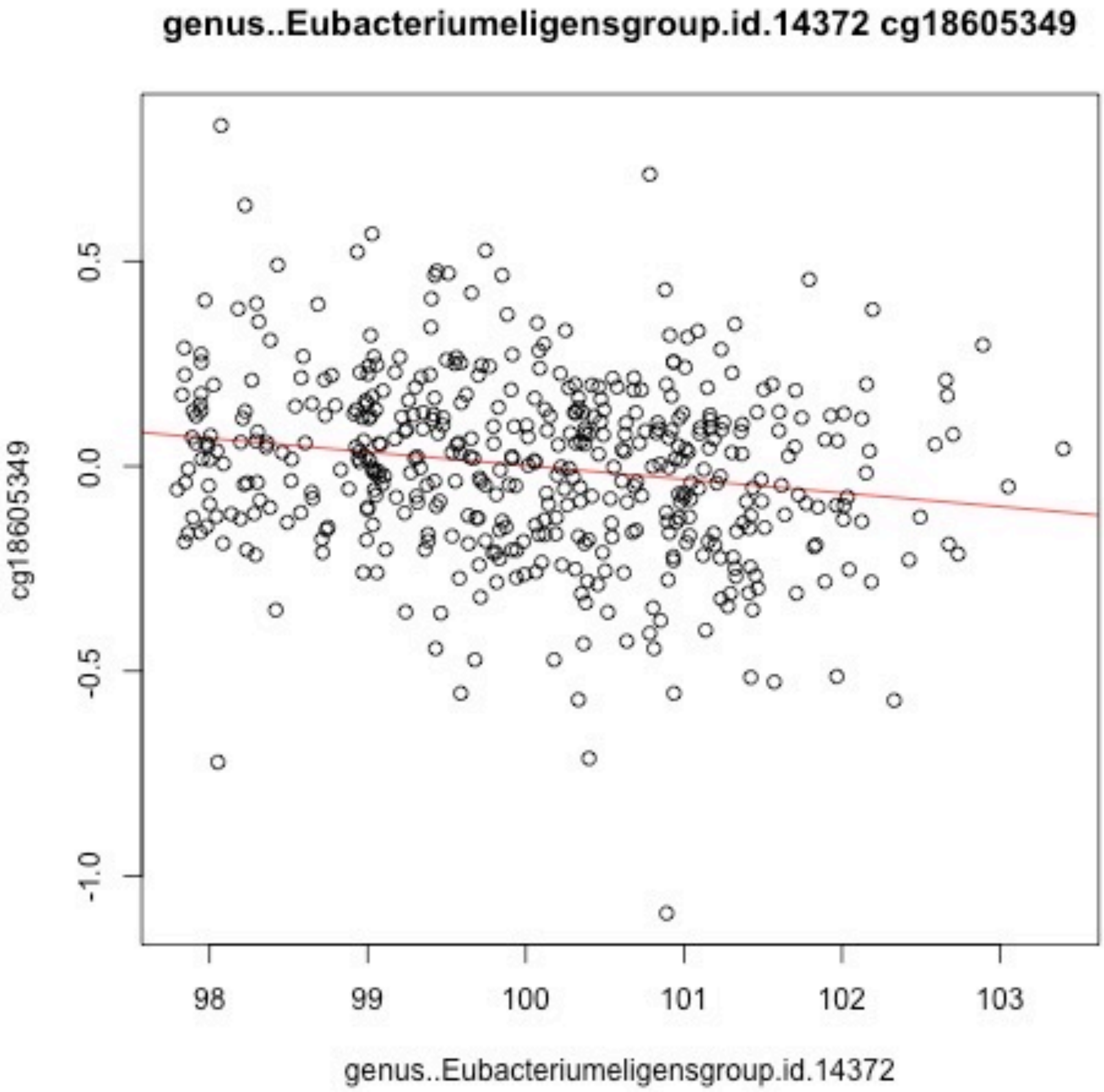
genus..Eubacteriumeligensgroup.id.14372 cg18605349
cg18605349
0.5
0.0
-0.5
-1.0
98
99
100
101
102
103
genus..Eubacteriumeligensgroup.id.14372

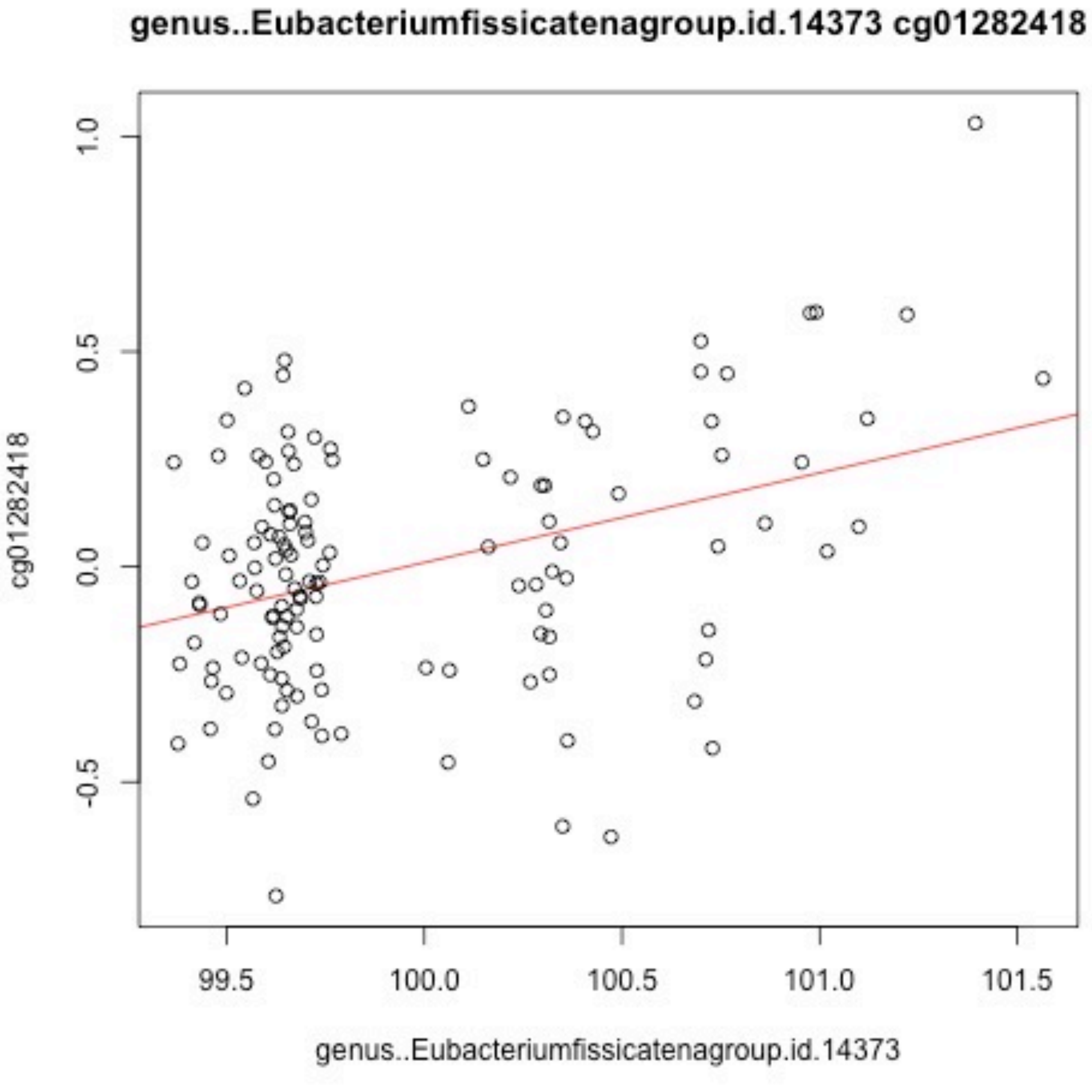
genus..Eubacteriumfissicatenagroup.id.14373 cg01282418
cg01282418
1.0
0.5
0.0
-0.5
99.5
100.0
100.5
101.0
101.5
genus..Eubacteriumfissicatenagroup.id.14373

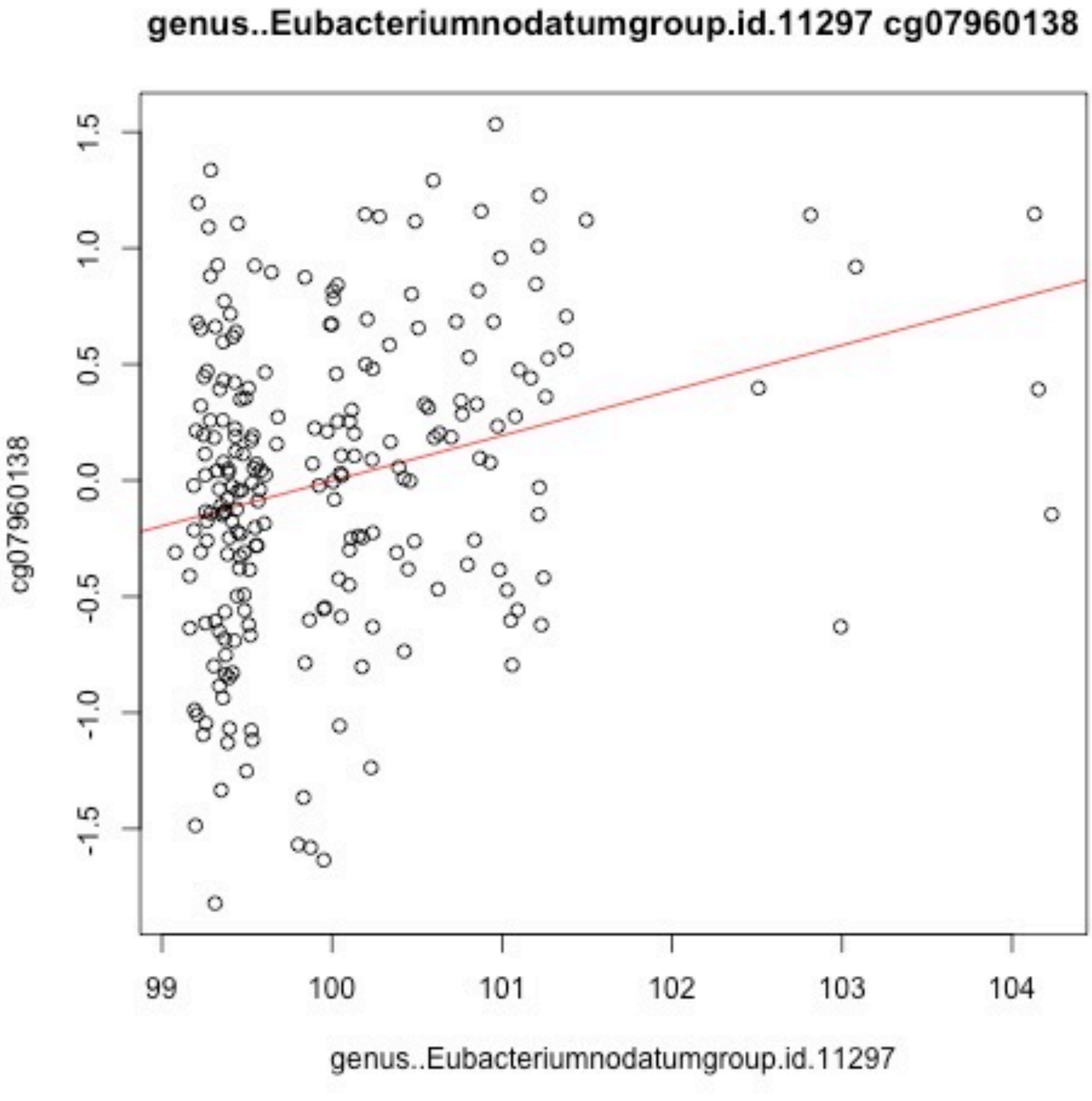

genus..Eubacteriumnodatumgroup.id.11297 cg07960138
cg07960138
1.5
1.0
0.5
0.0
-0.5
-1.0
-1.5
99
100
101
102
103
104
genus..Eubacteriumnodatumgroup.id.11297

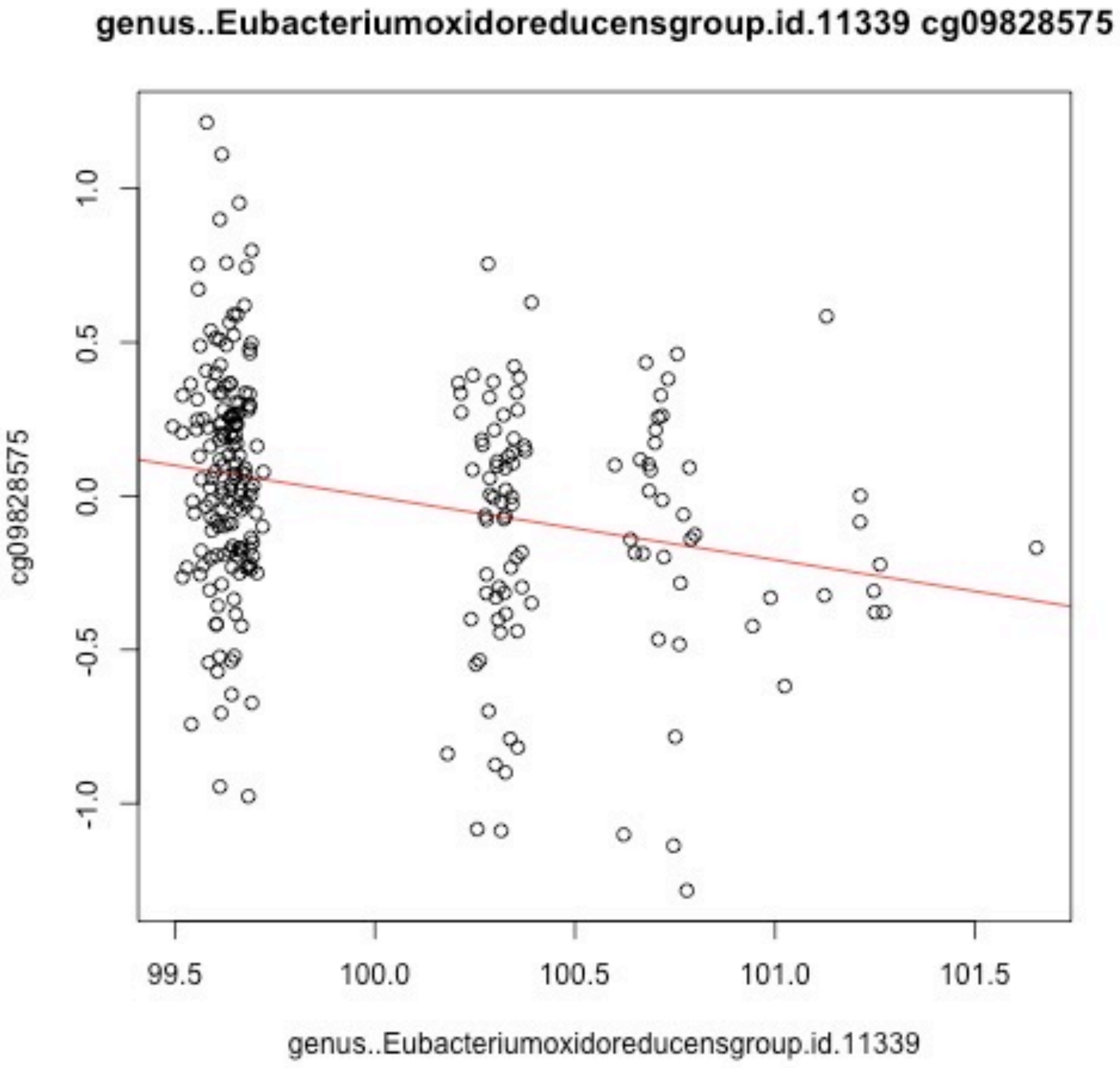
genus..Eubacteriumoxidoreducensgroup.id.11339 cg09828575
cg09828575
1.0
0.5
0.0
-0.5
-1.0
99.5
100.0
100.5
101.0
101.5
genus..Eubacteriumoxidoreducensgroup.id.11339

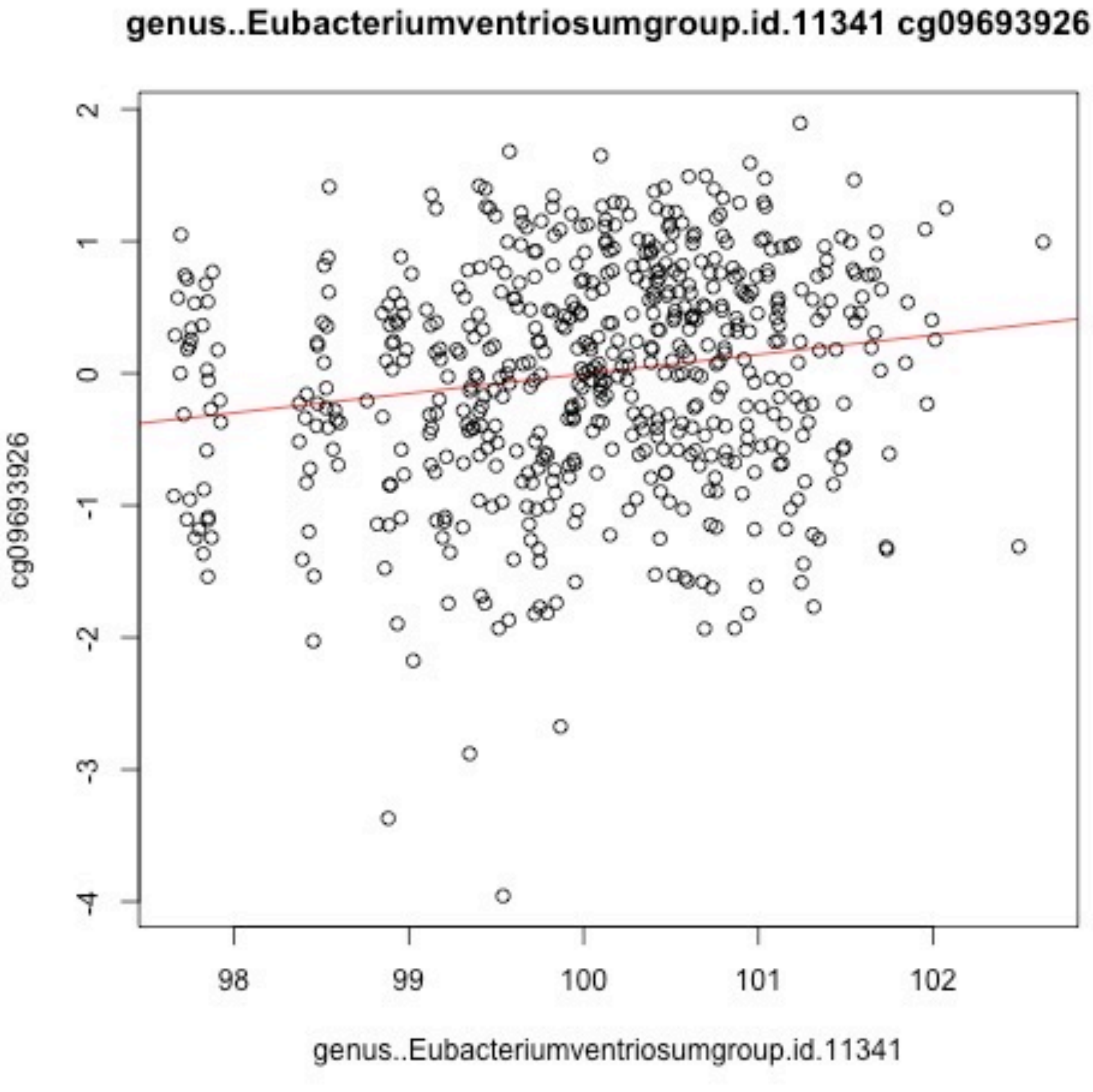

genus..Eubacteriumventriosumgroup.id.11341 cg09693926
cg09693926
genus..Eubacteriumventriosumgroup.id.11341
2
1
0
-1
-2
-3
-4
98
99
100
101
102

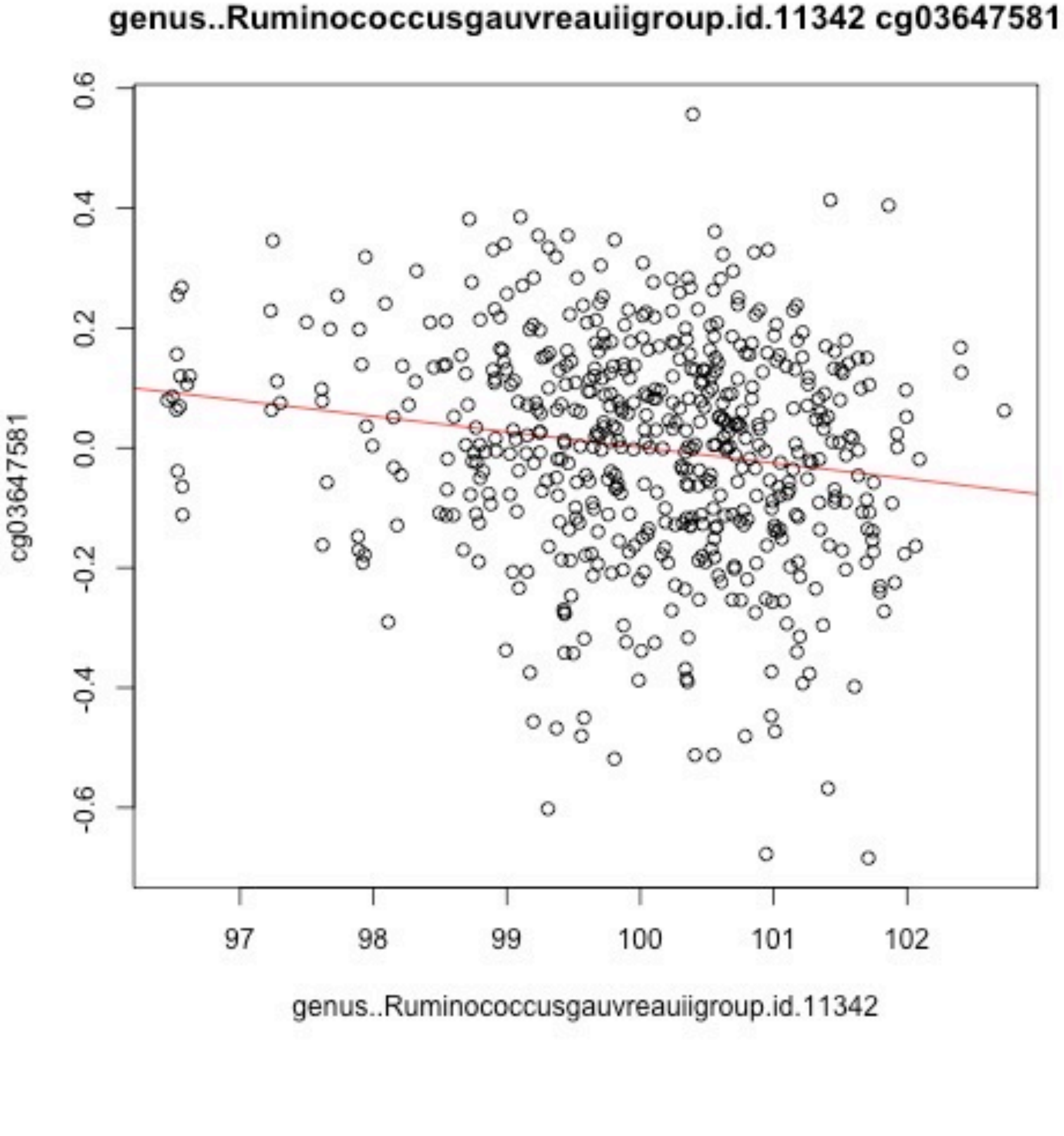

genus..Ruminococcusgauvreauiigroup.id.11342 cg03647581
0.6
0.4
0.2
0.0
-0.2
-0.4
-0.6
cg03647581
97
98
99
100
101
102
genus..Ruminococcusgauvreauiigroup.id.11342

genus..Ruminococcusgauvreauiigroup.id.11342 cg20400838

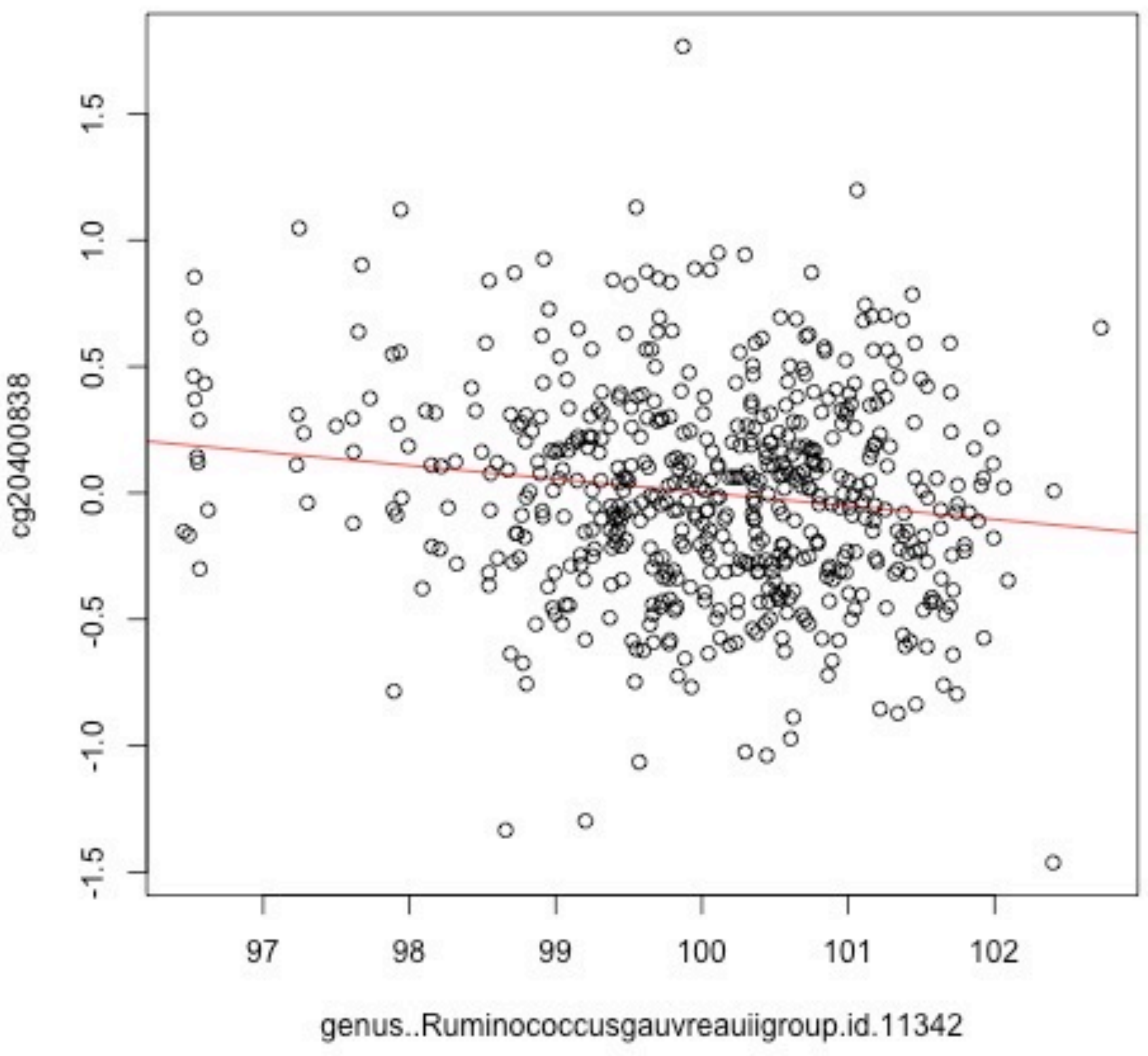

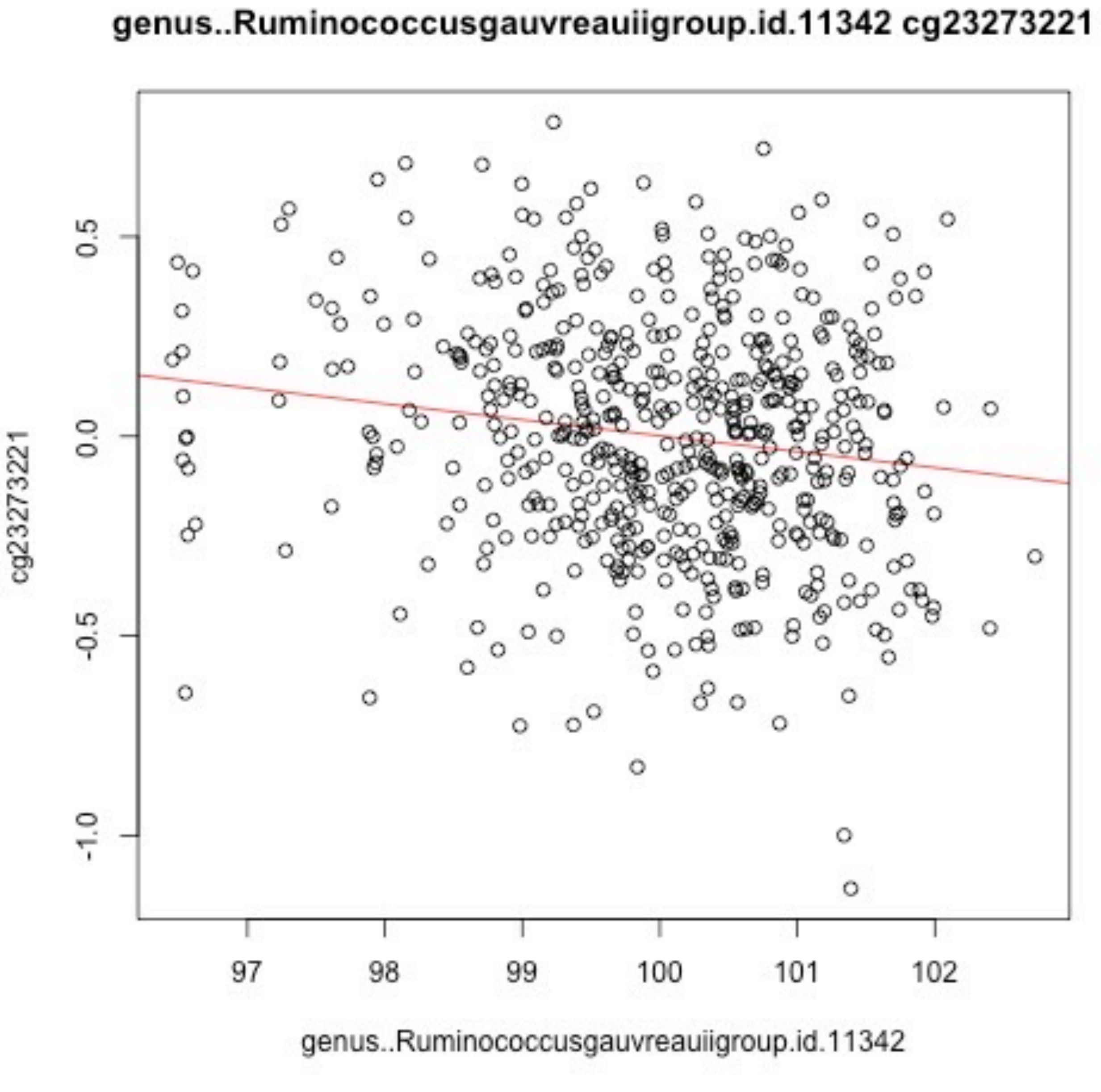

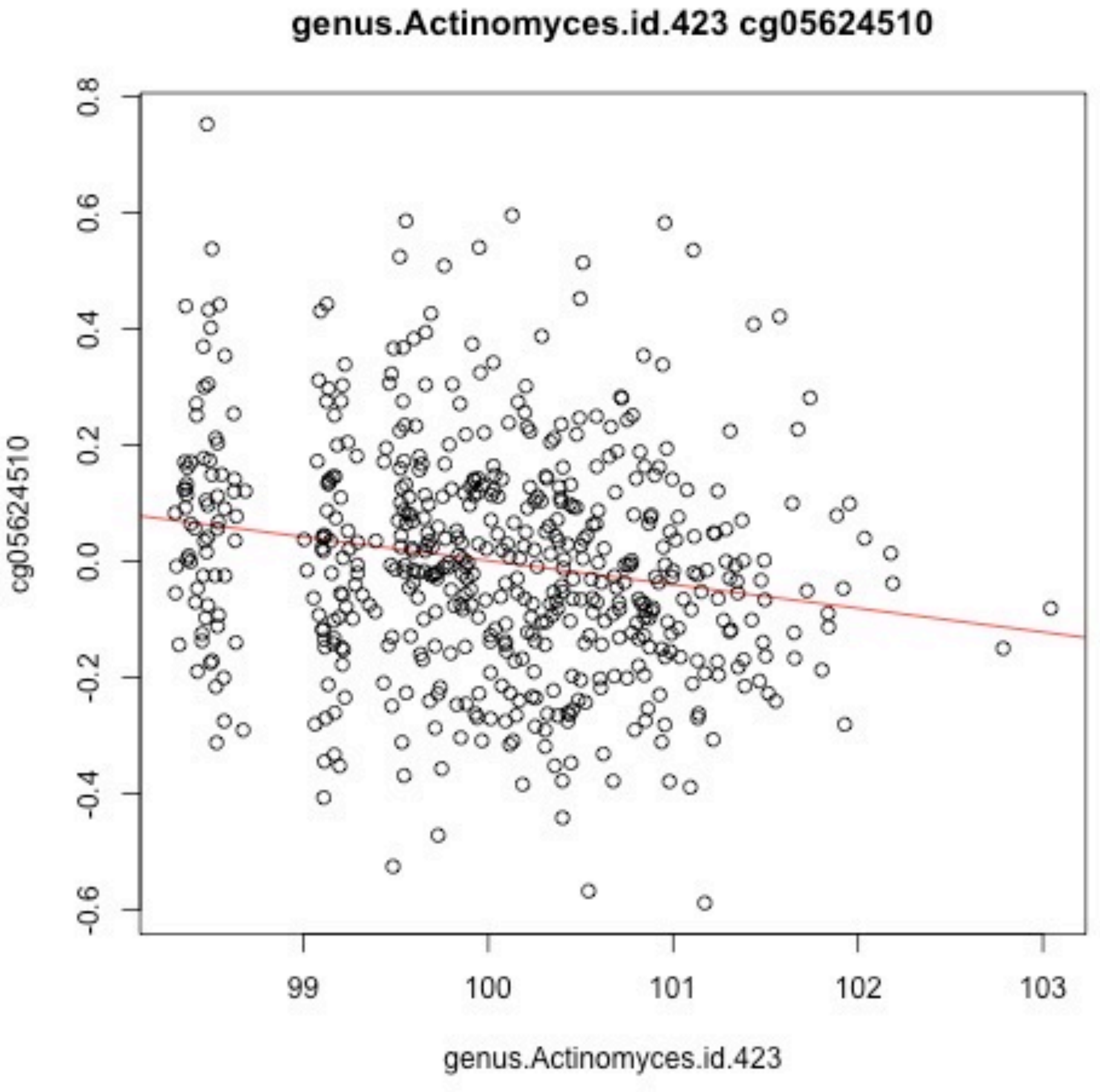
genus.Actinomyces.id.423 cg05624510
cg05624510
0.8
0.6
0.4
0.2
0.0
-0.2
-0.4
-0.6
99
100
101
102
103
genus.Actinomyces.id.423

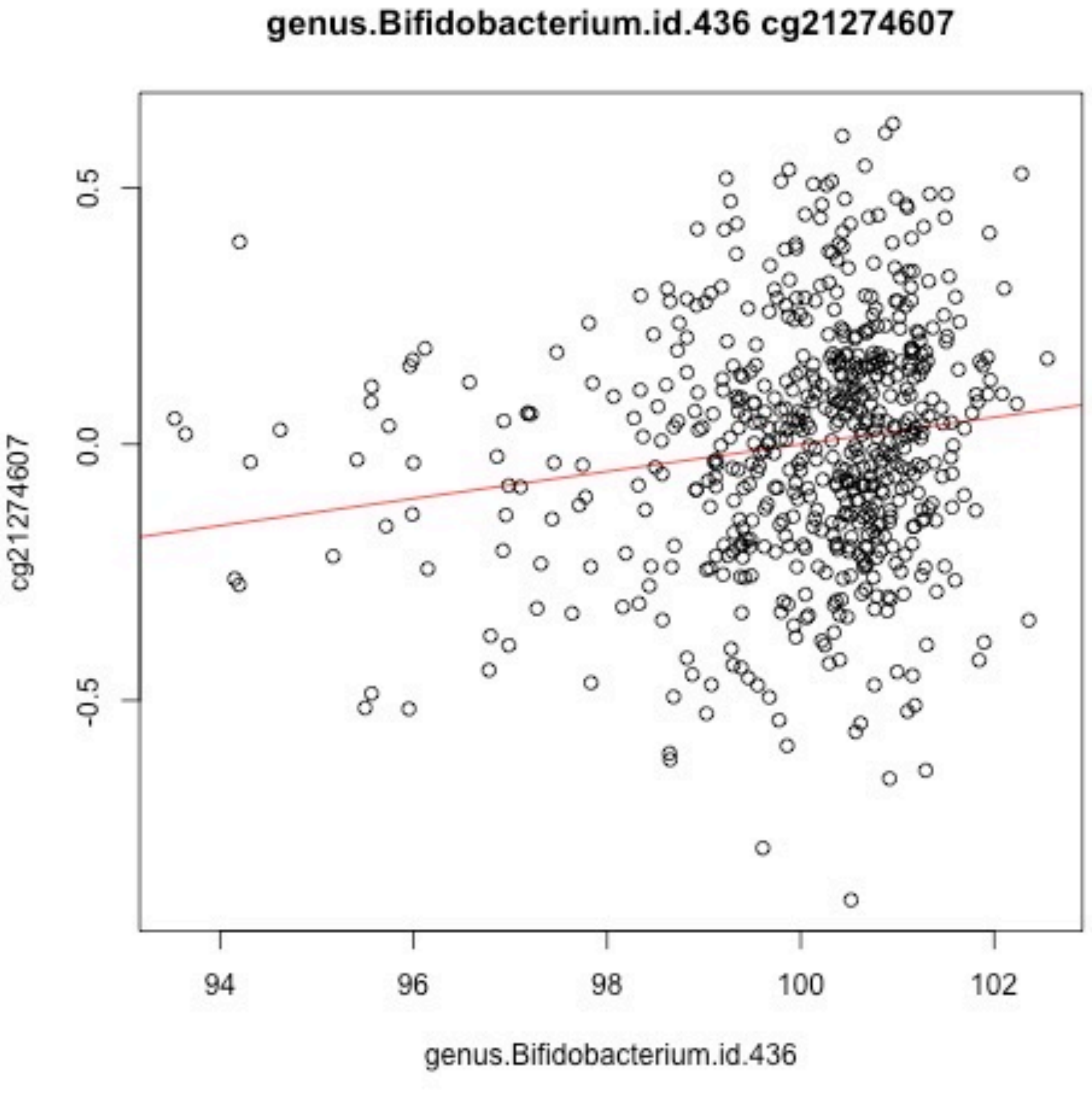
genus.Bifidobacterium.id.436 cg21274607
cg21274607
0.5
0.0
-0.5
94
96
98
100
102
genus.Bifidobacterium.id.436

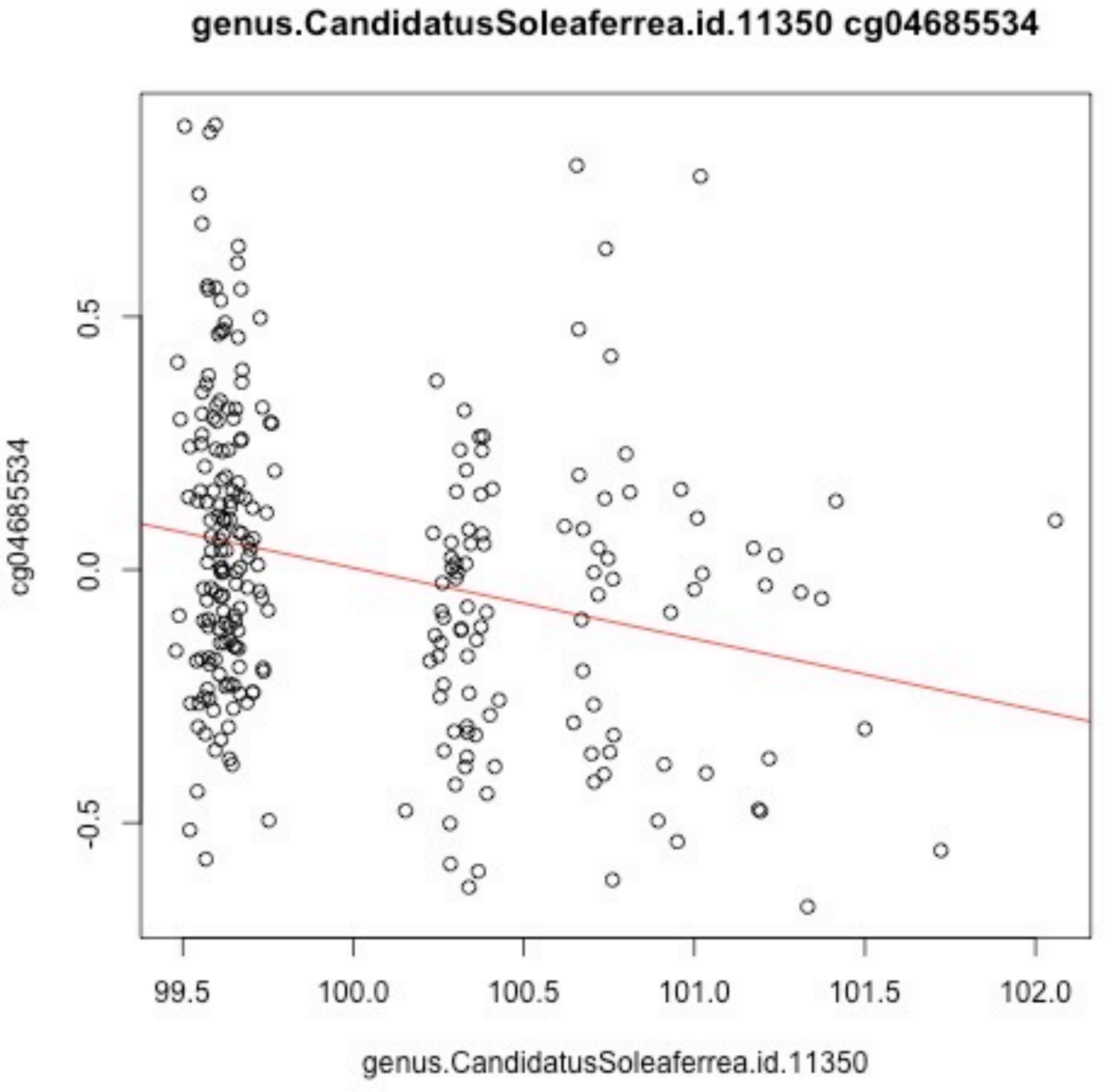
genus.CandidatusSoleaferrea.id.11350 cg04685534
cg04685534
0.5
0.0
-0.5
99.5
100.0
100.5
101.0
101.5
102.0
genus.CandidatusSoleaferrea.id.11350

genus.Clostridiumsensustricto1.id.1873 cg08238777

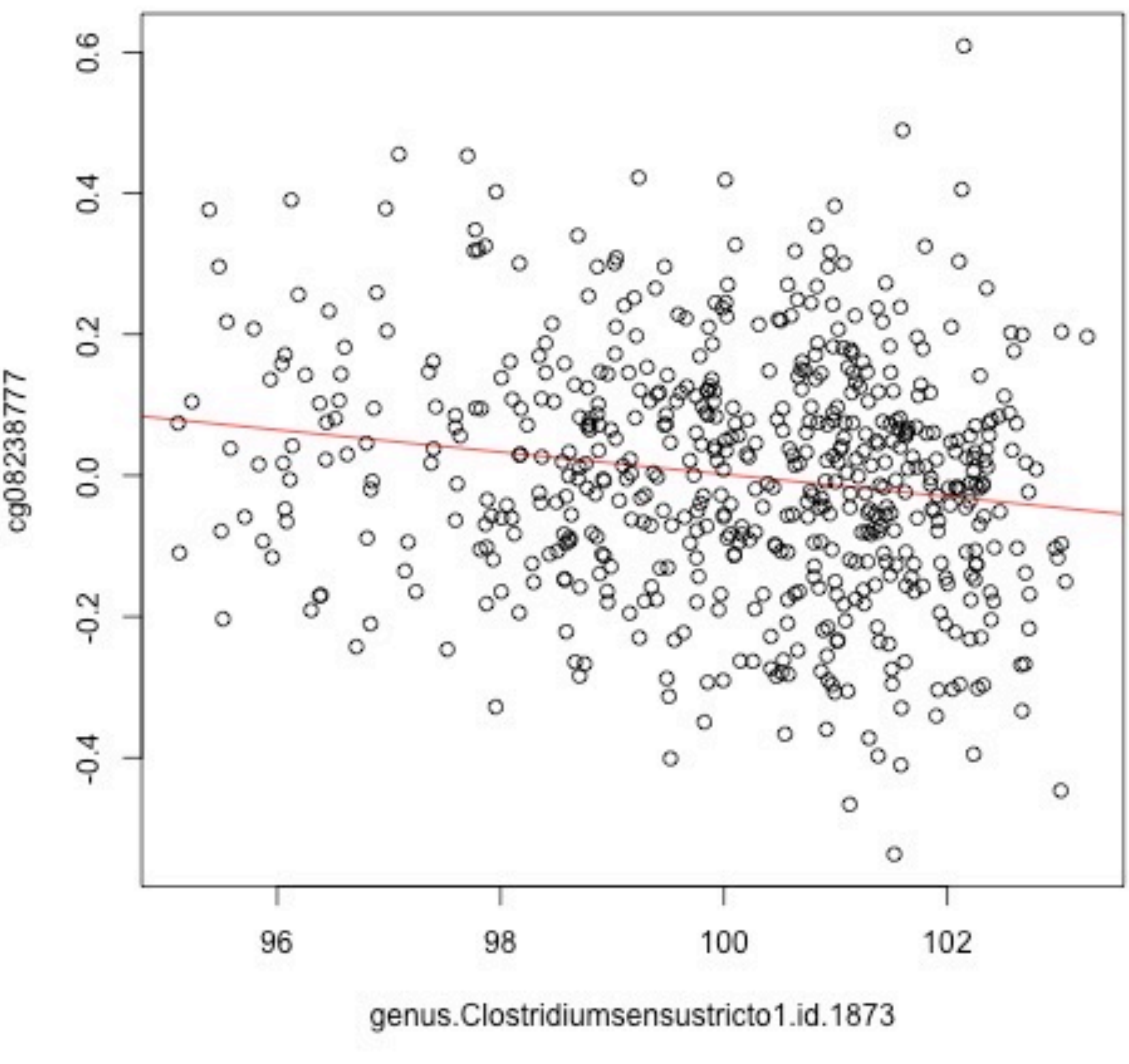

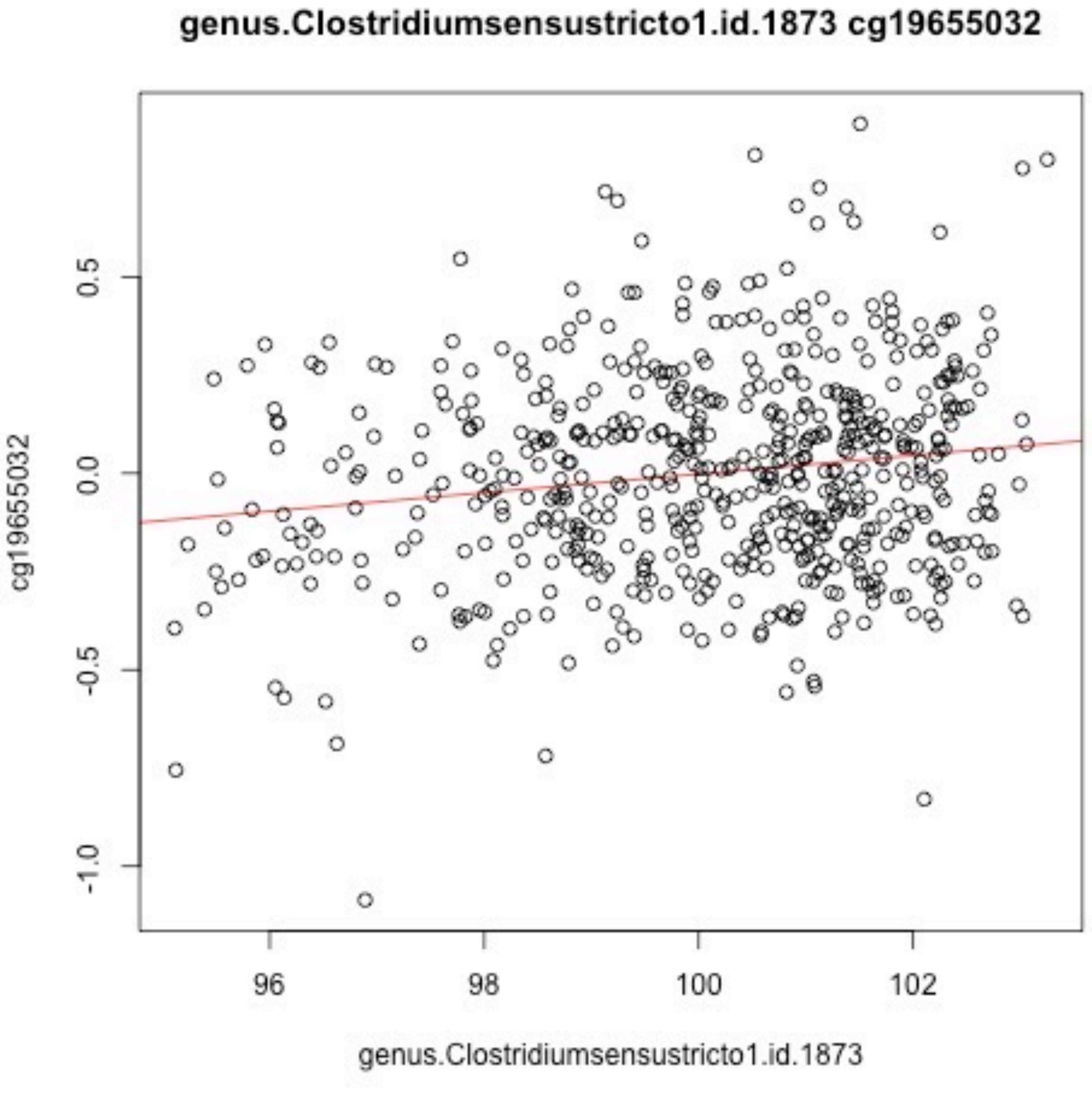
genus.Clostridiumsensustricto1.id.1873 cg19655032
cg19655032
0.5
0.0
-0.5
-1.0
96
98
100
102
genus.Clostridiumsensustricto1.id.1873

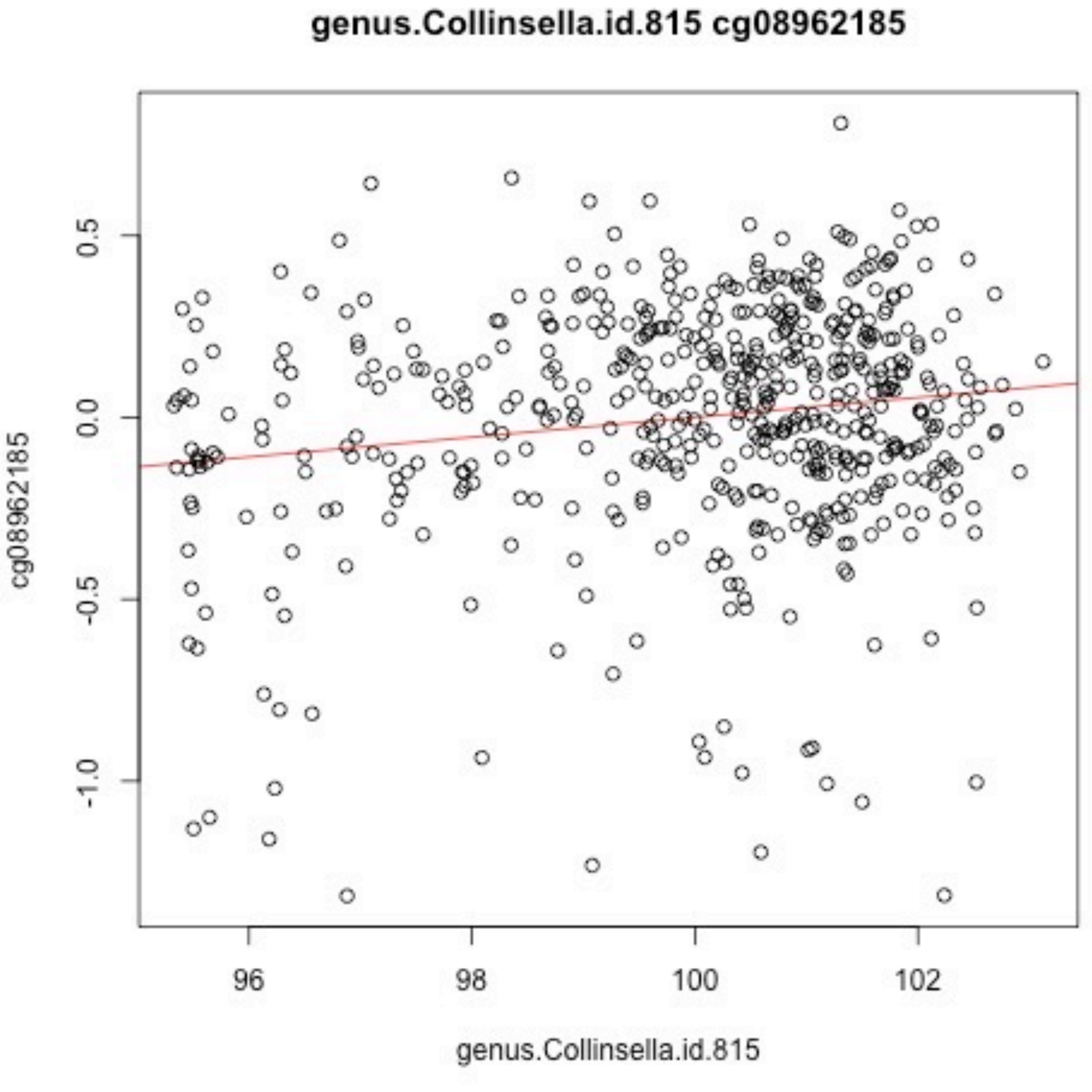
genus.Collinsella.id.815 cg08962185
cg08962185
genus.Collinsella.id.815
0.5
0.0
-0.5
-1.0
96
98
100
102

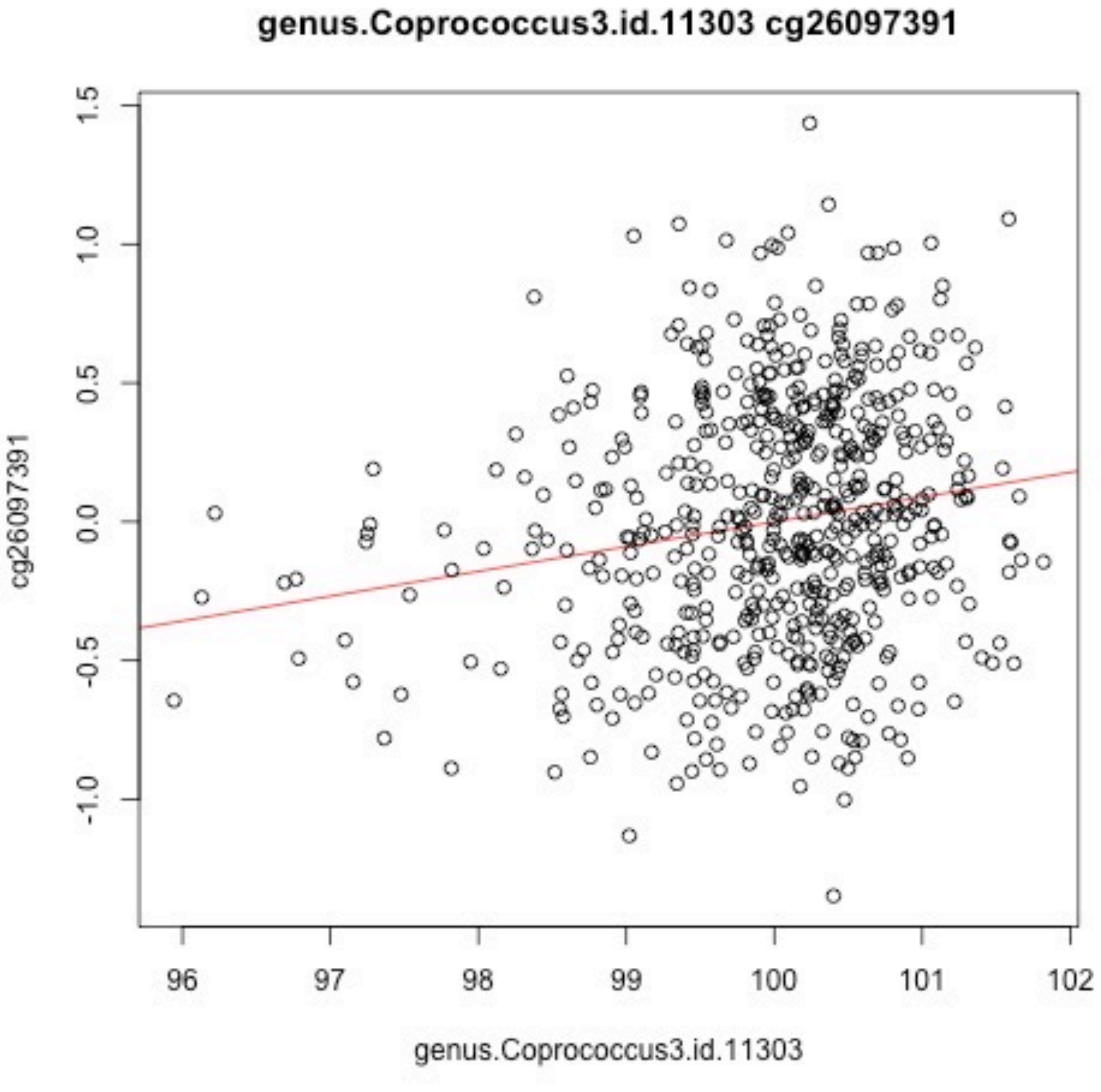
genus.Coprococcus3.id.11303 cg26097391
1.5
1.0
0.5
0.0
-0.5
-1.0
cg26097391
96
97
98
99
100
101
102
genus.Coprococcus3.id.11303

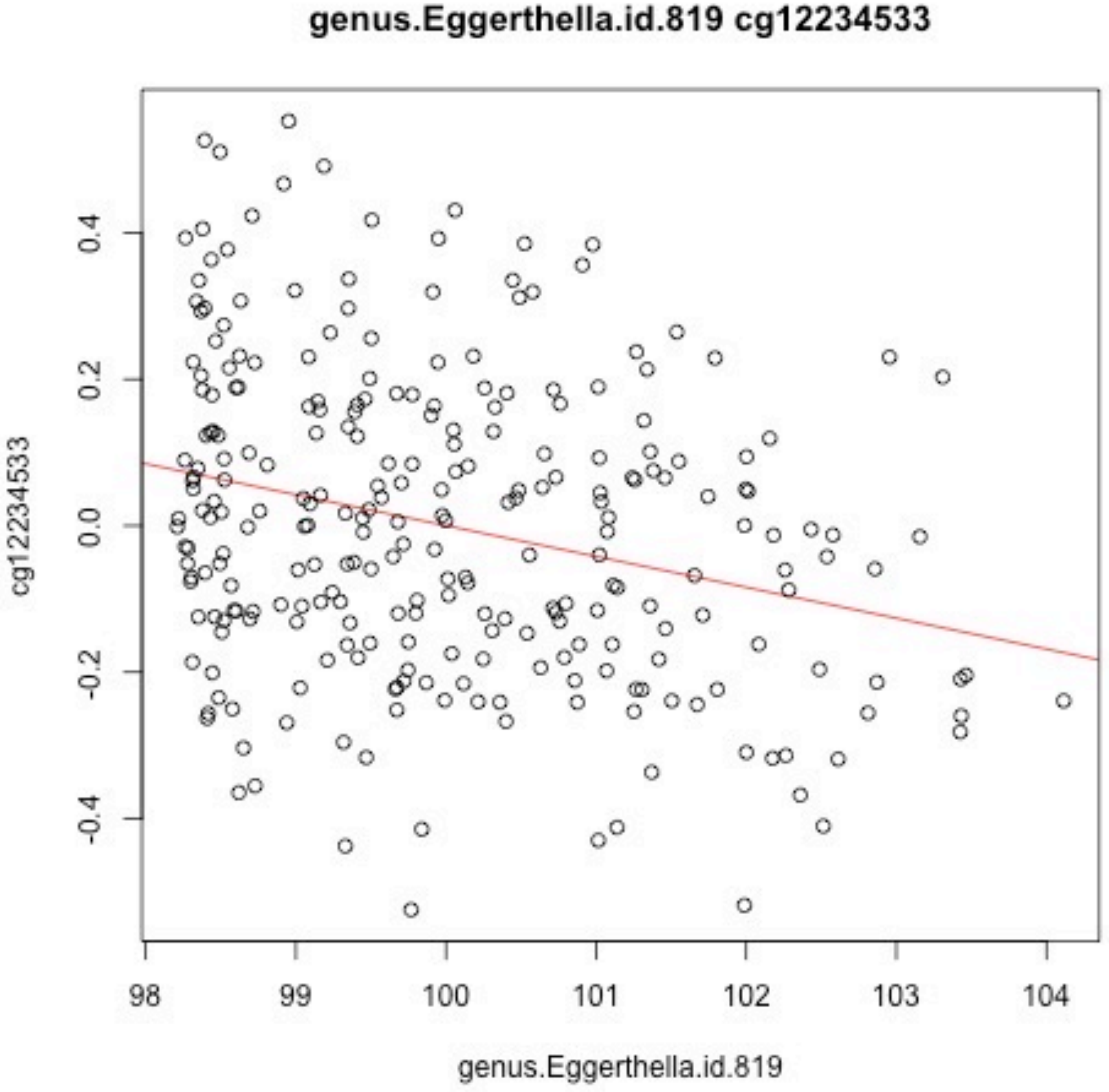
genus.Eggerthella.id.819 cg12234533
cg12234533
0.4
0.2
0.0
-0.2
-0.4
98
99
100
101
102
103
104
genus.Eggerthella.id.819

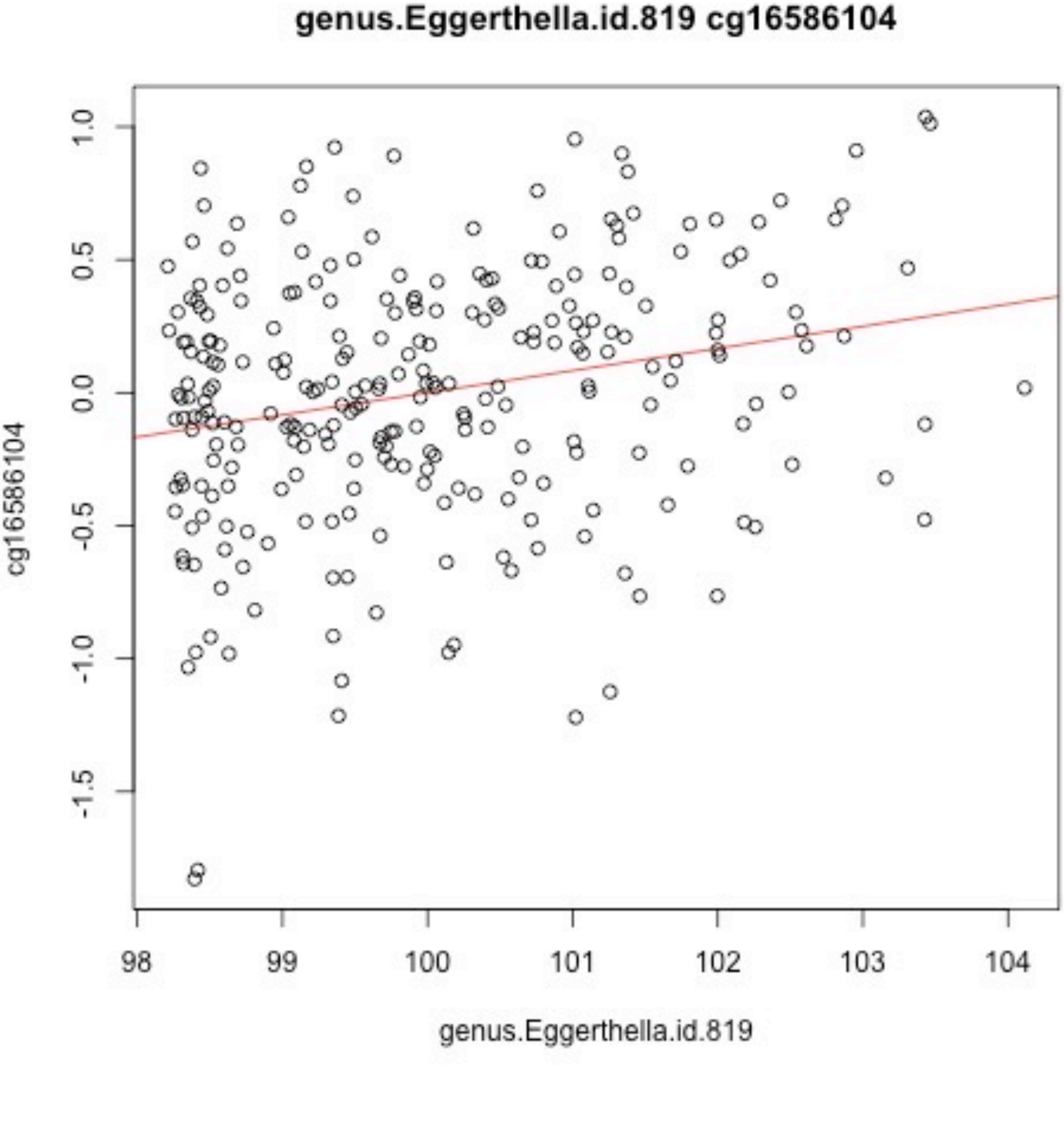
genus.Eggerthella.id.819 cg16586104
cg16586104
1.0
0.5
0.0
-0.5
-1.0
-1.5
98
99
100
101
102
103
104
genus.Eggerthella.id.819

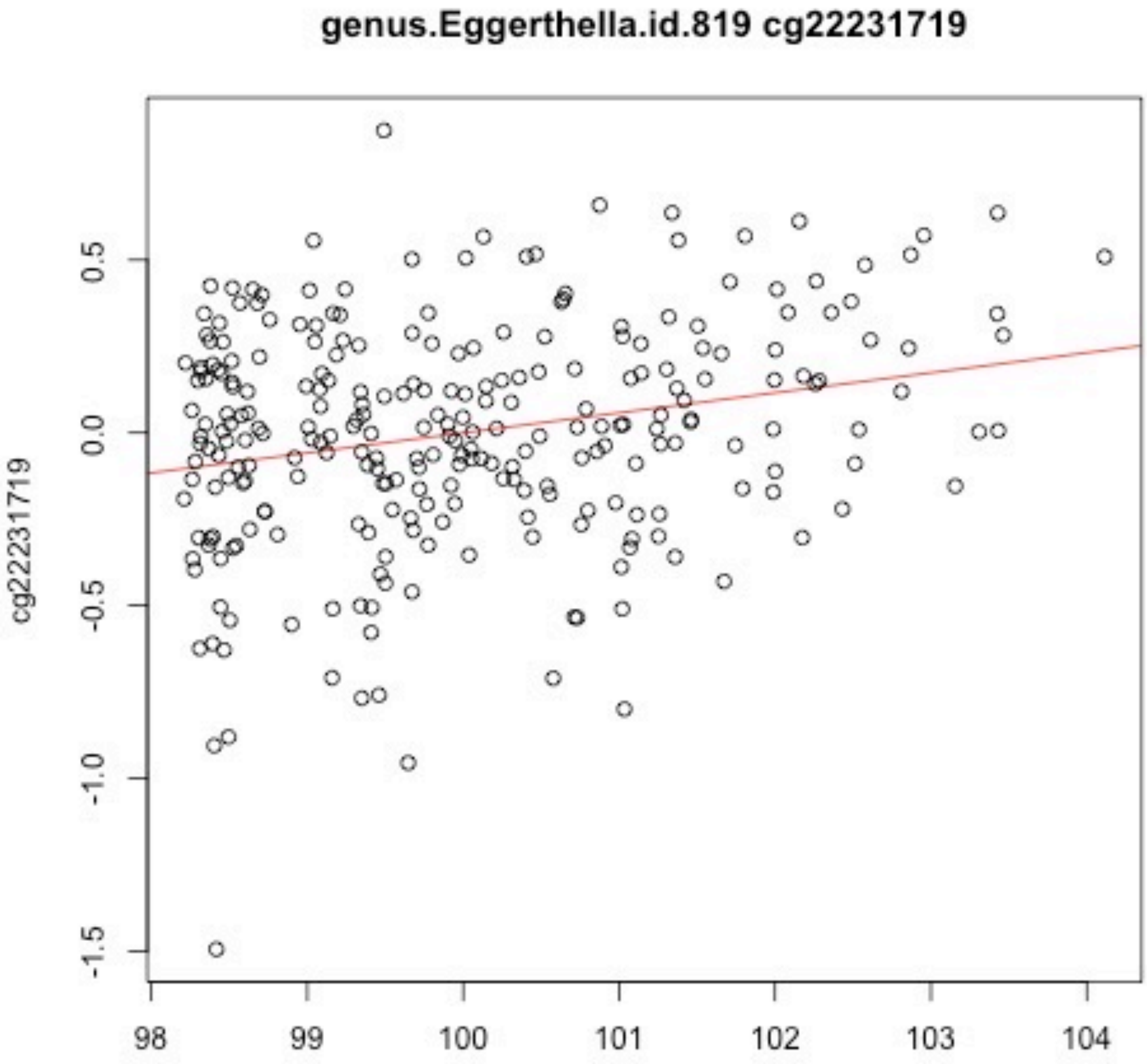
genus.Eggerthella.id.819 cg22231719
cg22231719
0.5
0.0
-0.5
-1.0
-1.5
98
99
100
101
102
103
104
genus.Eggerthella.id.819

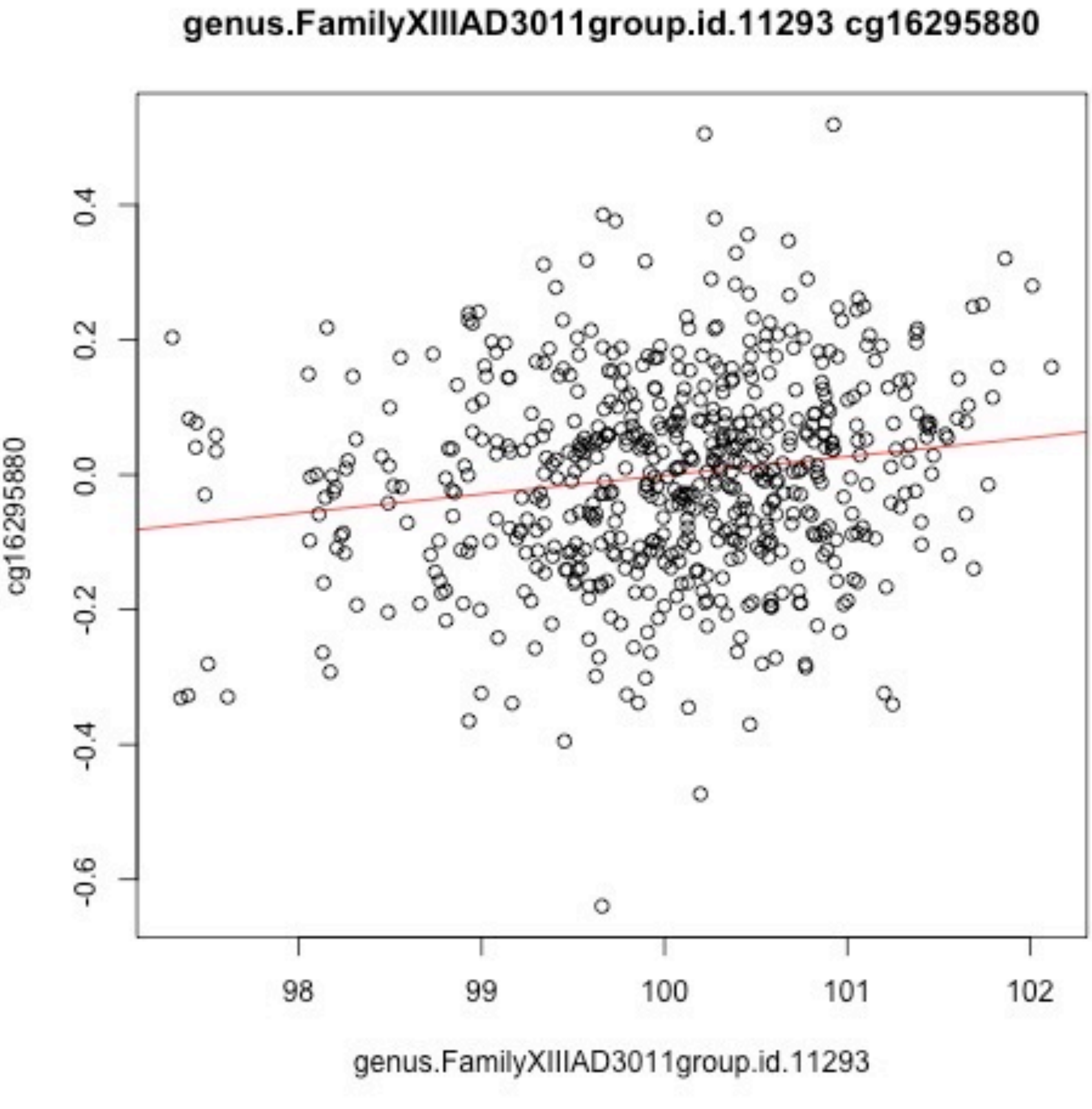
genus.FamilyXIIIAD3011group.id.11293 cg16295880
0.4
0.2
0.0
-0.2
-0.4
-0.6
cg16295880
98
99
100
101
102
genus.FamilyXIIIAD3011group.id.11293

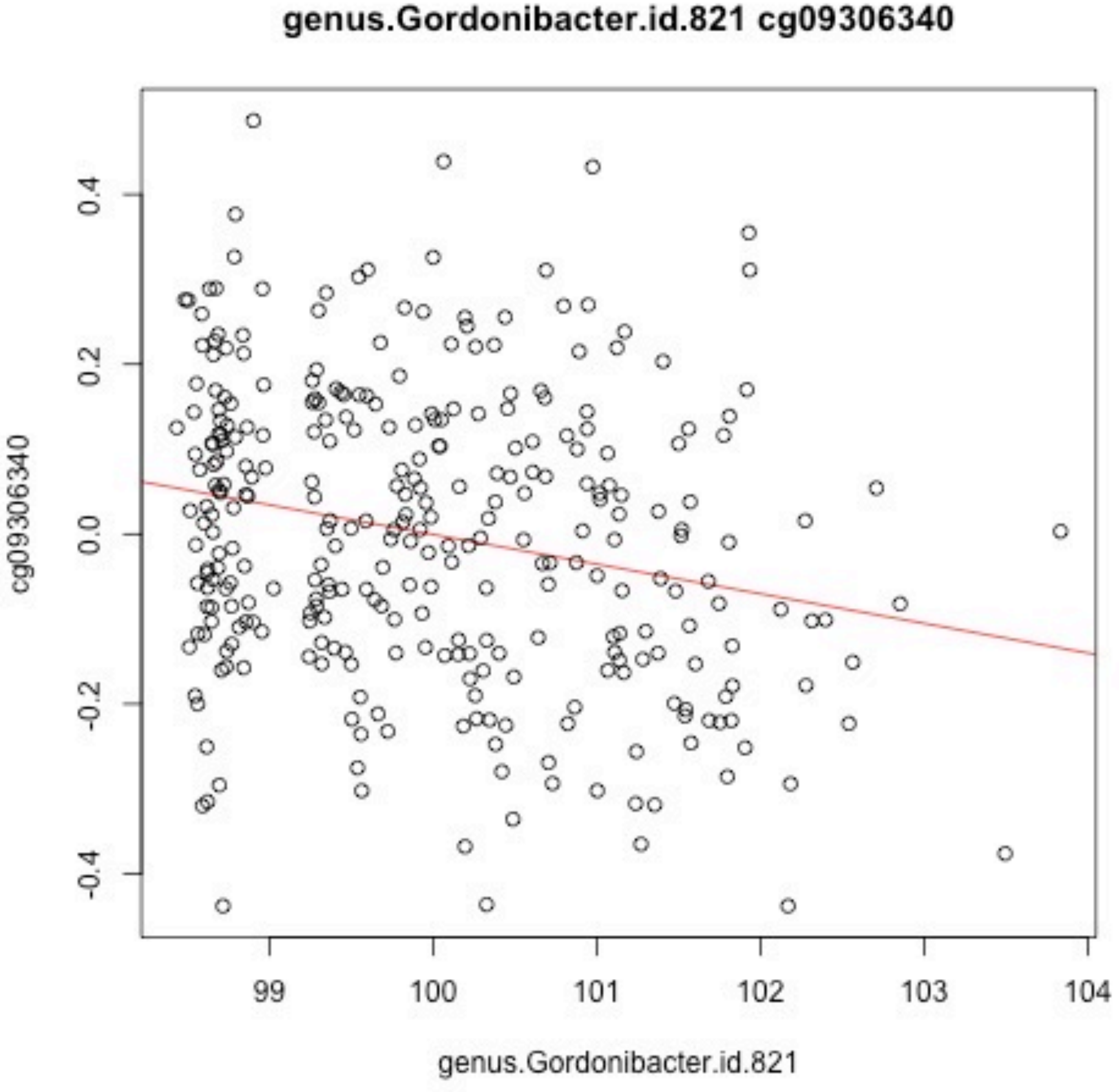

genus.Gordonibacter.id.821 cg09306340
cg09306340
0.4
0.2
0.0
-0.2
-0.4
99
100
101
102
103
104
genus.Gordonibacter.id.821

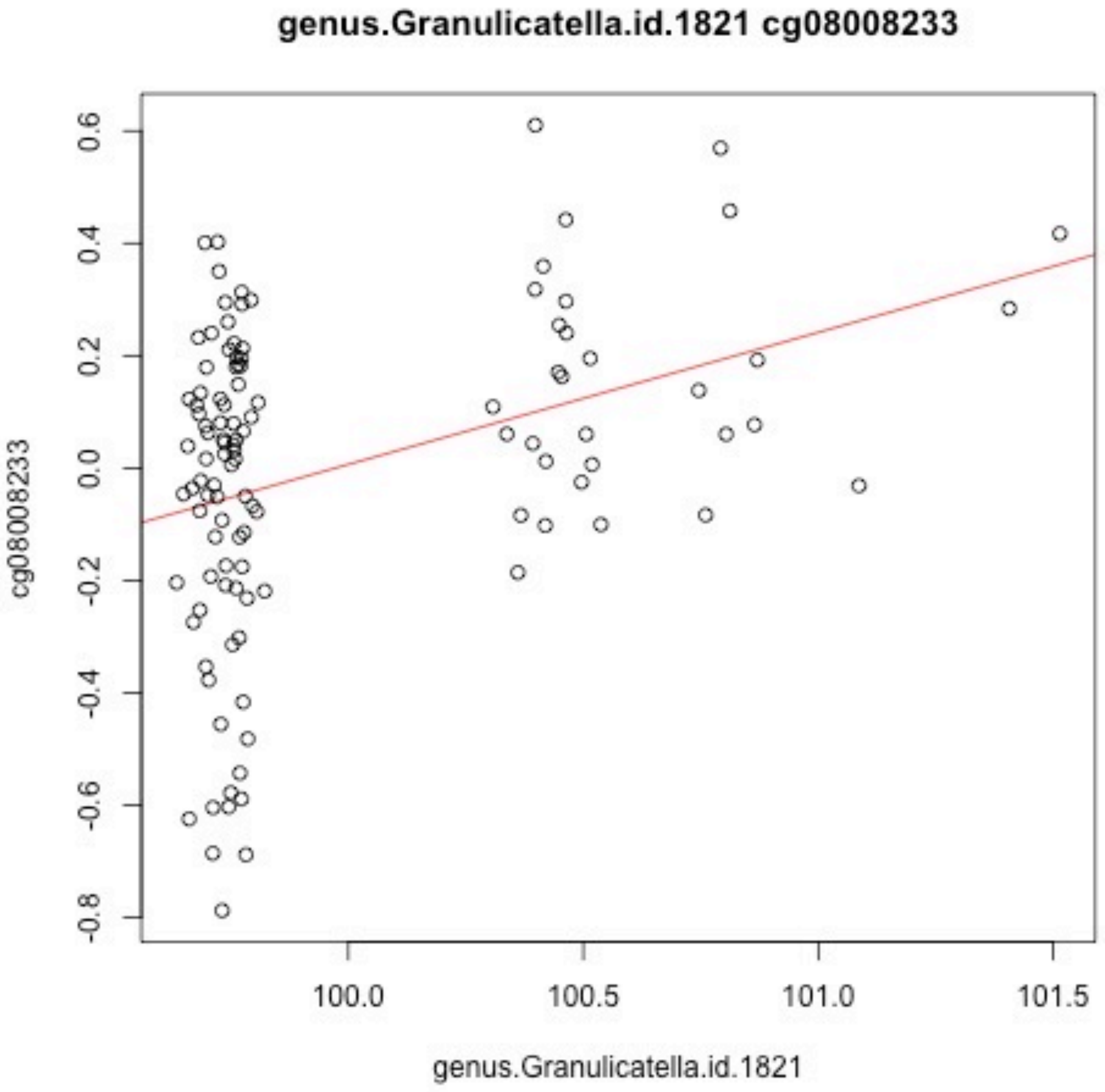
genus.Granulicatella.id.1821 cg08008233
cg08008233
0.6
0.4
0.2
0.0
-0.2
-0.4
-0.6
-0.8
100.0
100.5
101.0
101.5
genus.Granulicatella.id.1821

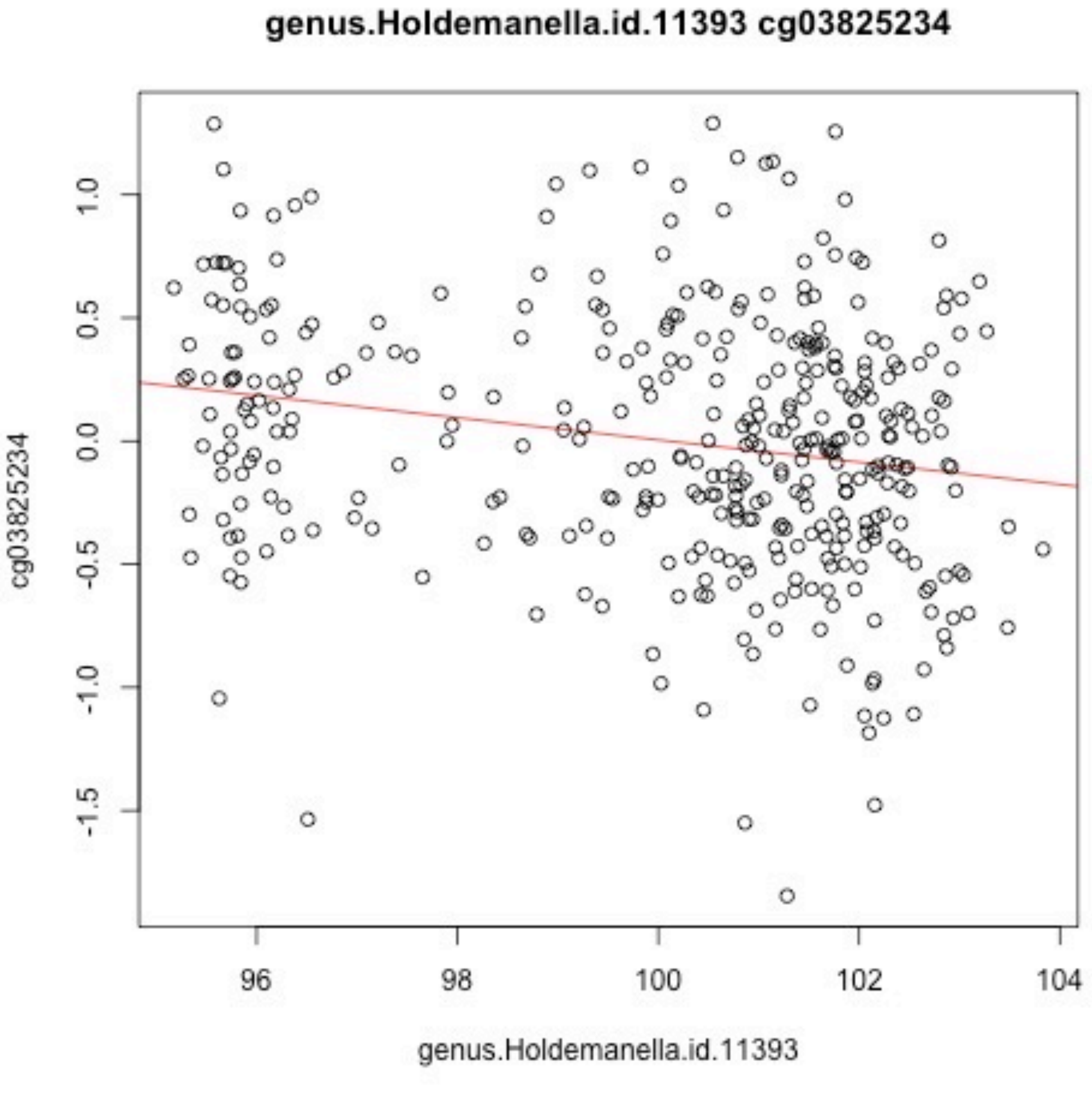
genus.Holdemanella.id.11393 cg03825234
cg03825234
genus.Holdemanella.id.11393
1.0
0.5
0.0
-0.5
-1.0
-1.5
96
98
100
102
104

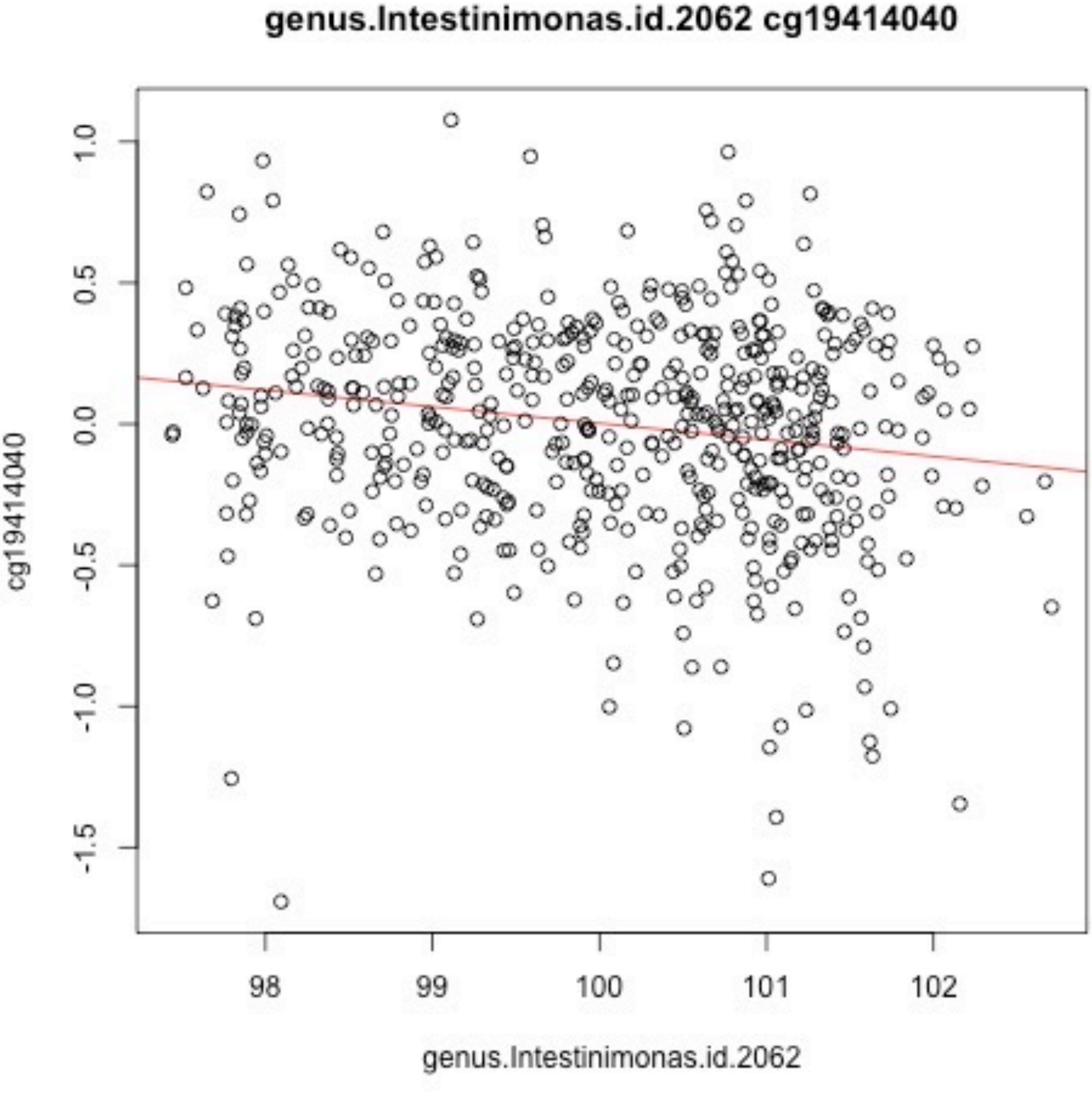

genus.Intestinimonas.id.2062 cg19414040
cg19414040
1.0
0.5
0.0
-0.5
-1.0
-1.5
98
99
100
101
102
genus.Intestinimonas.id.2062

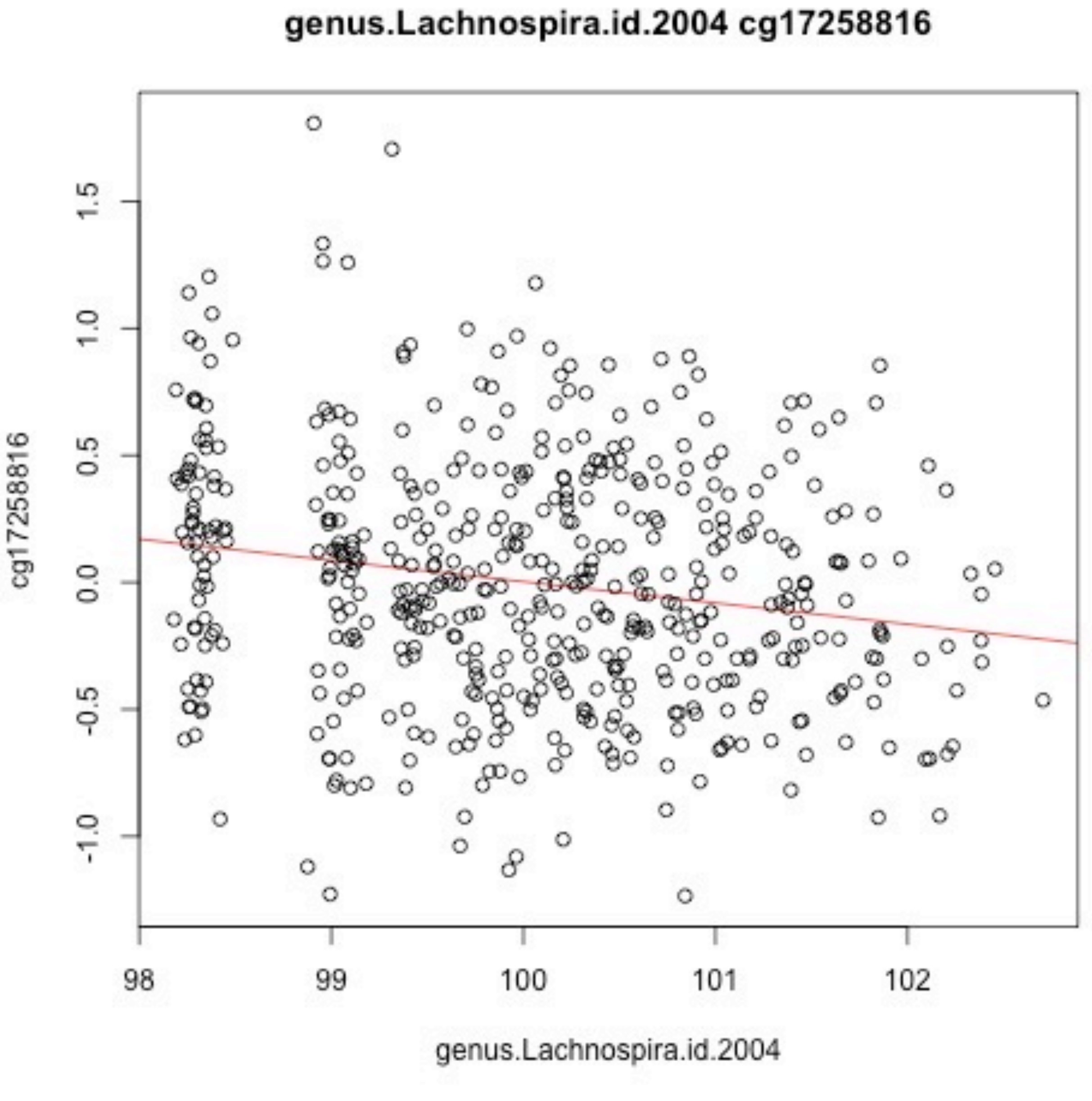
genus.Lachnospira.id.2004 cg17258816
cg17258816
1.5
1.0
0.5
0.0
-0.5
-1.0
98
99
100
101
102
genus.Lachnospira.id.2004

### genus.LachnospiraceaeNC2004group.id.11316 cg00278329

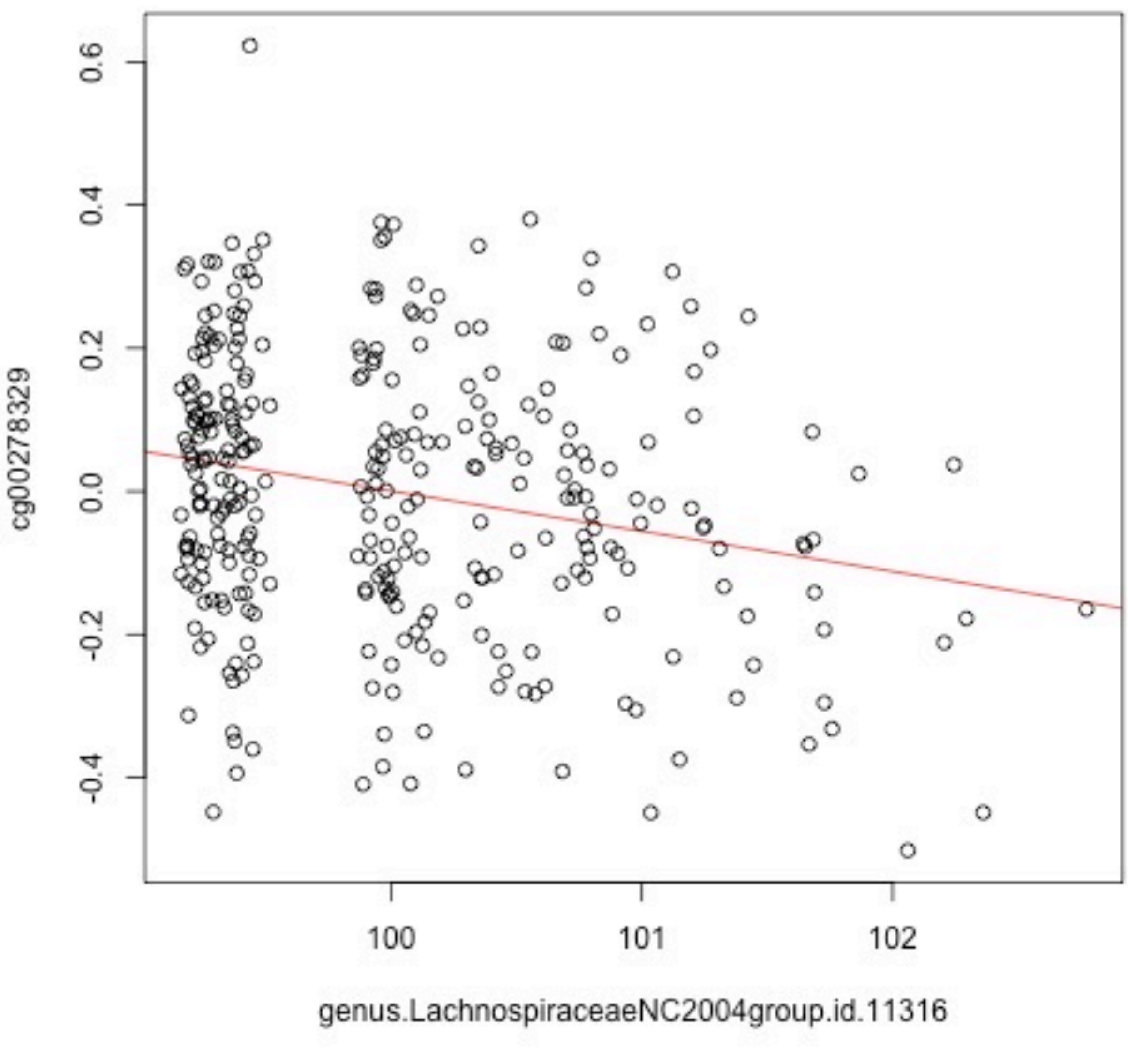

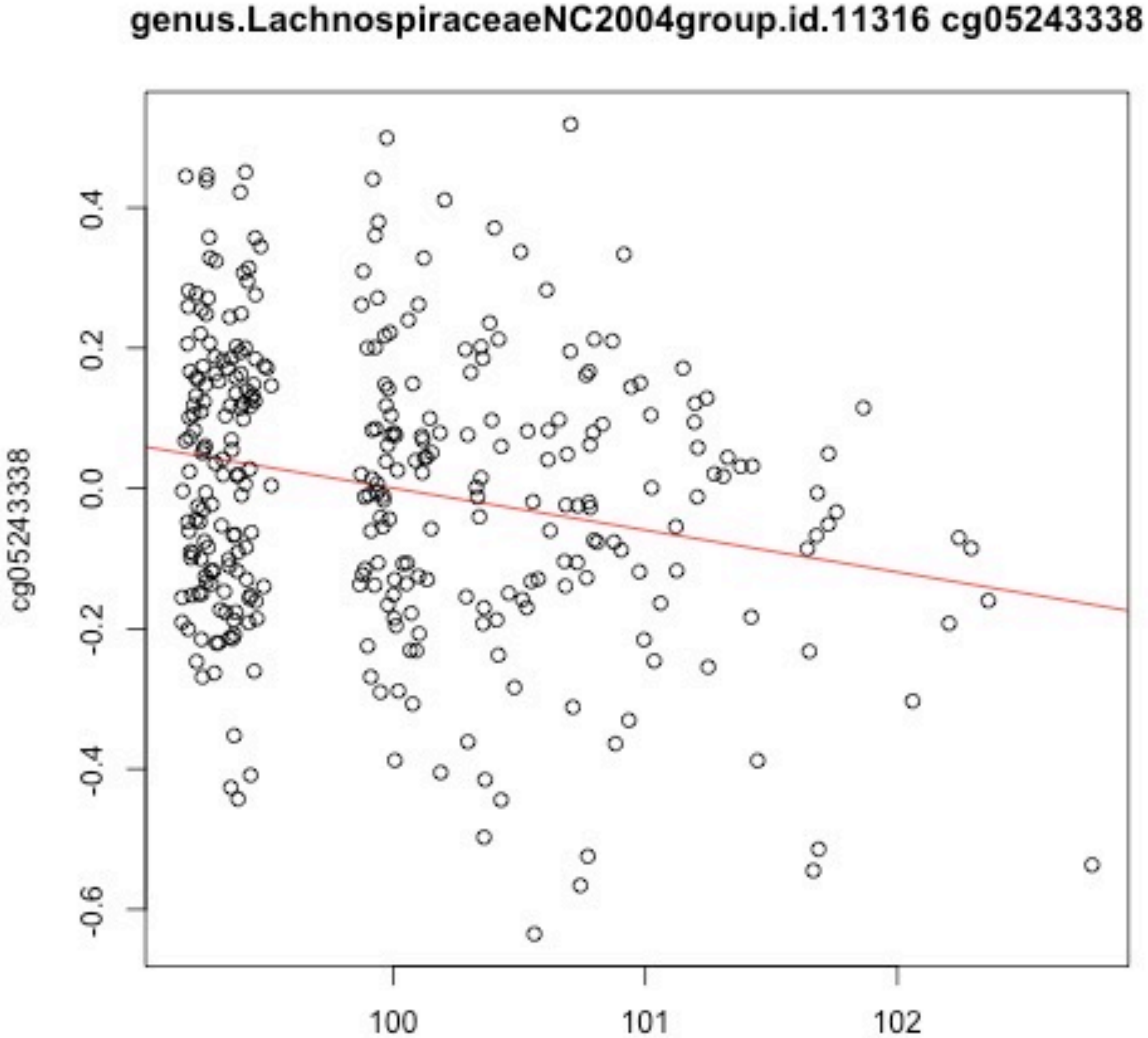
genus.LachnospiraceaeNC2004group.id.11316 cg05243338
cg05243338
0.4
0.2
0.0
-0.2
-0.4
-0.6
100
101
102
genus.LachnospiraceaeNC2004group.id.11316

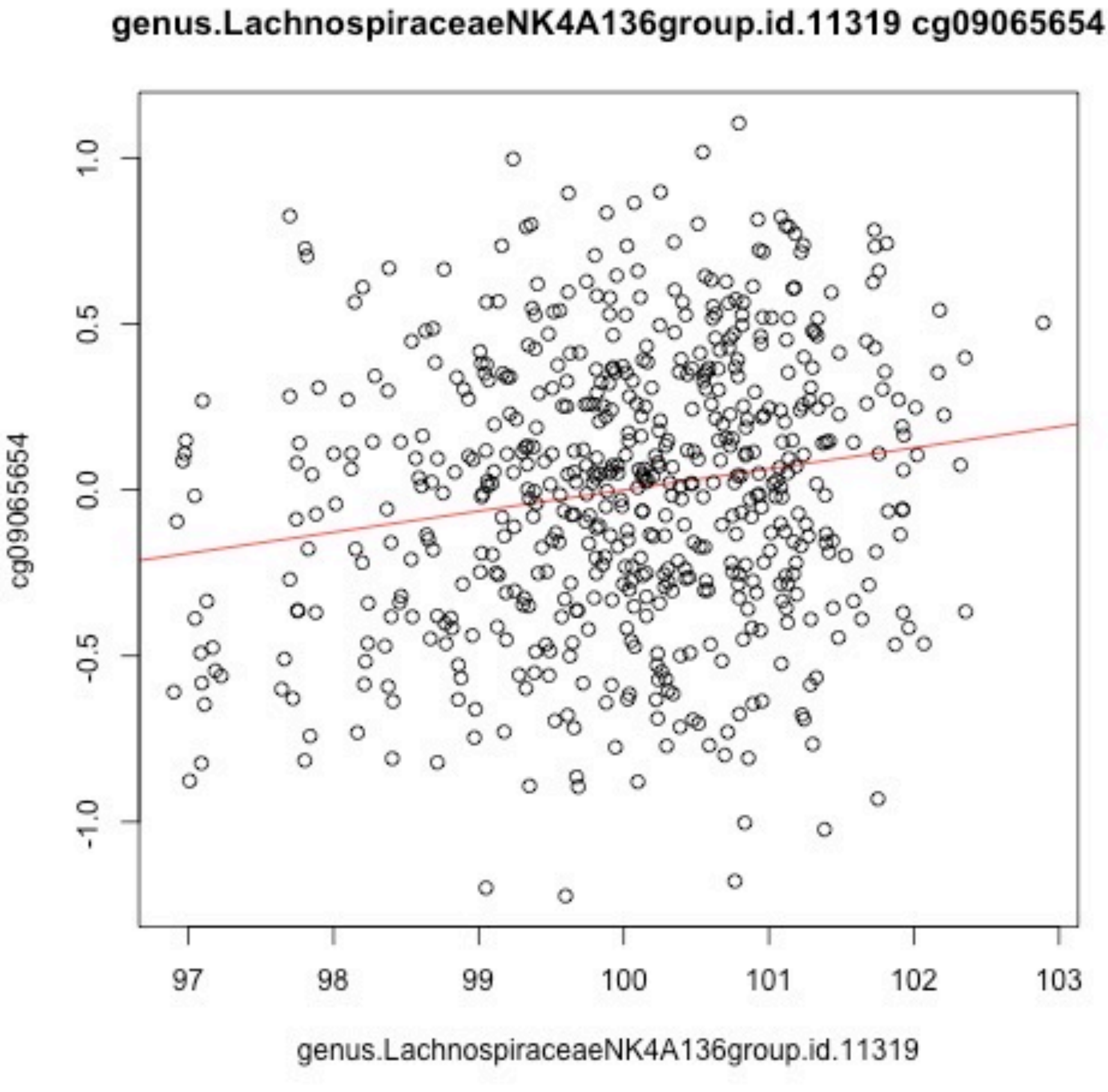
genus.LachnospiraceaeNK4A136group.id.11319 cg09065654
cg09065654
1.0
0.5
0.0
-0.5
-1.0
97
98
99
100
101
102
103
genus.LachnospiraceaeNK4A136group.id.11319

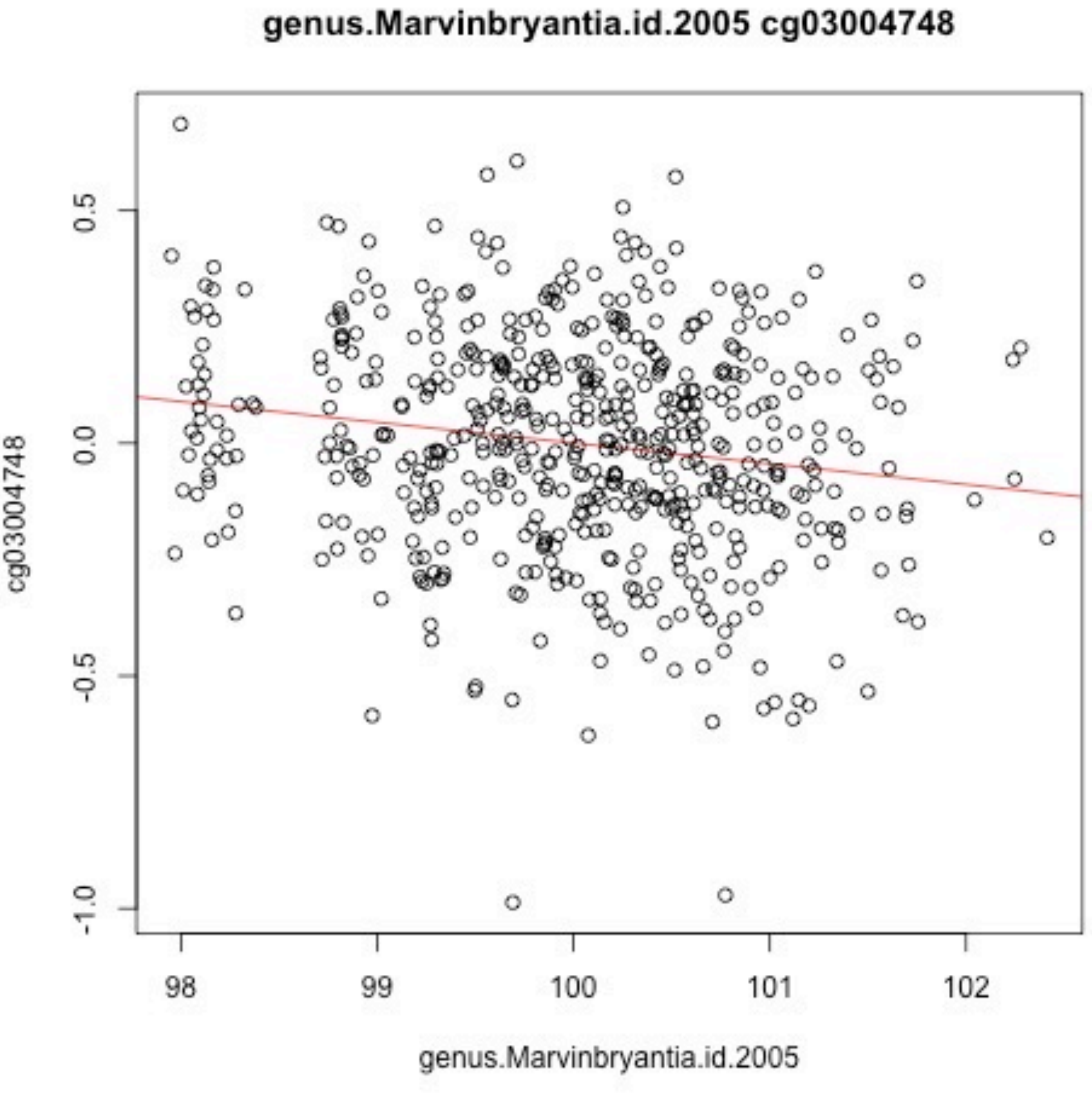
genus.Marvinbryantia.id.2005 cg03004748
cg03004748
0.5
0.0
-0.5
-1.0
98
99
100
101
102
genus.Marvinbryantia.id.2005

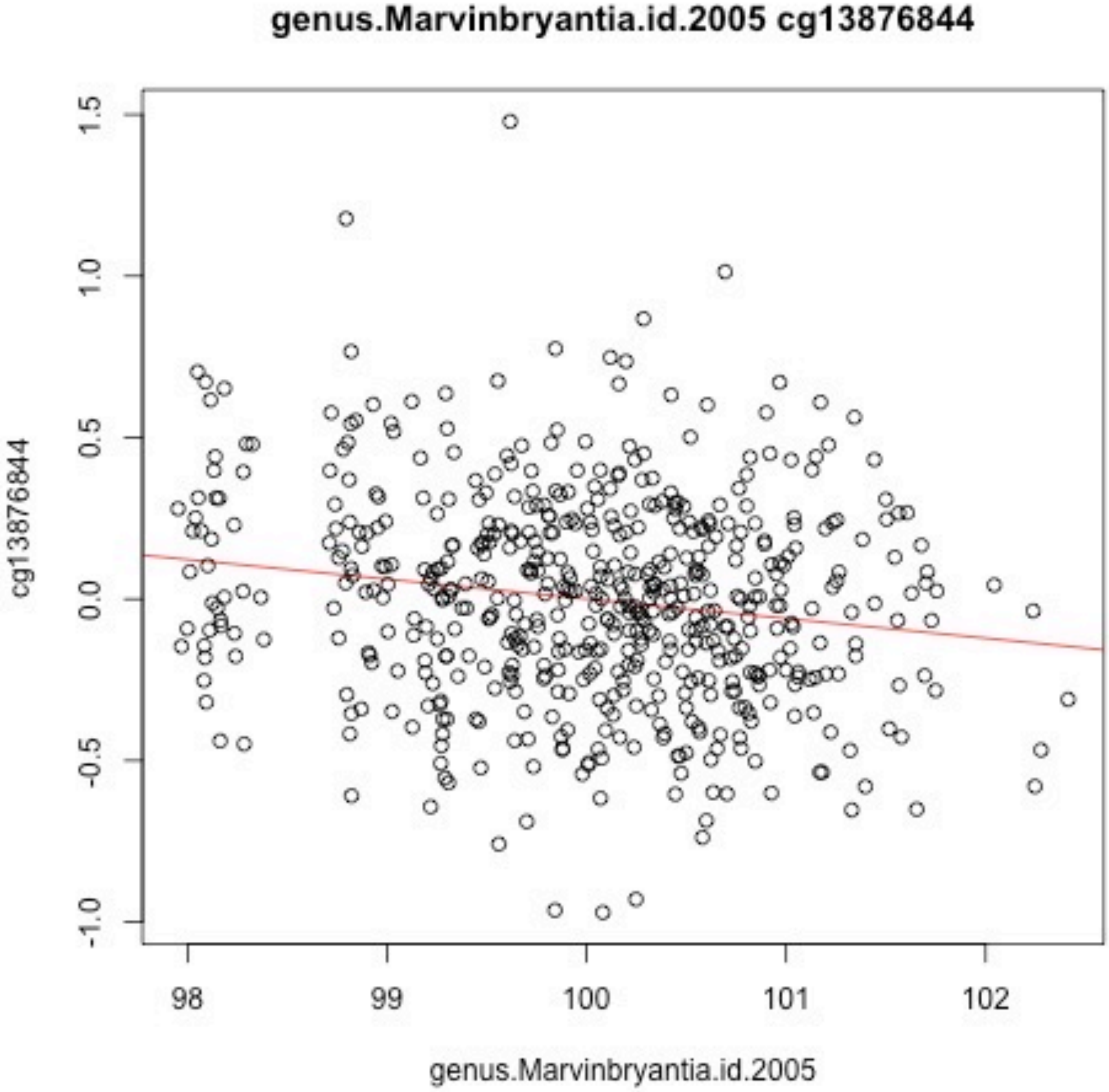
genus.Marvinbryantia.id.2005 cg13876844
cg13876844
1.5
1.0
0.5
0.0
-0.5
-1.0
98
99
100
101
102
genus.Marvinbryantia.id.2005

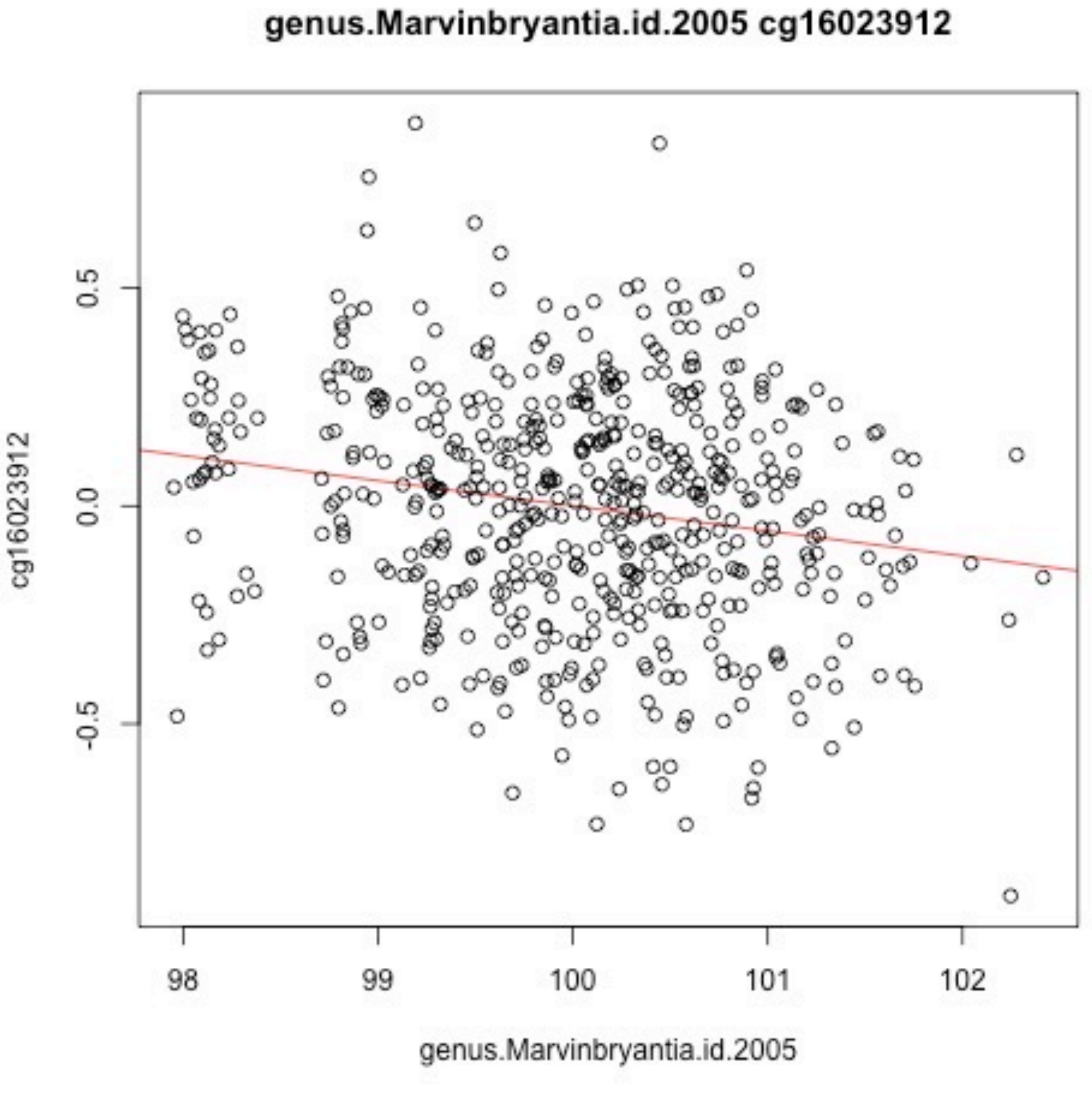
genus.Marvinbryantia.id.2005 cg16023912
cg16023912
0.5
0.0
-0.5
98
99
100
101
102
genus.Marvinbryantia.id.2005

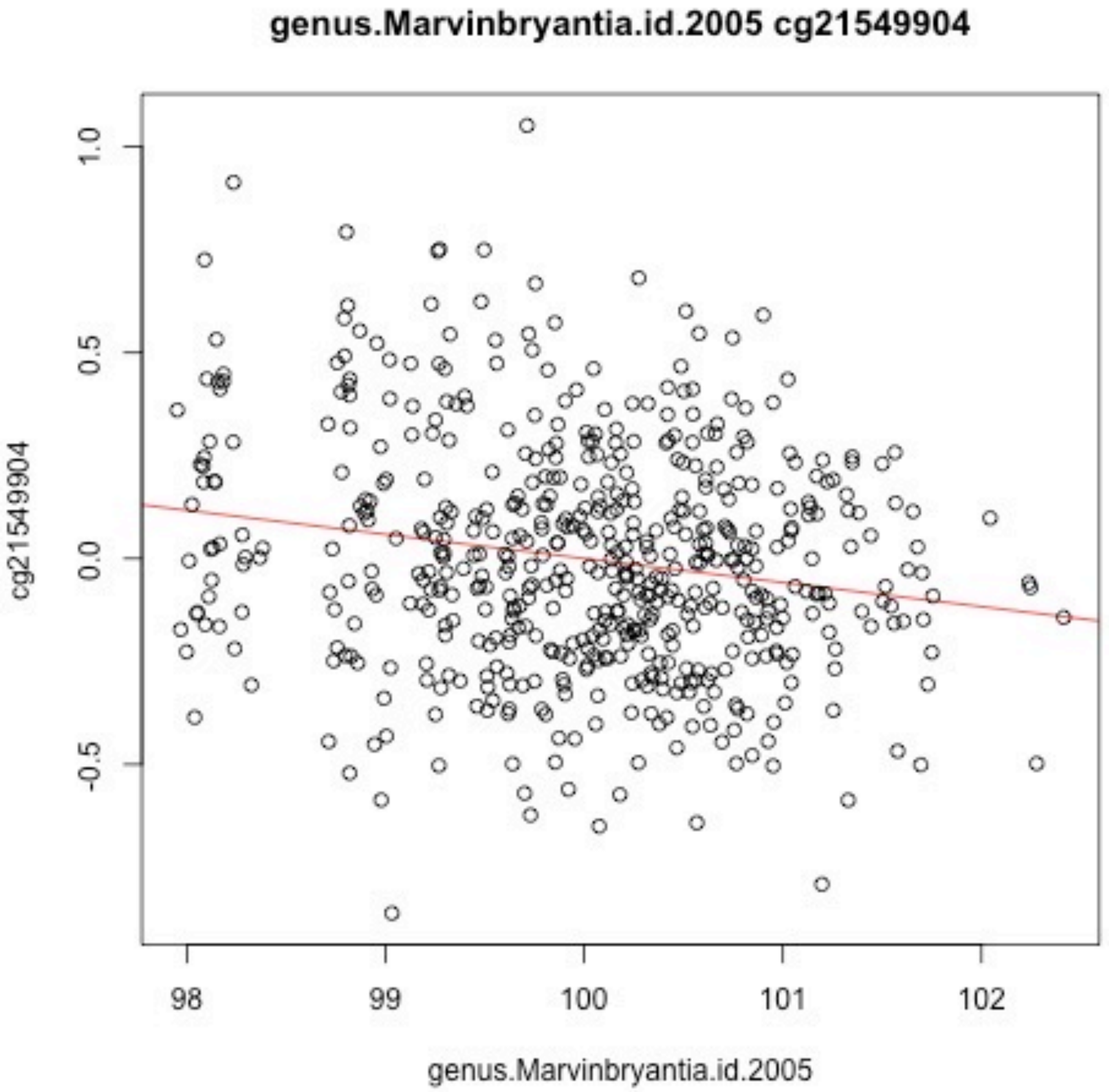
genus.Marvinbryantia.id.2005 cg21549904
cg21549904
1.0
0.5
0.0
-0.5
98
99
100
101
102
genus.Marvinbryantia.id.2005

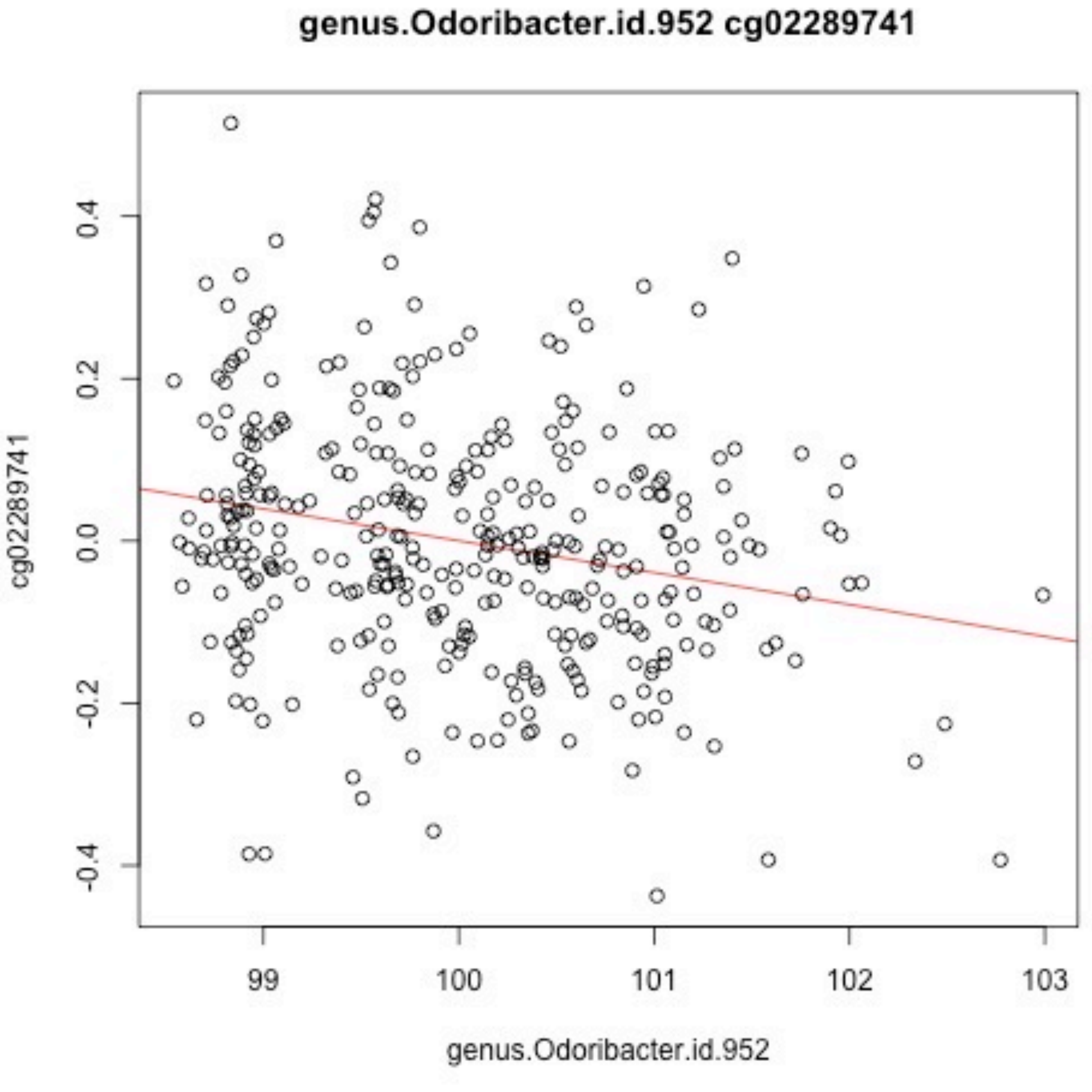

genus.Odoribacter.id.952 cg02289741
cg02289741
genus.Odoribacter.id.952
-0.4
-0.2
0.0
0.2
0.4
99
100
101
102
103

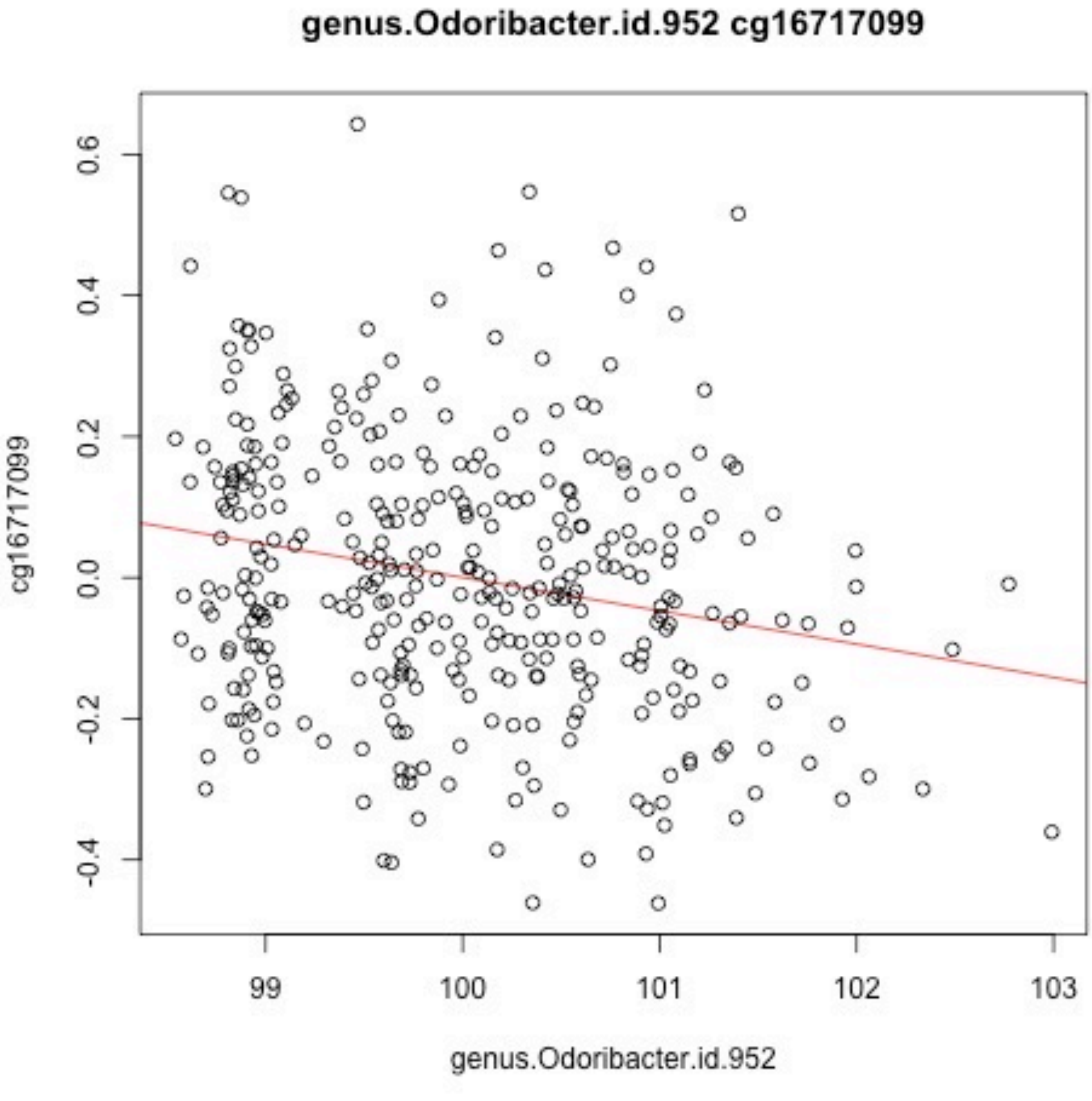

genus.Odoribacter.id.952 cg16717099
cg16717099
0.6
0.4
0.2
0.0
-0.2
-0.4
99
100
101
102
103
genus.Odoribacter.id.952

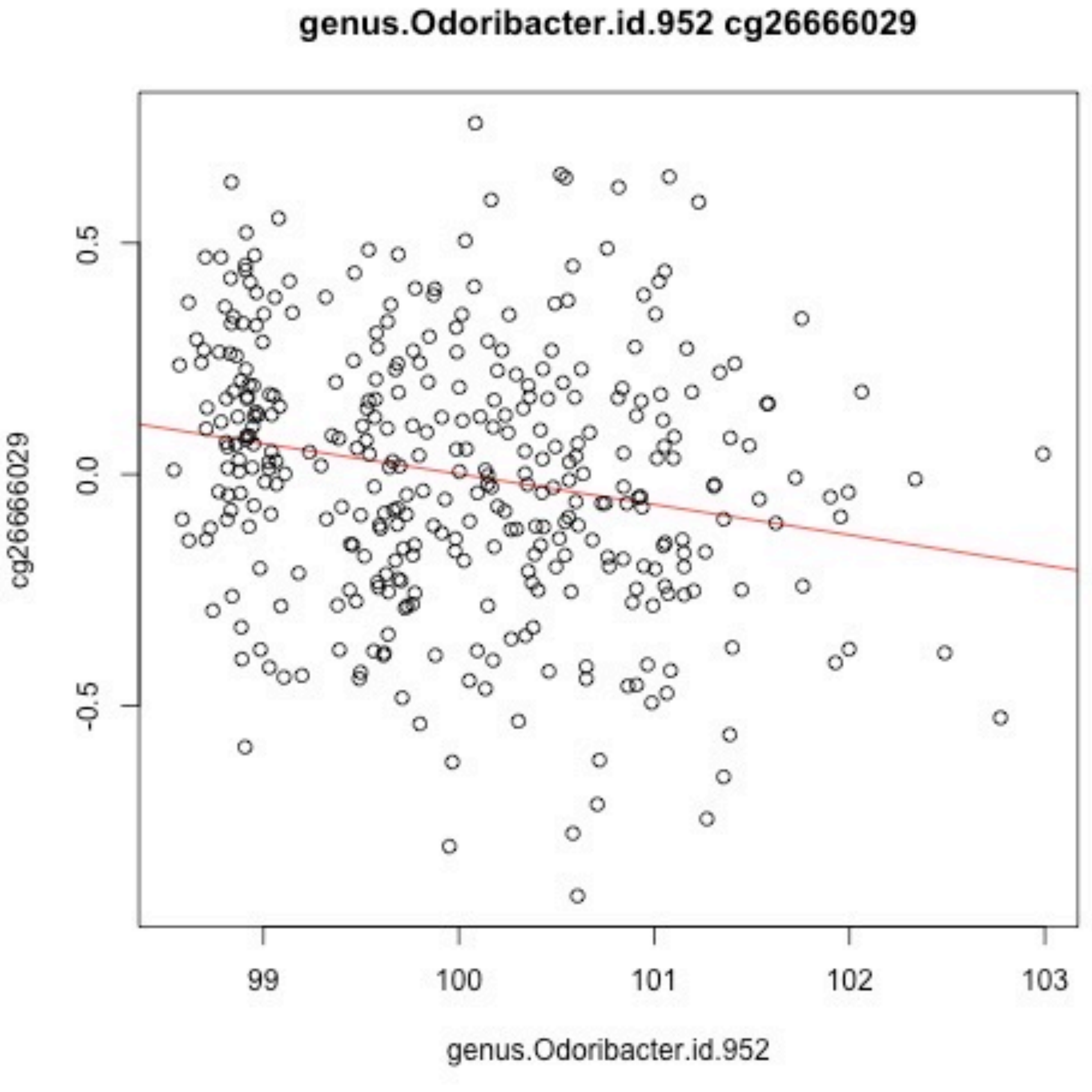
genus.Odoribacter.id.952 cg26666029
cg26666029
0.5
0.0
-0.5
99
100
101
102
103
genus.Odoribacter.id.952

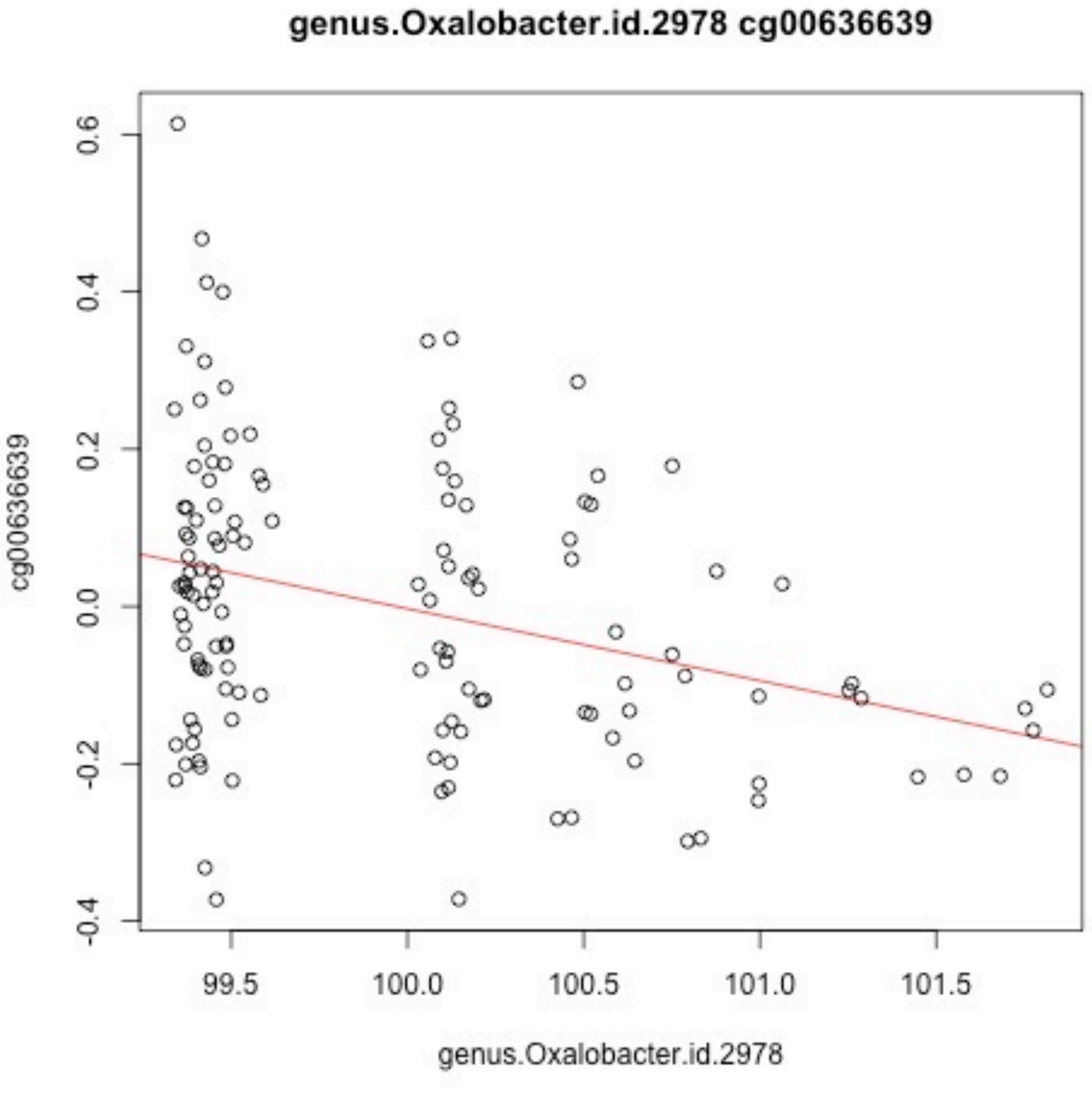
genus.Oxalobacter.id.2978 cg00636639
cg00636639
0.6
0.4
0.2
0.0
-0.2
-0.4
99.5
100.0
100.5
101.0
101.5
genus.Oxalobacter.id.2978

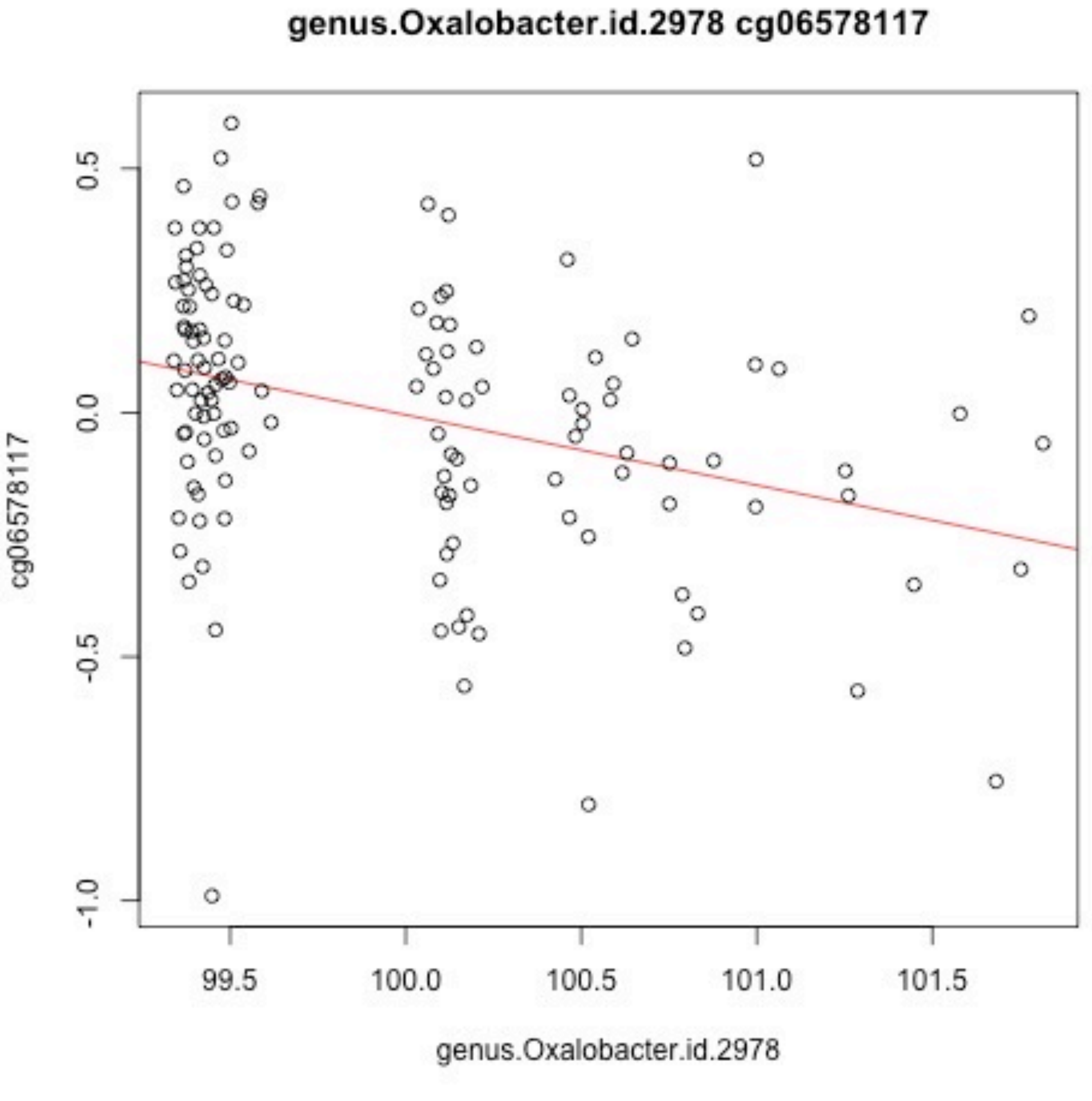
genus.Oxalobacter.id.2978 cg06578117
cg06578117
0.5
0.0
-0.5
-1.0
99.5
100.0
100.5
101.0
101.5
genus.Oxalobacter.id.2978

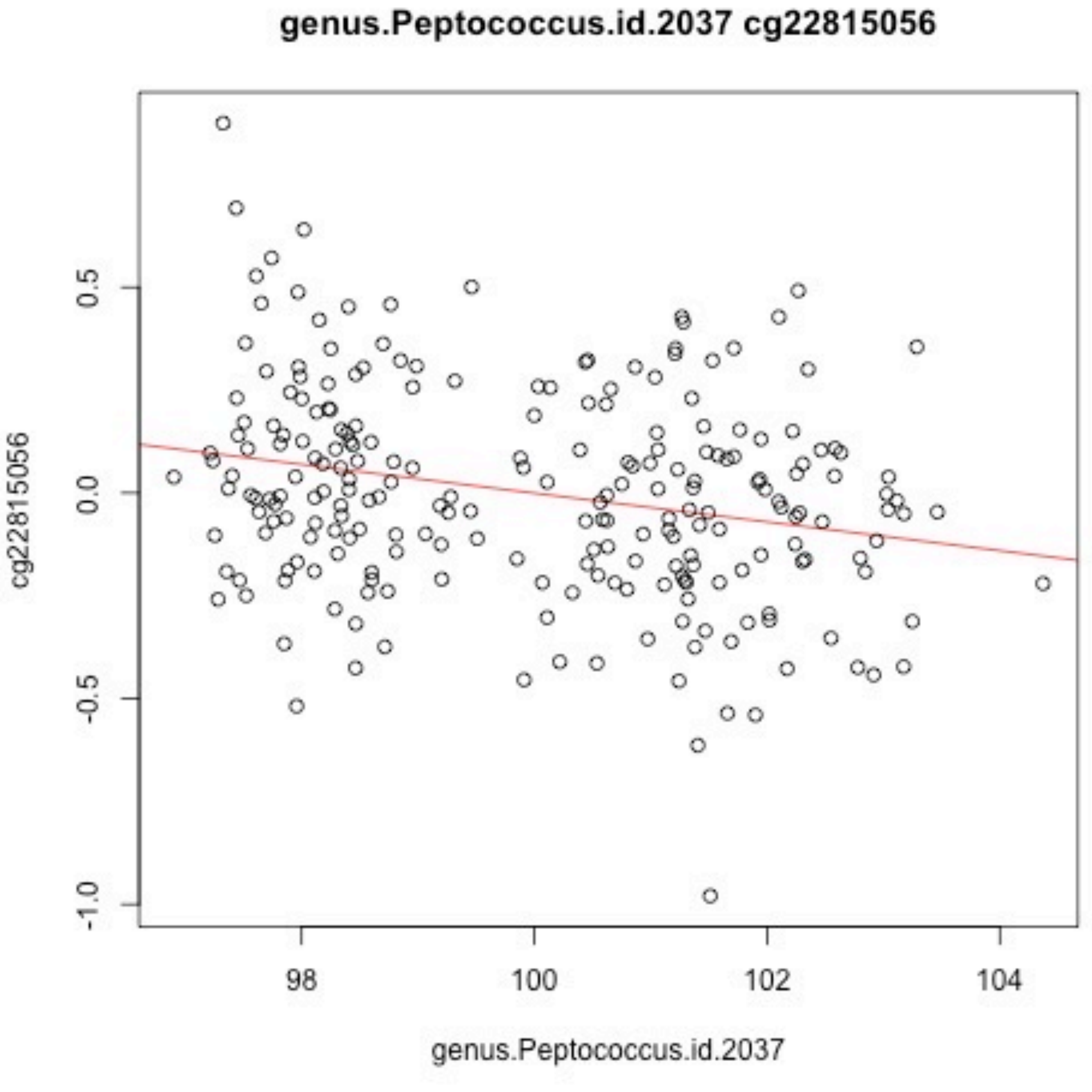
genus.Peptococcus.id.2037 cg22815056
cg22815056
0.5
0.0
-0.5
-1.0
98
100
102
104
genus.Peptococcus.id.2037

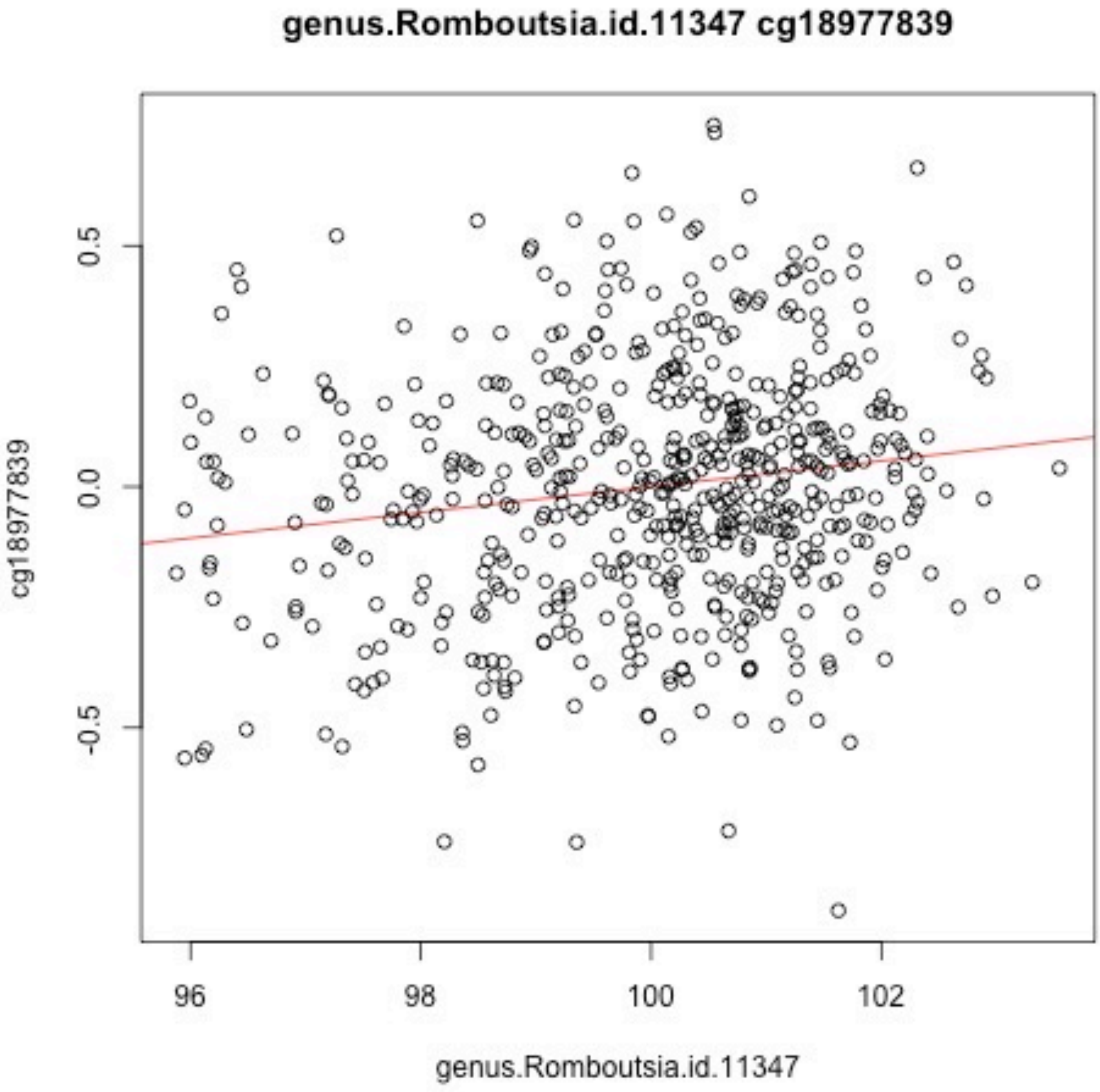
genus.Romboutsia.id.11347 cg18977839
cg18977839
0.5
0.0
-0.5
96
98
100
102
genus.Romboutsia.id.11347

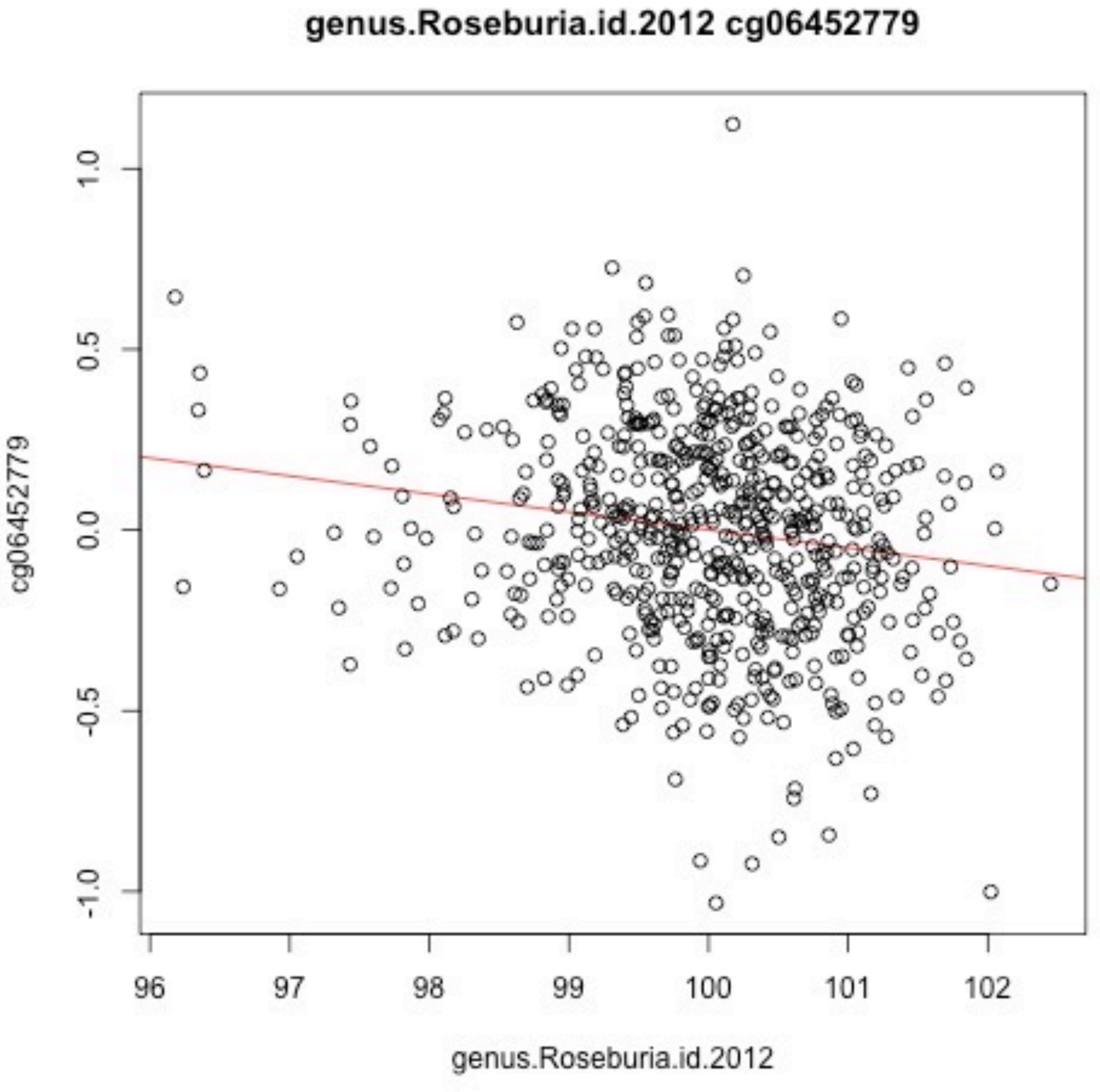
genus.Roseburia.id.2012 cg06452779
cg06452779
1.0
0.5
0.0
-0.5
-1.0
96
97
98
99
100
101
102
genus.Roseburia.id.2012

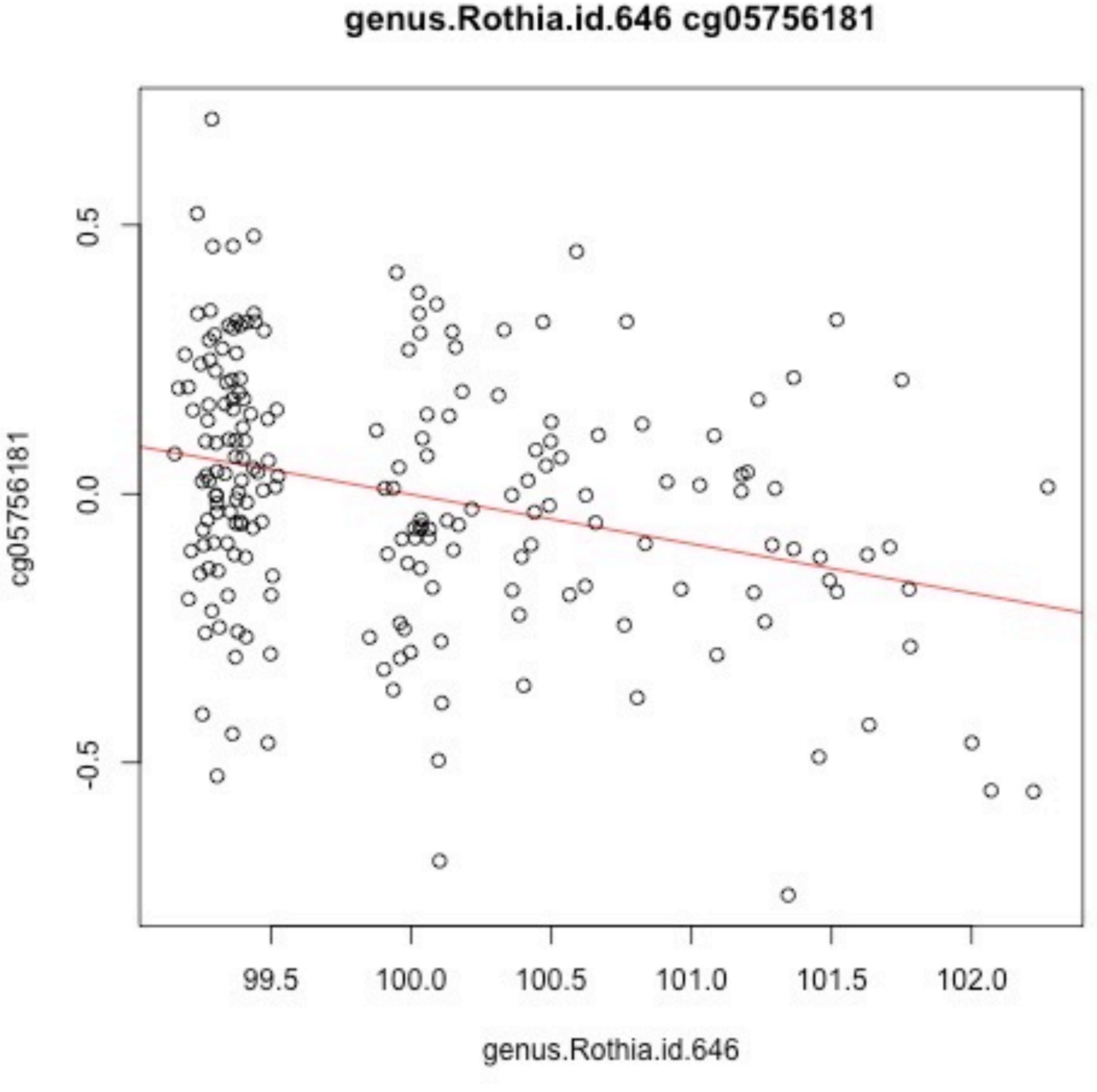
genus.Rothia.id.646 cg05756181
cg05756181
0.5
0.0
-0.5
99.5
100.0
100.5
101.0
101.5
102.0
genus.Rothia.id.646

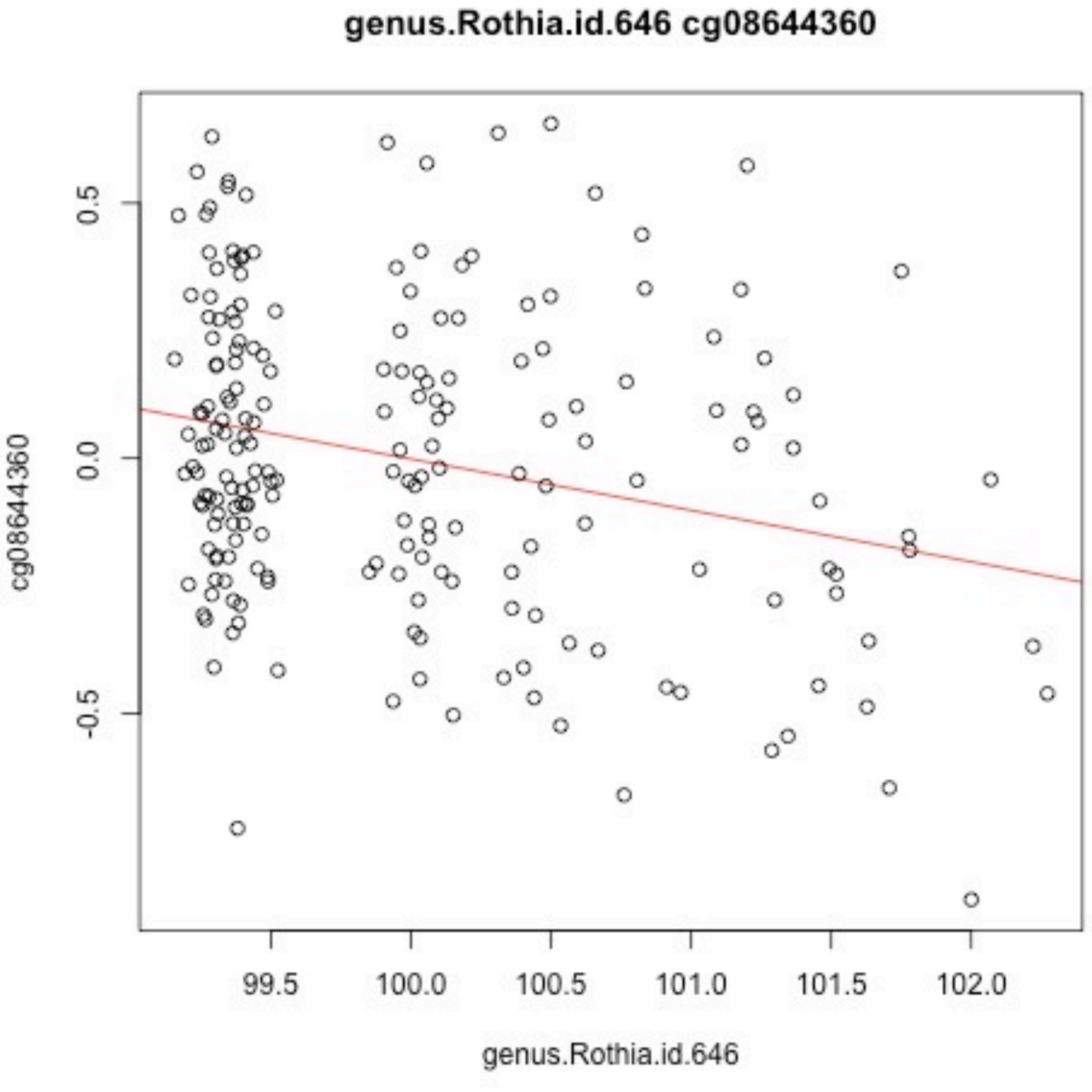

genus.Rothia.id.646 cg08644360
cg08644360
0.5
0.0
-0.5
99.5
100.0
100.5
101.0
101.5
102.0
genus.Rothia.id.646

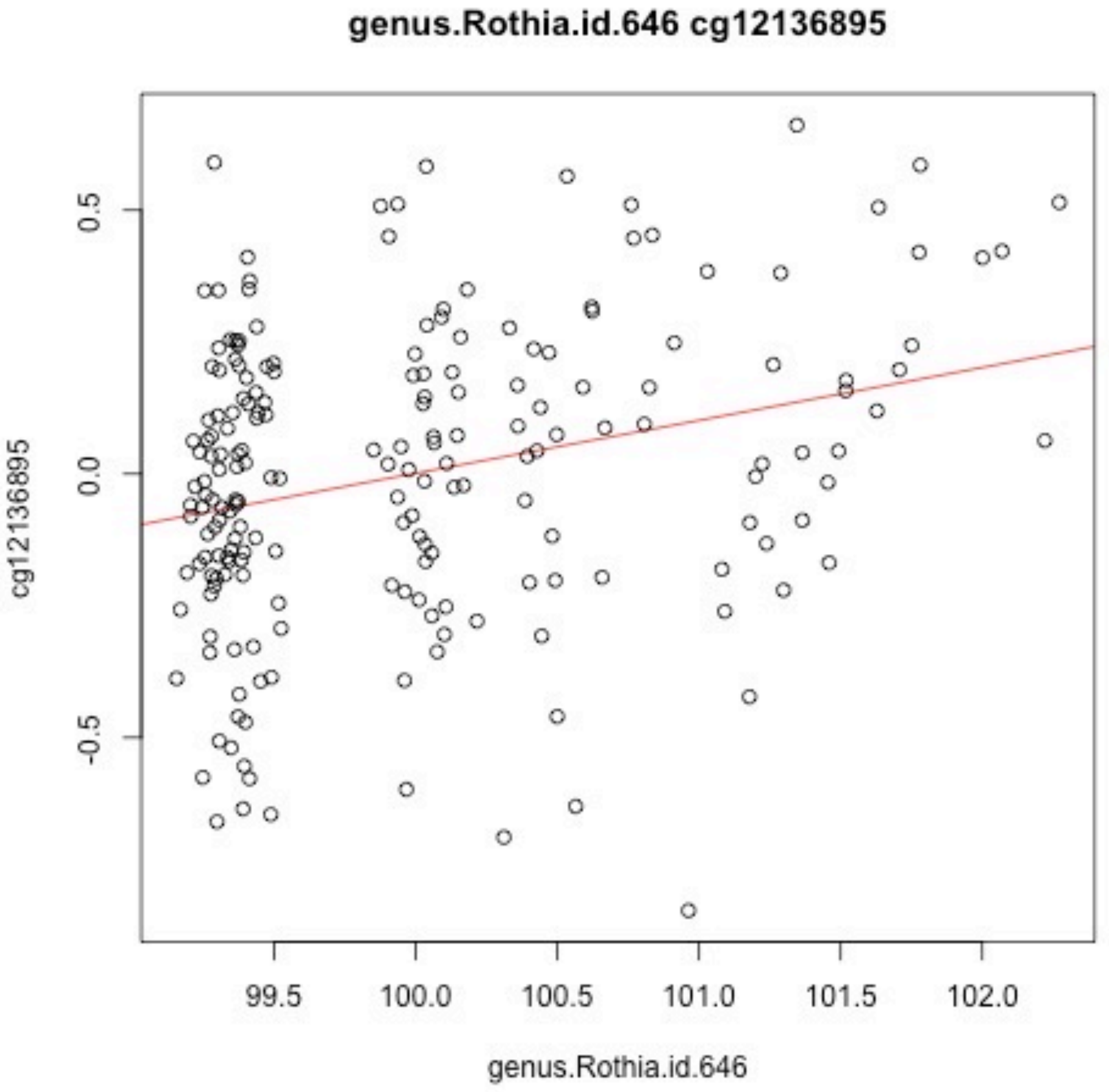
genus.Rothia.id.646 cg12136895
cg12136895
0.5
0.0
-0.5
99.5
100.0
100.5
101.0
101.5
102.0
genus.Rothia.id.646

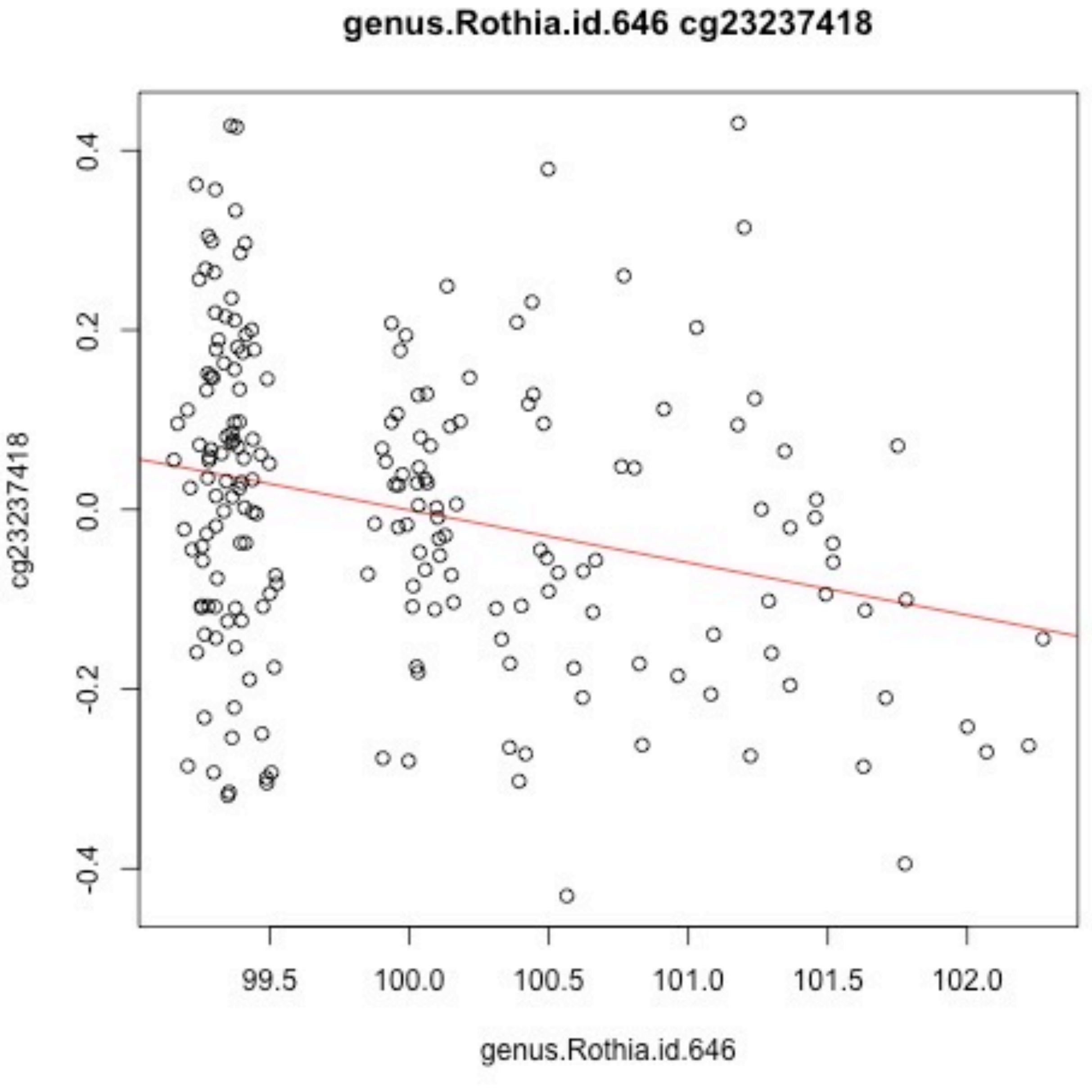
genus.Rothia.id.646 cg23237418
cg23237418
0.4
0.2
0.0
-0.2
-0.4
99.5
100.0
100.5
101.0
101.5
102.0
genus.Rothia.id.646

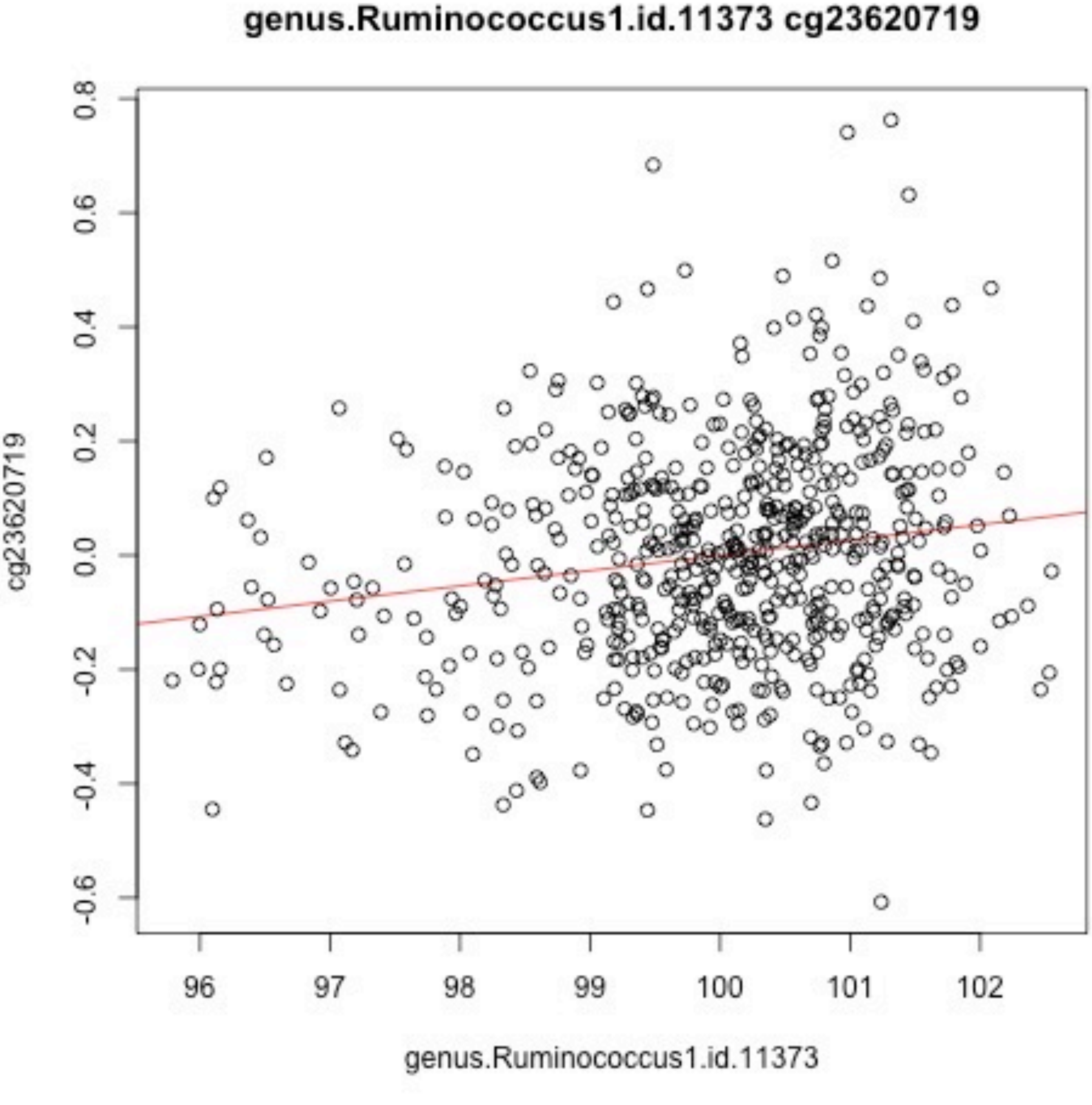
genus.Ruminococcus1.id.11373 cg23620719
cg23620719
0.8
0.6
0.4
0.2
0.0
-0.2
-0.4
-0.6
96
97
98
99
100
101
102
genus.Ruminococcus1.id.11373

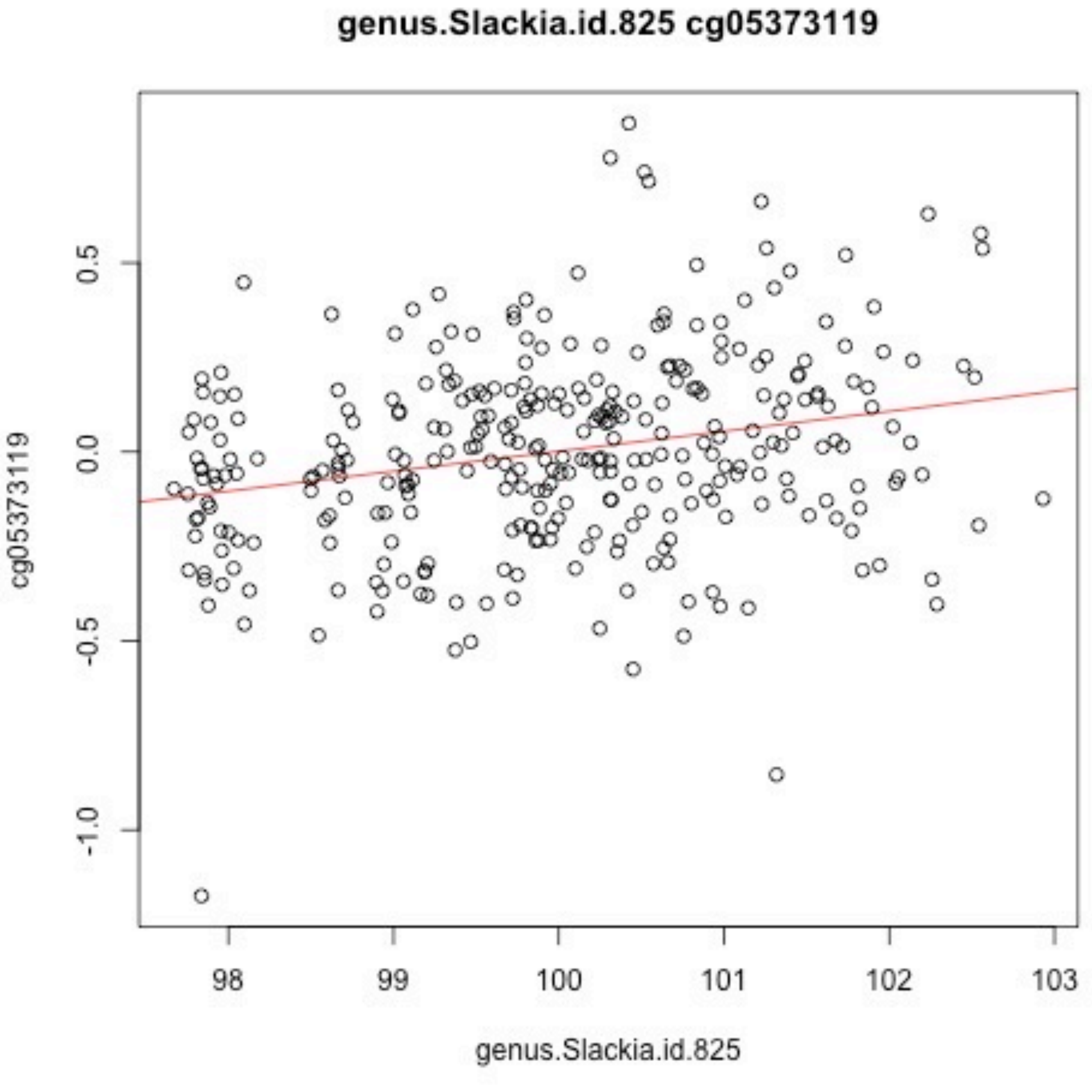
genus.Slackia.id.825 cg05373119
cg05373119
0.5
0.0
-0.5
-1.0
98
99
100
101
102
103
genus.Slackia.id.825

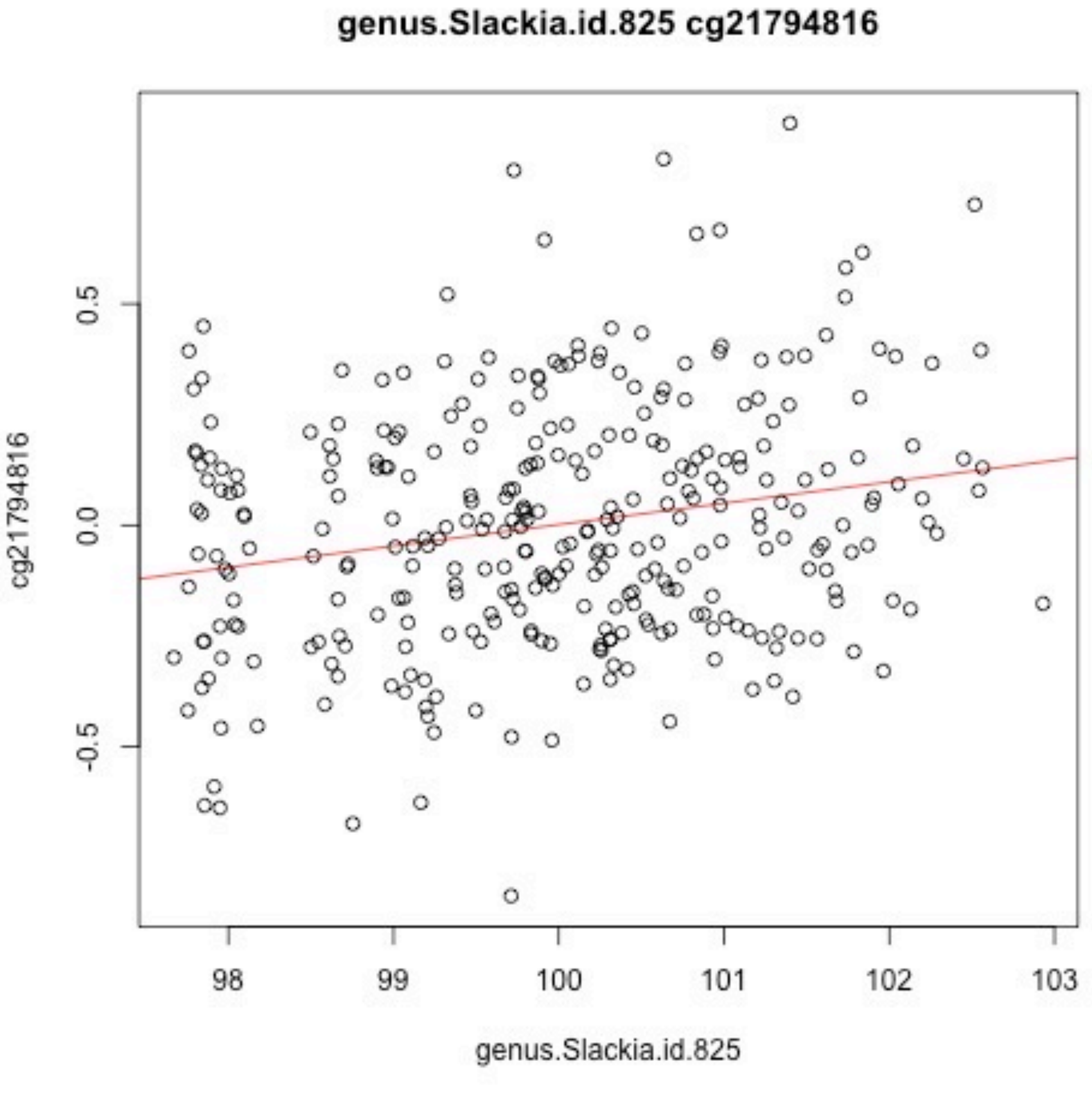
genus.Slackia.id.825 cg21794816
cg21794816
0.5
0.0
-0.5
98
99
100
101
102
103
genus.Slackia.id.825

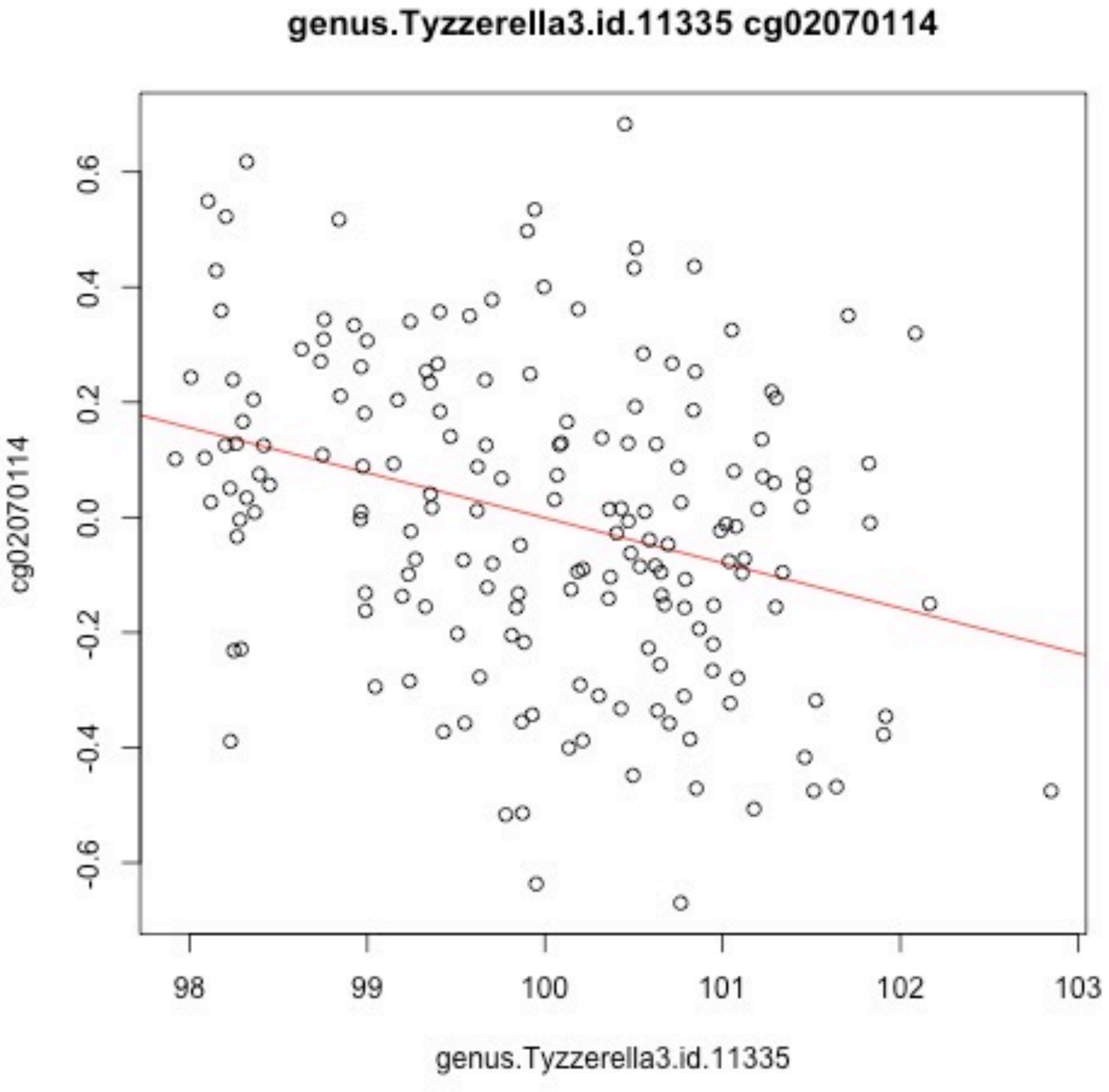
genus.Tyzzerella3.id.11335 cg02070114
cg02070114
0.6
0.4
0.2
0.0
-0.2
-0.4
-0.6
98
99
100
101
102
103
genus.Tyzzerella3.id.11335

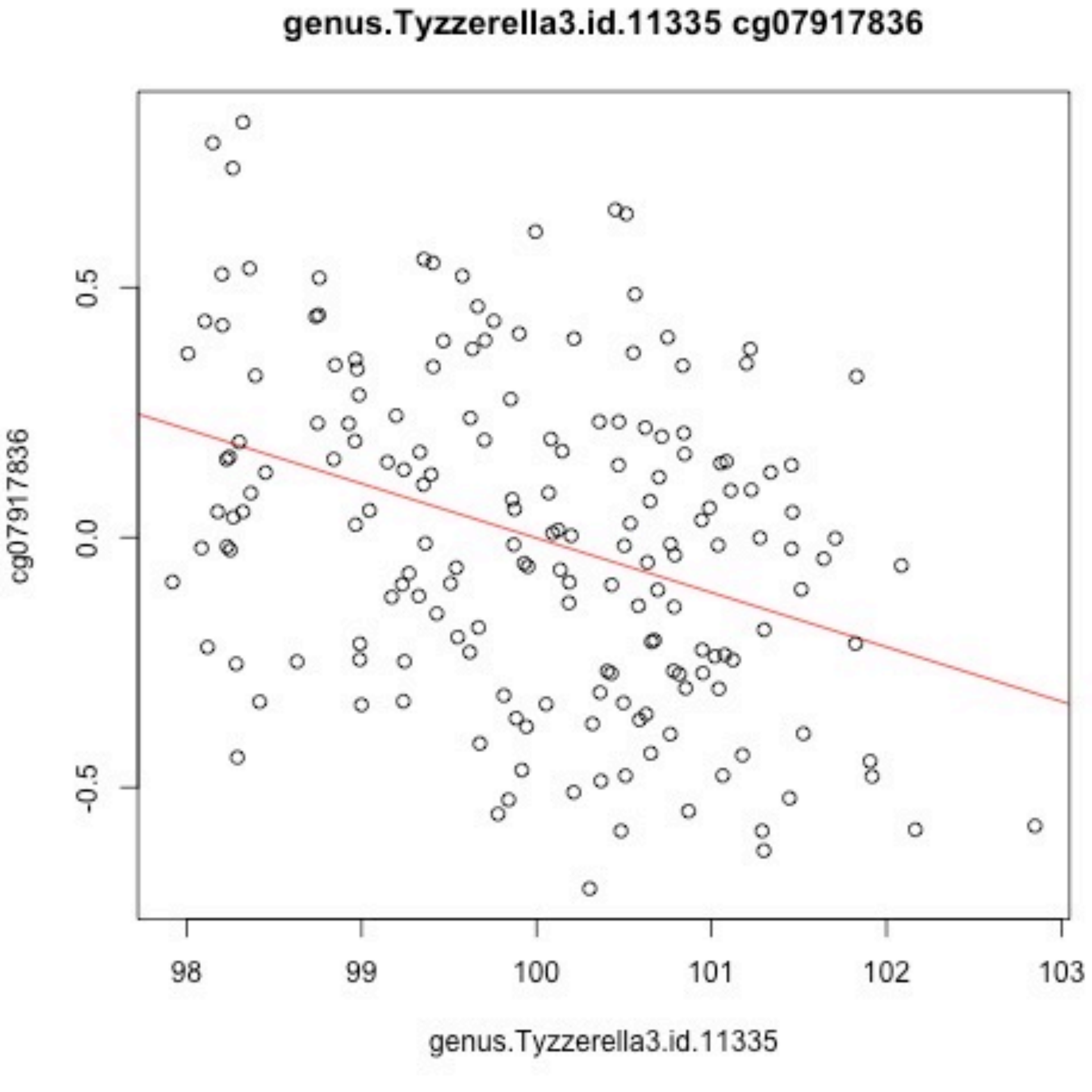
genus.Tyzzerella3.id.11335 cg07917836
cg07917836
0.5
0.0
-0.5
98
99
100
101
102
103
genus.Tyzzerella3.id.11335

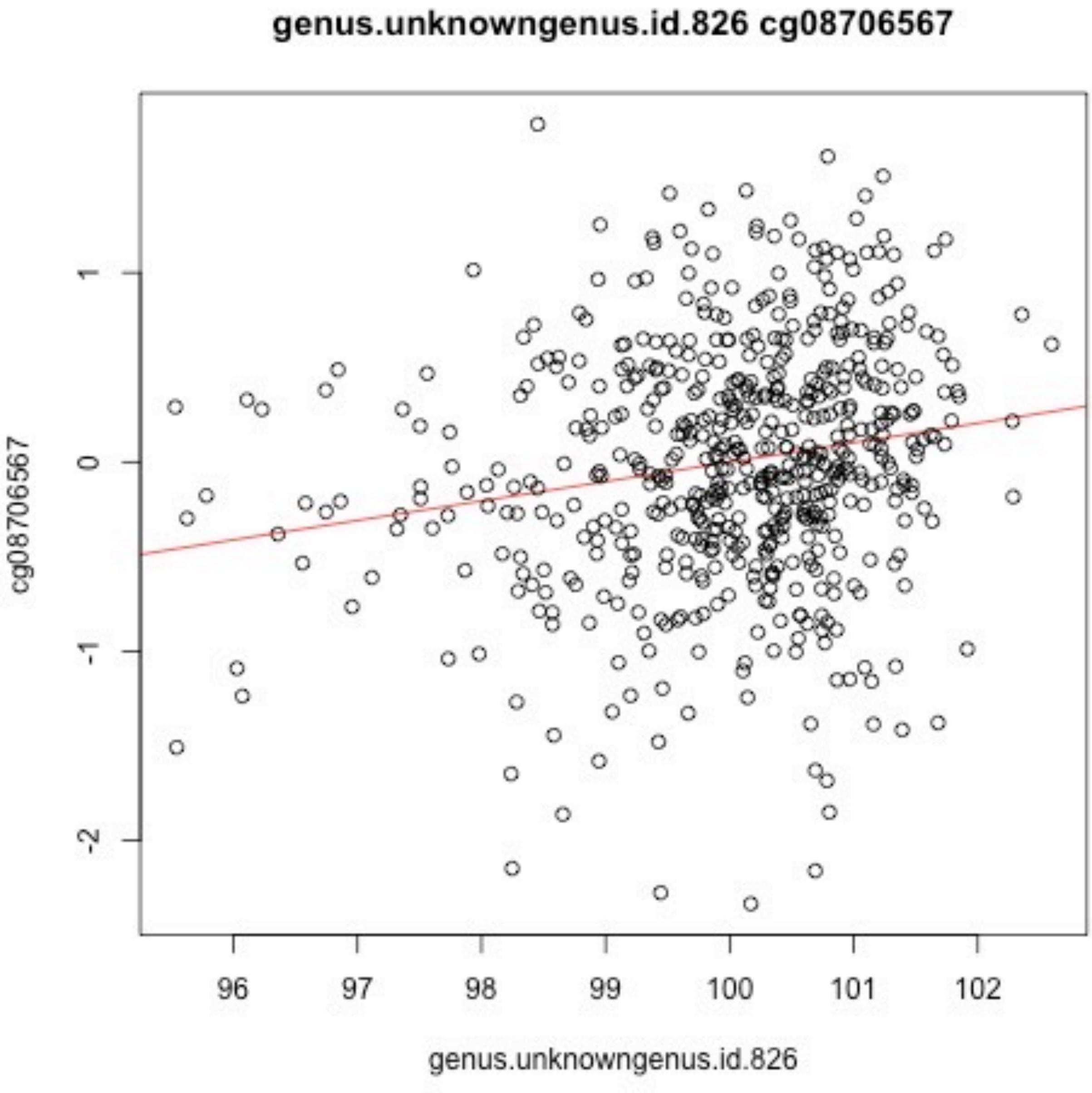

genus.unknowngenus.id.826 cg08706567
cg08706567
genus.unknowngenus.id.826
1
0
-1
-2
96
97
98
99
100
101
102

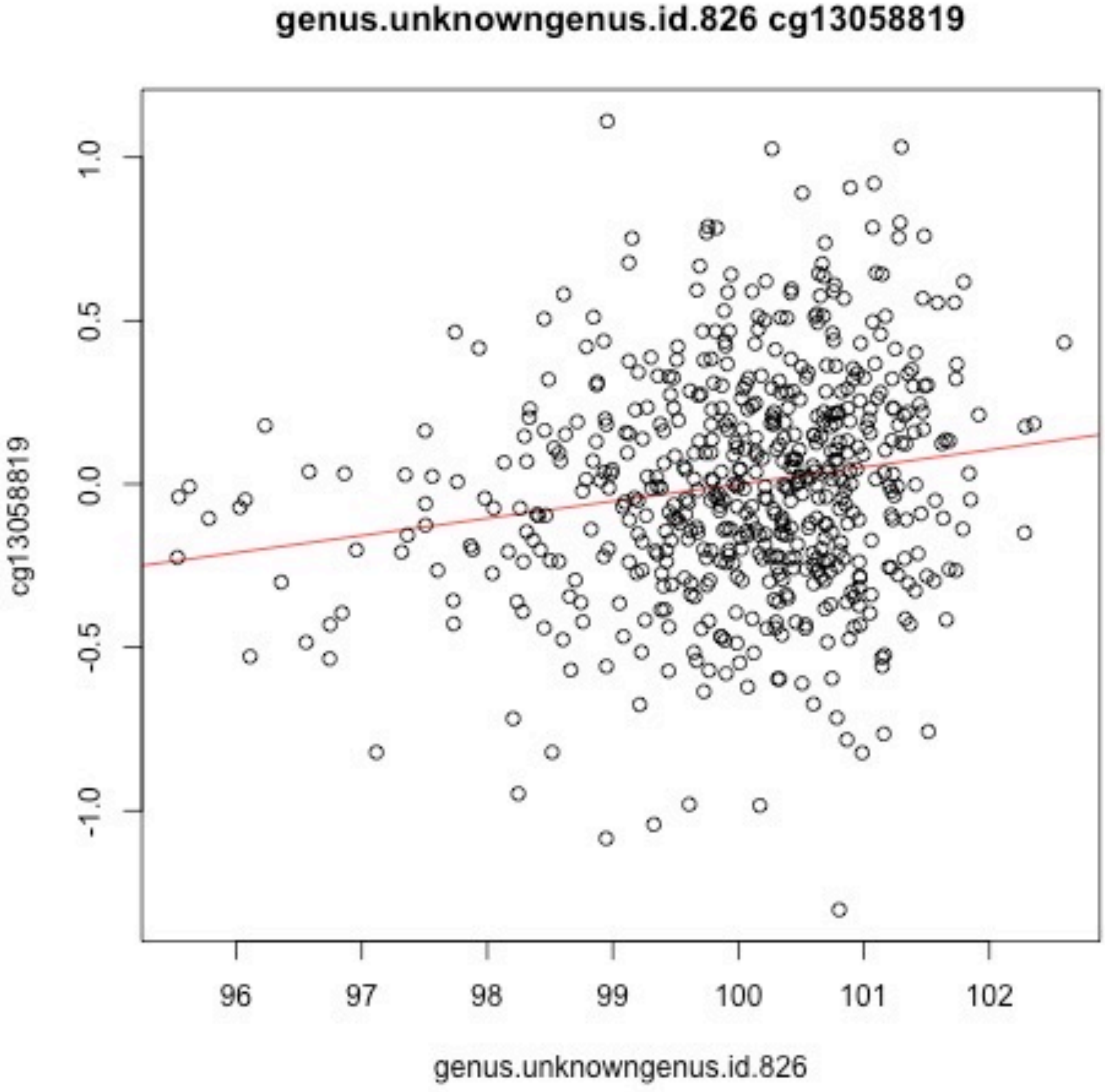
genus.unknowngenus.id.826 cg13058819
cg13058819
1.0
0.5
0.0
-0.5
-1.0
96
97
98
99
100
101
102
genus.unknowngenus.id.826

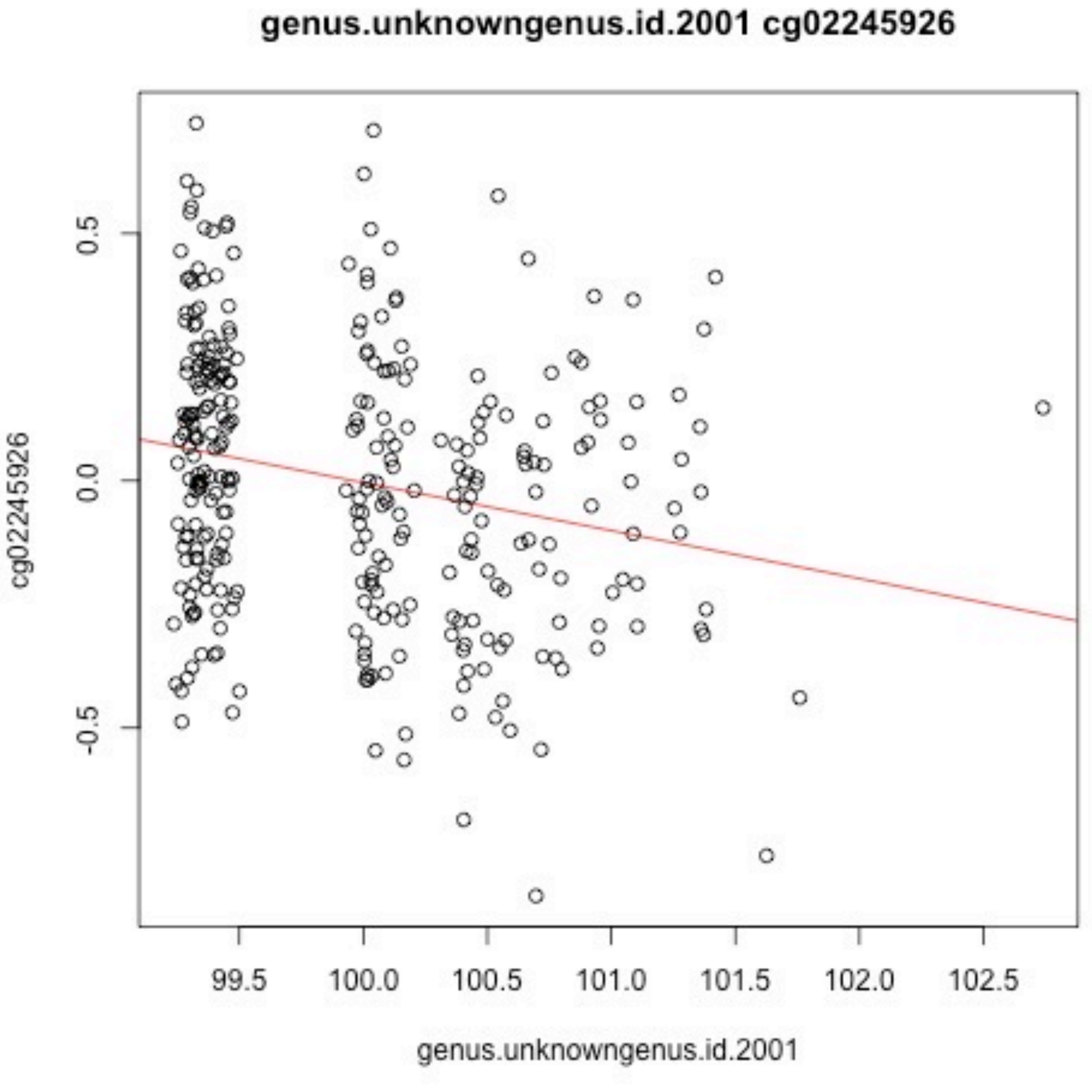
genus.unknowngenus.id.2001 cg02245926
cg02245926
0.5
0.0
-0.5
99.5
100.0
100.5
101.0
101.5
102.0
102.5
genus.unknowngenus.id.2001

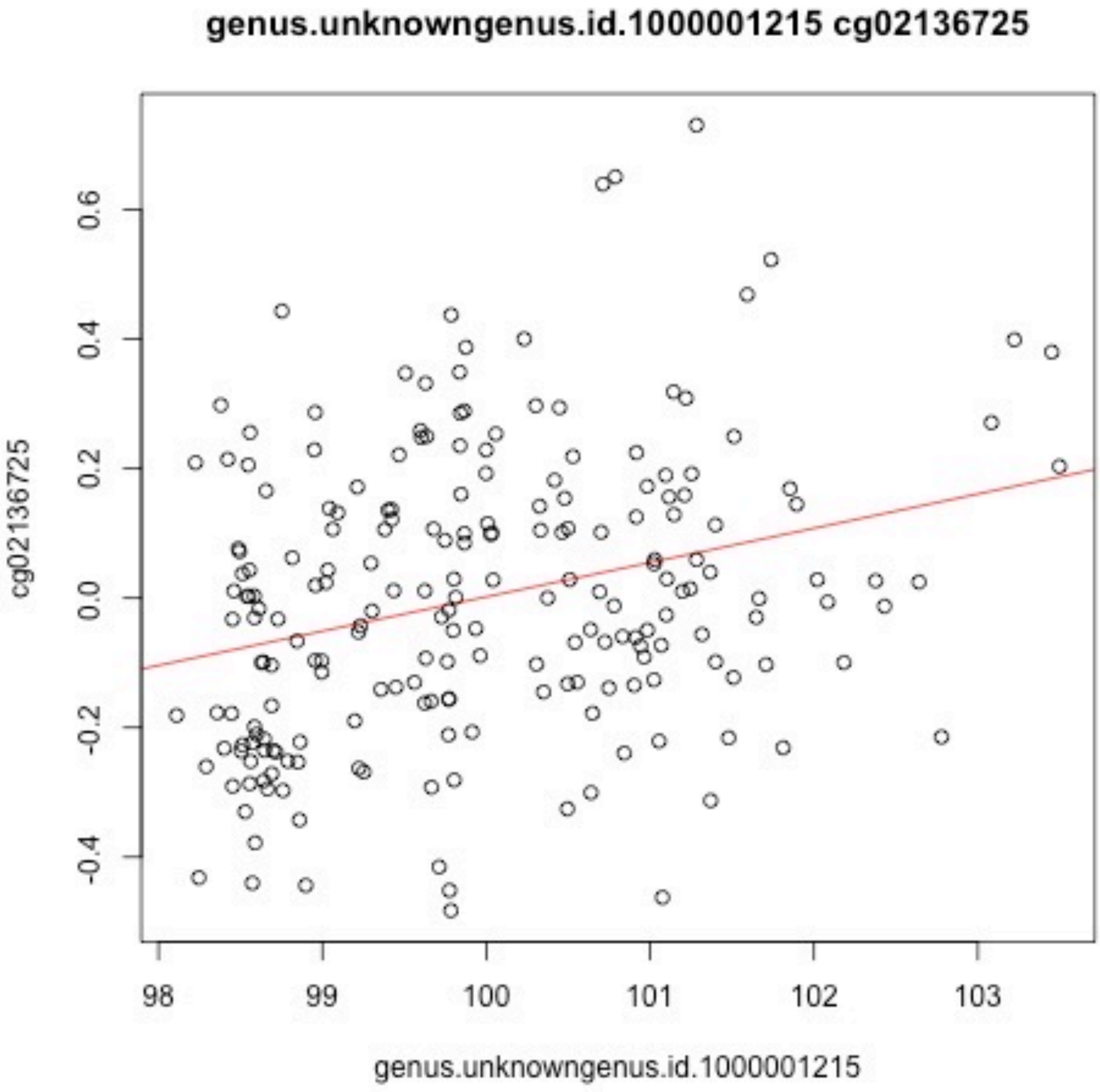
genus.unknowngenus.id.1000001215 cg02136725
cg02136725
-0.4
-0.2
0.0
0.2
0.4
0.6
98
99
100
101
102
103
genus.unknowngenus.id.1000001215

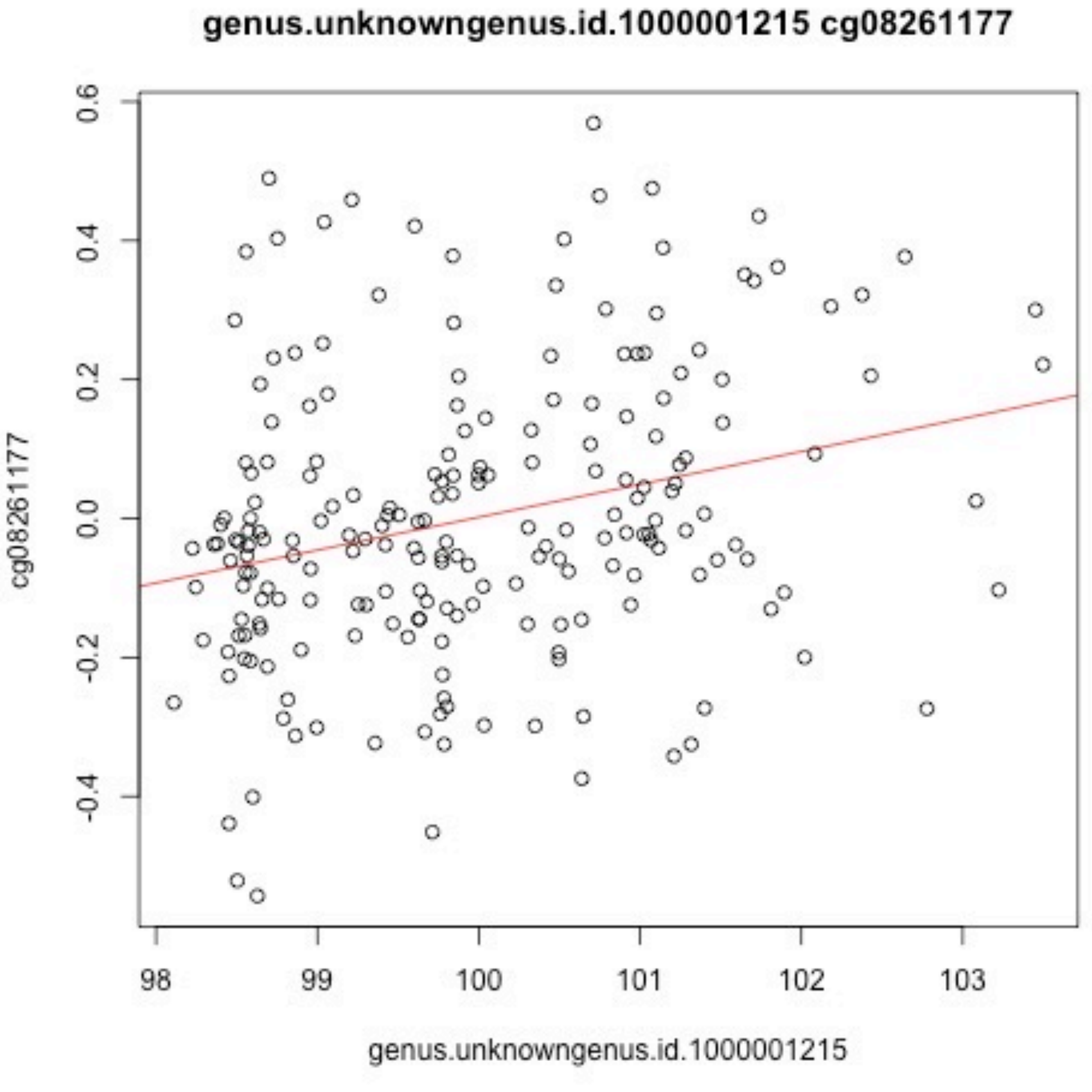

genus.unknowngenus.id.1000001215 cg08261177
cg08261177
0.6
0.4
0.2
0.0
-0.2
-0.4
98
99
100
101
102
103
genus.unknowngenus.id.1000001215

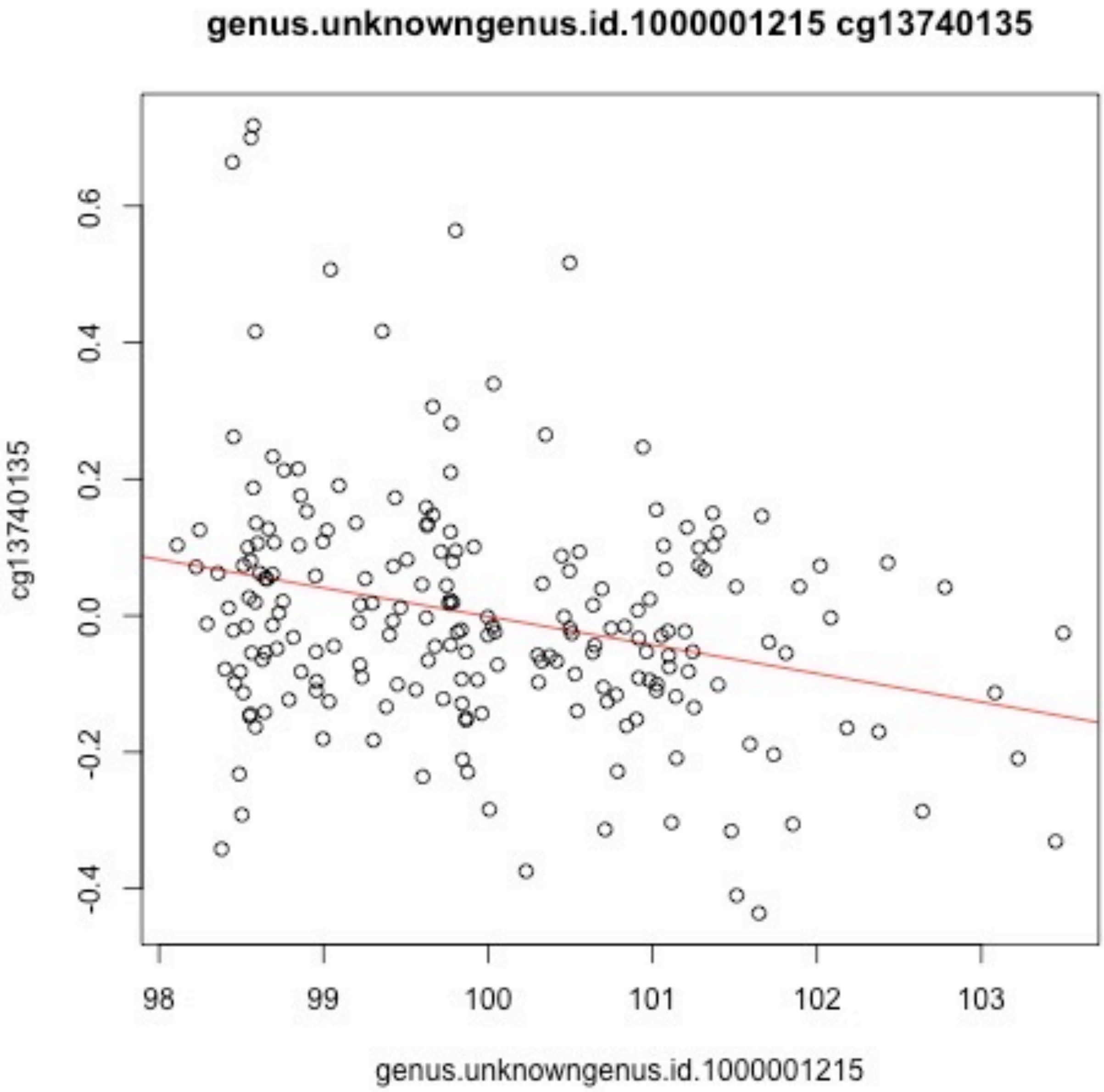
genus.unknowngenus.id.1000001215 cg13740135
cg13740135
0.6
0.4
0.2
0.0
-0.2
-0.4
98
99
100
101
102
103
genus.unknowngenus.id.1000001215

### genus.unknowngenus.id.1000005472 cg20893022

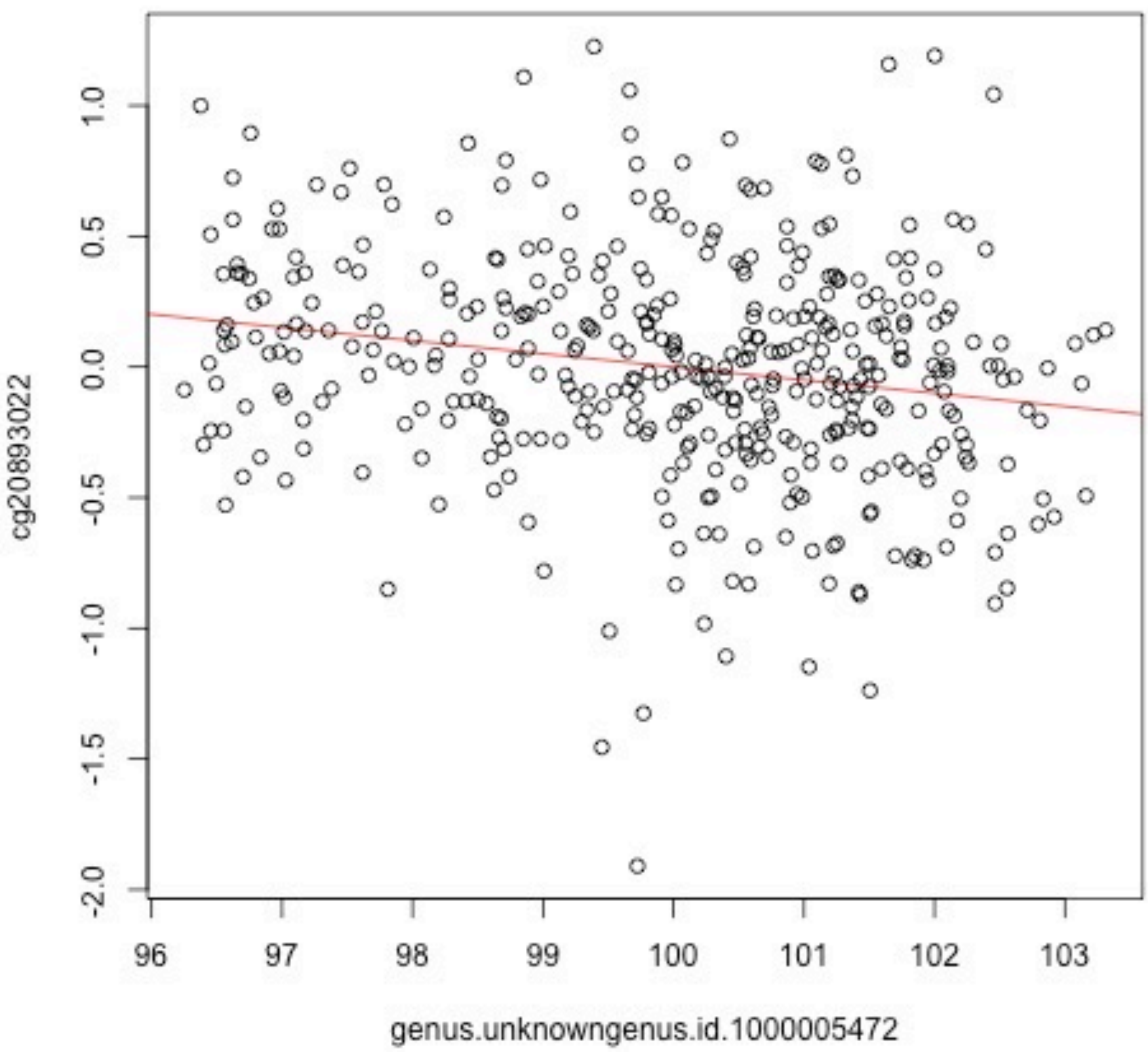

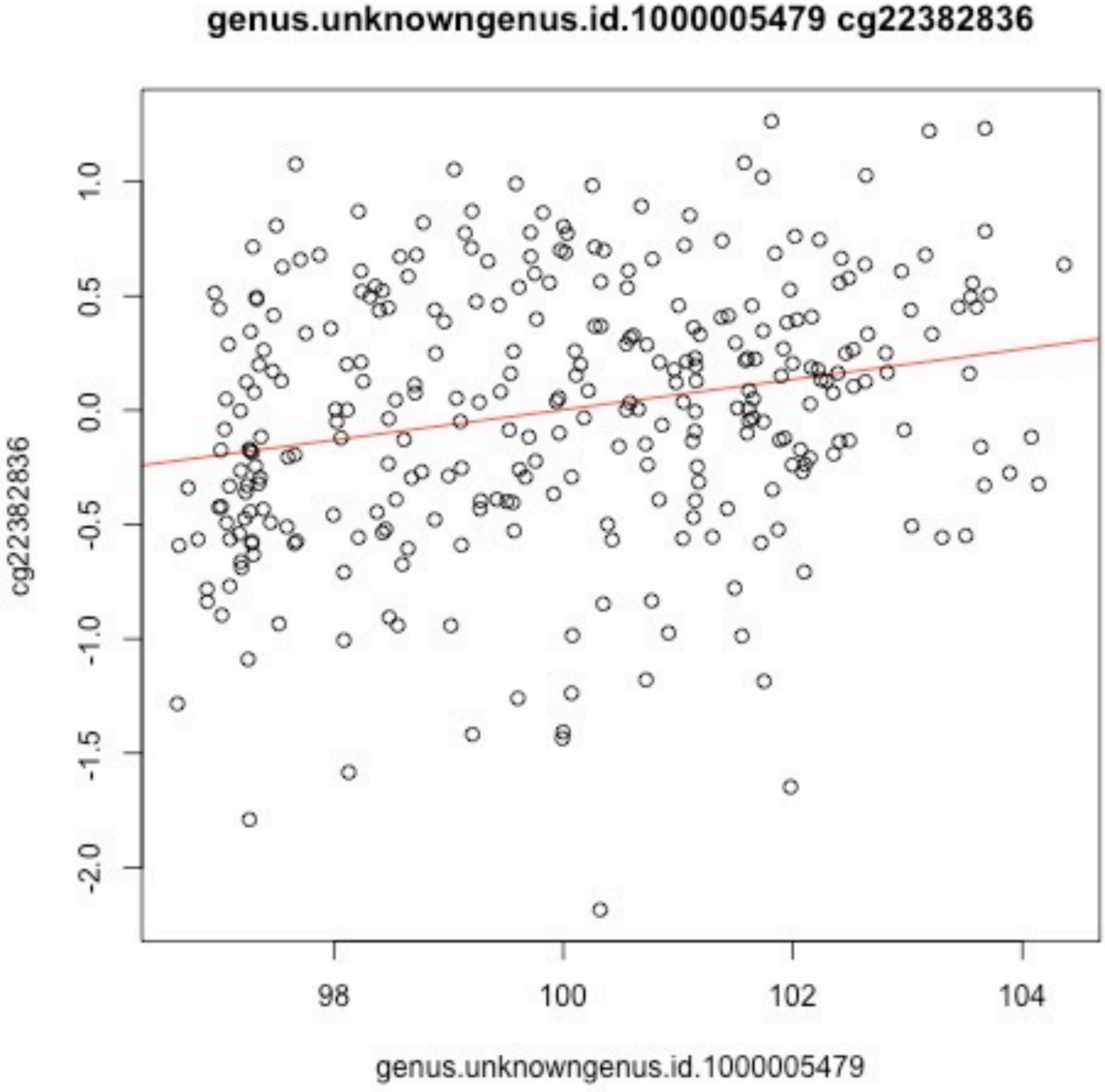

genus.unknowngenus.id.1000005479 cg22382836
cg22382836
1.0
0.5
0.0
-0.5
-1.0
-1.5
-2.0
98
100
102
104
genus.unknowngenus.id.1000005479

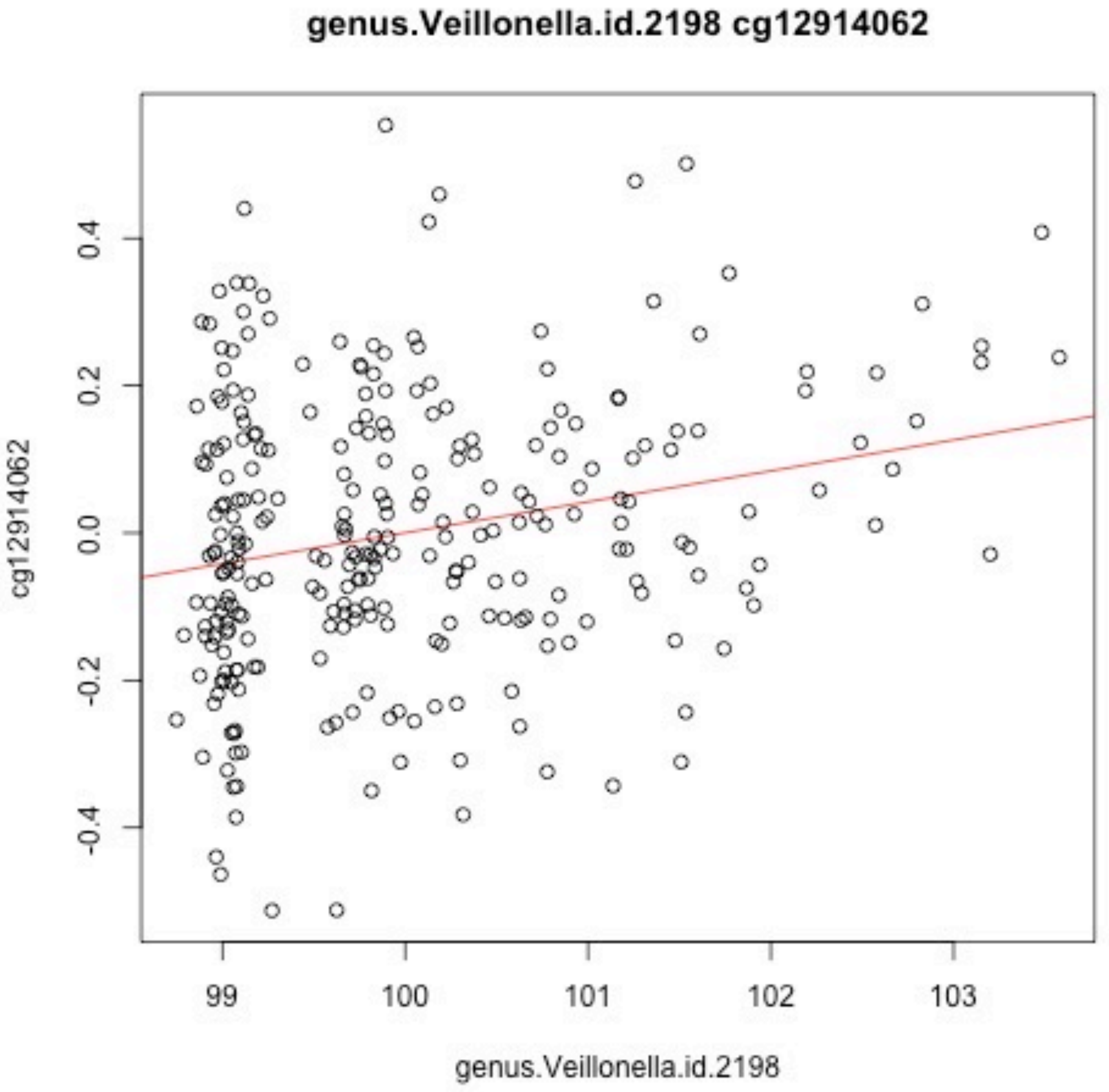
genus.Veillonella.id.2198 cg12914062
cg12914062
0.4
0.2
0.0
-0.2
-0.4
99
100
101
102
103
genus.Veillonella.id.2198

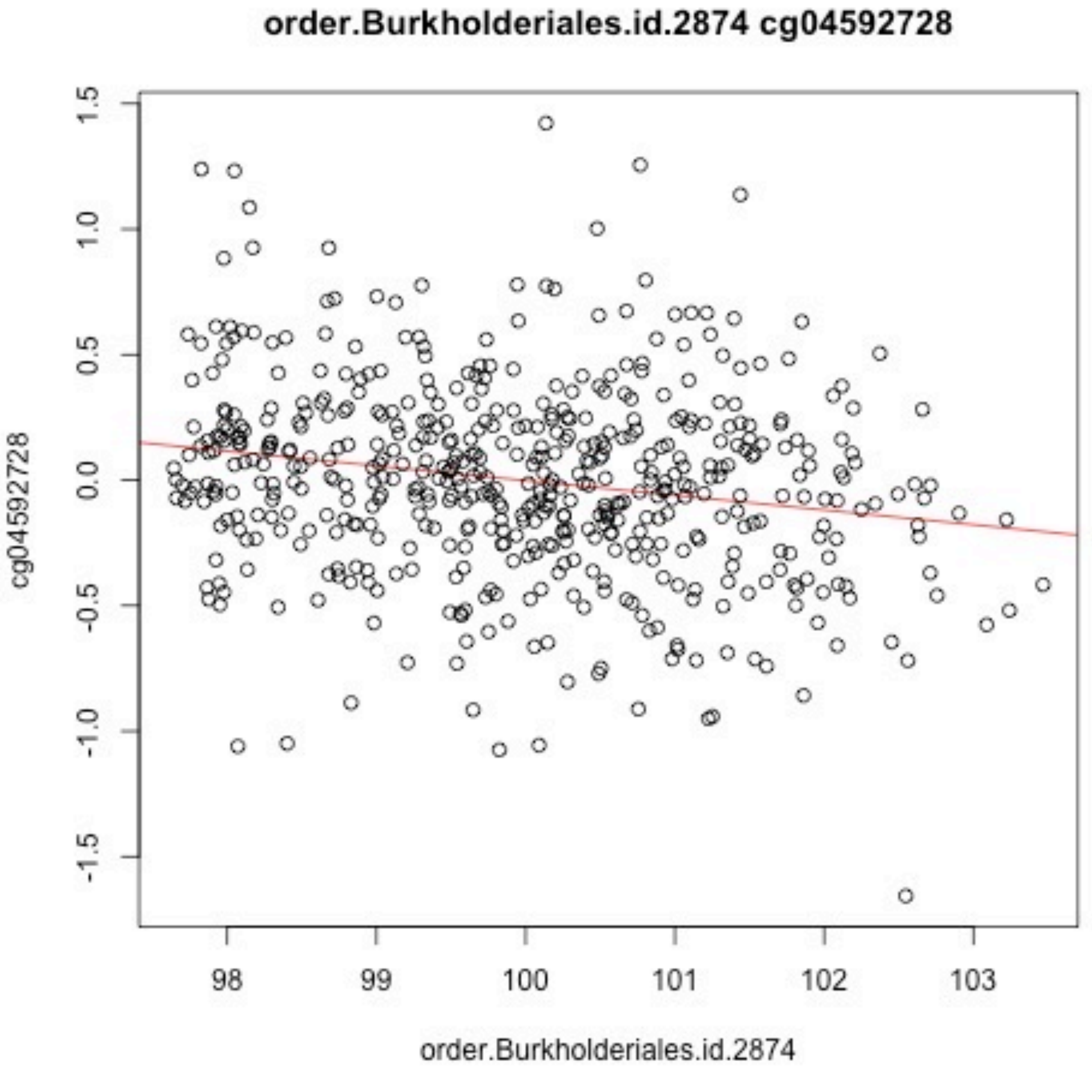
order.Burkholderiales.id.2874 cg04592728
cg04592728
1.5
1.0
0.5
0.0
-0.5
-1.0
-1.5
98
99
100
101
102
103
order.Burkholderiales.id.2874

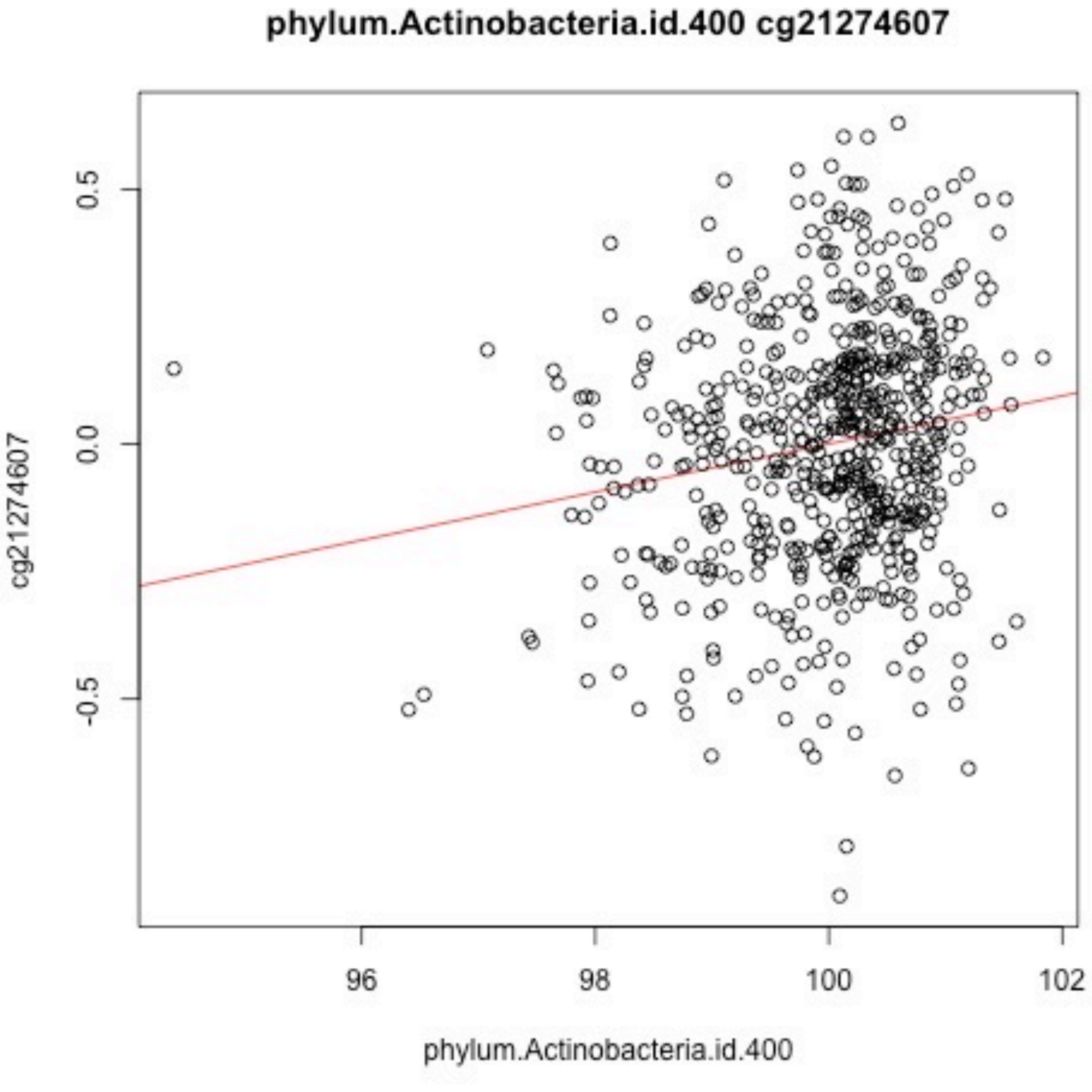
phylum.Actinobacteria.id.400 cg21274607
0.5
0.0
-0.5
cg21274607
96
98
100
102
phylum.Actinobacteria.id.400

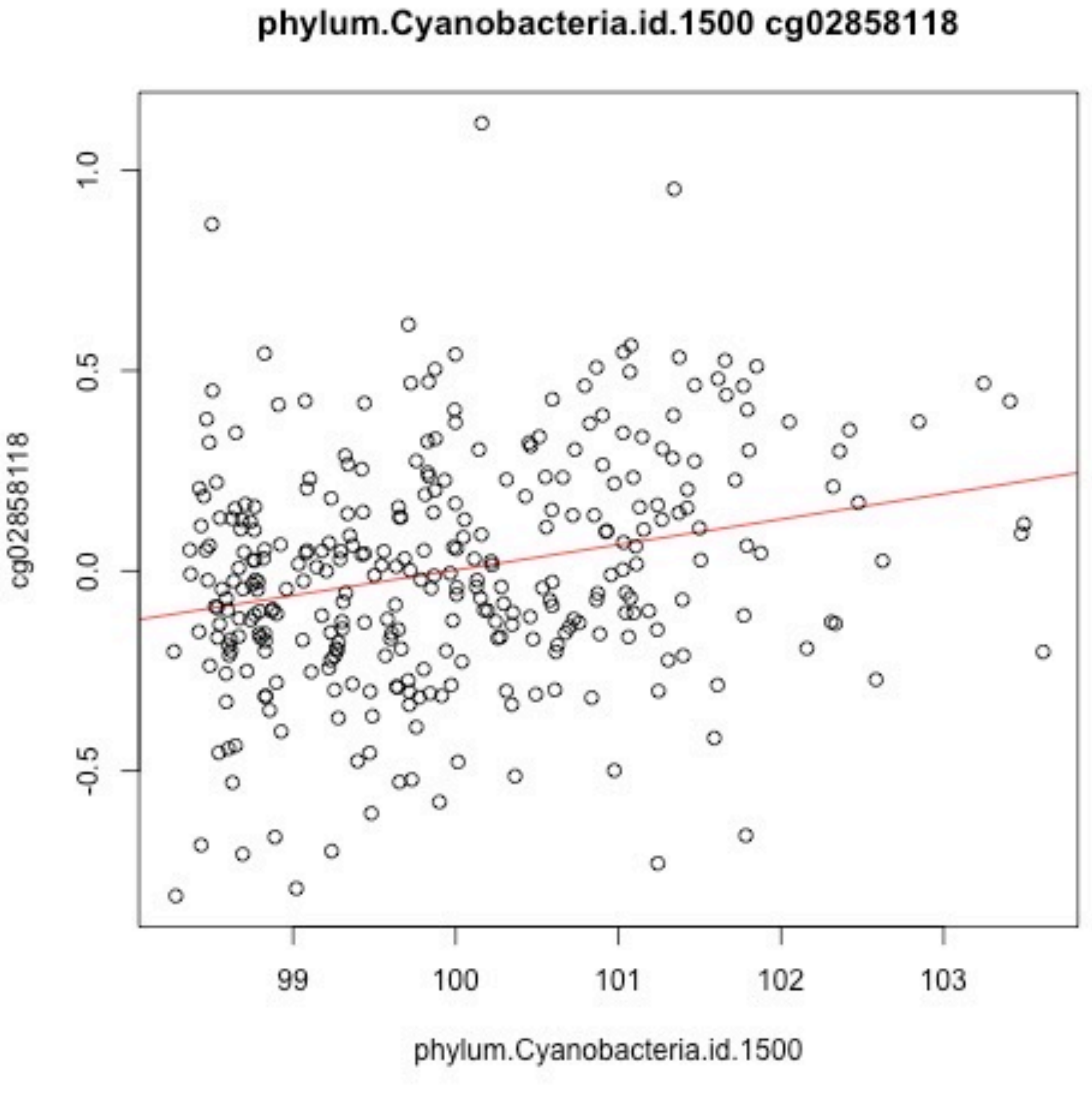
phylum.Cyanobacteria.id.1500 cg02858118
cg02858118
1.0
0.5
0.0
-0.5
99
100
101
102
103
phylum.Cyanobacteria.id.1500

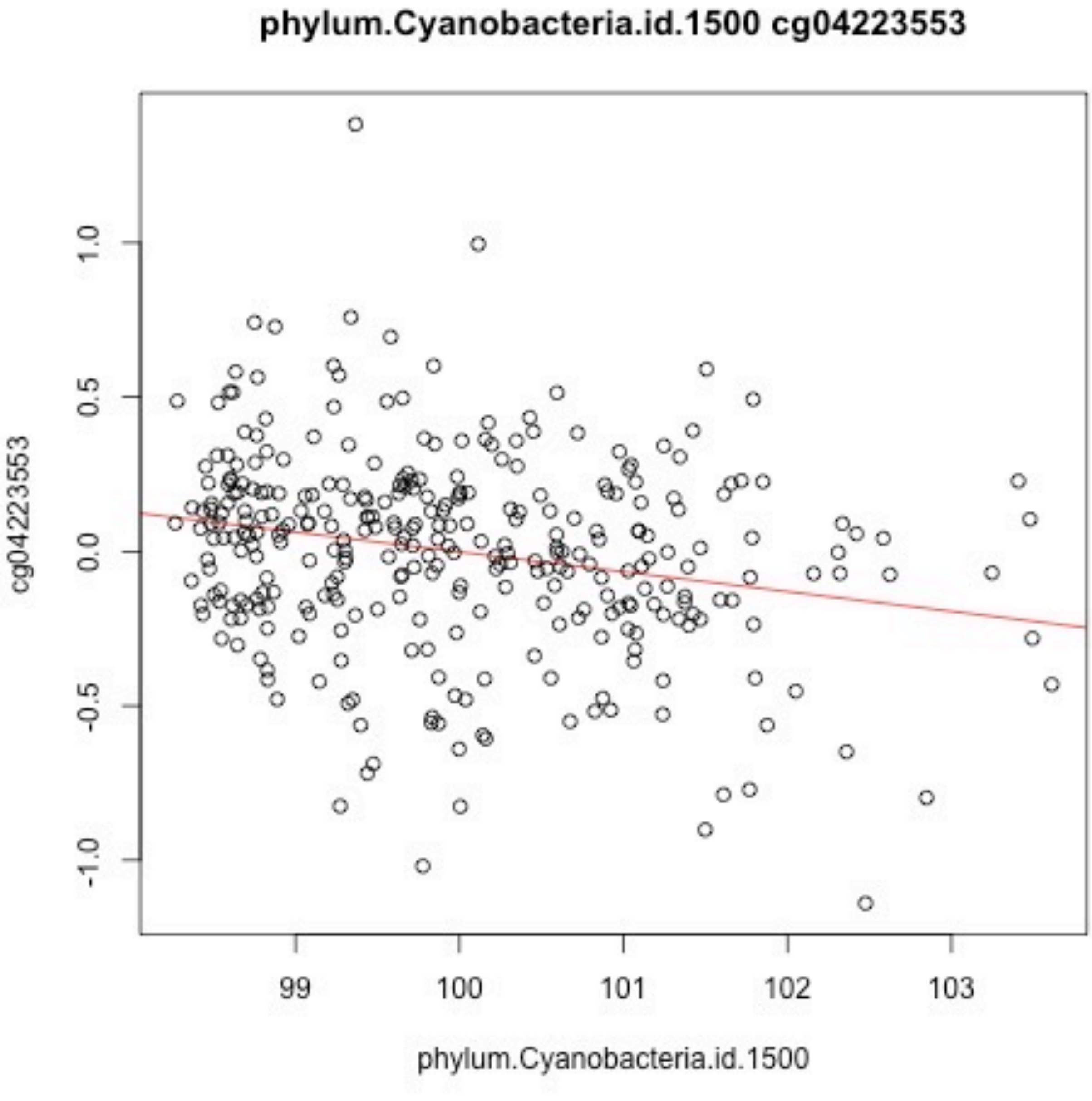
phylum.Cyanobacteria.id.1500 cg04223553
cg04223553
1.0
0.5
0.0
-0.5
-1.0
99
100
101
102
103
phylum.Cyanobacteria.id.1500

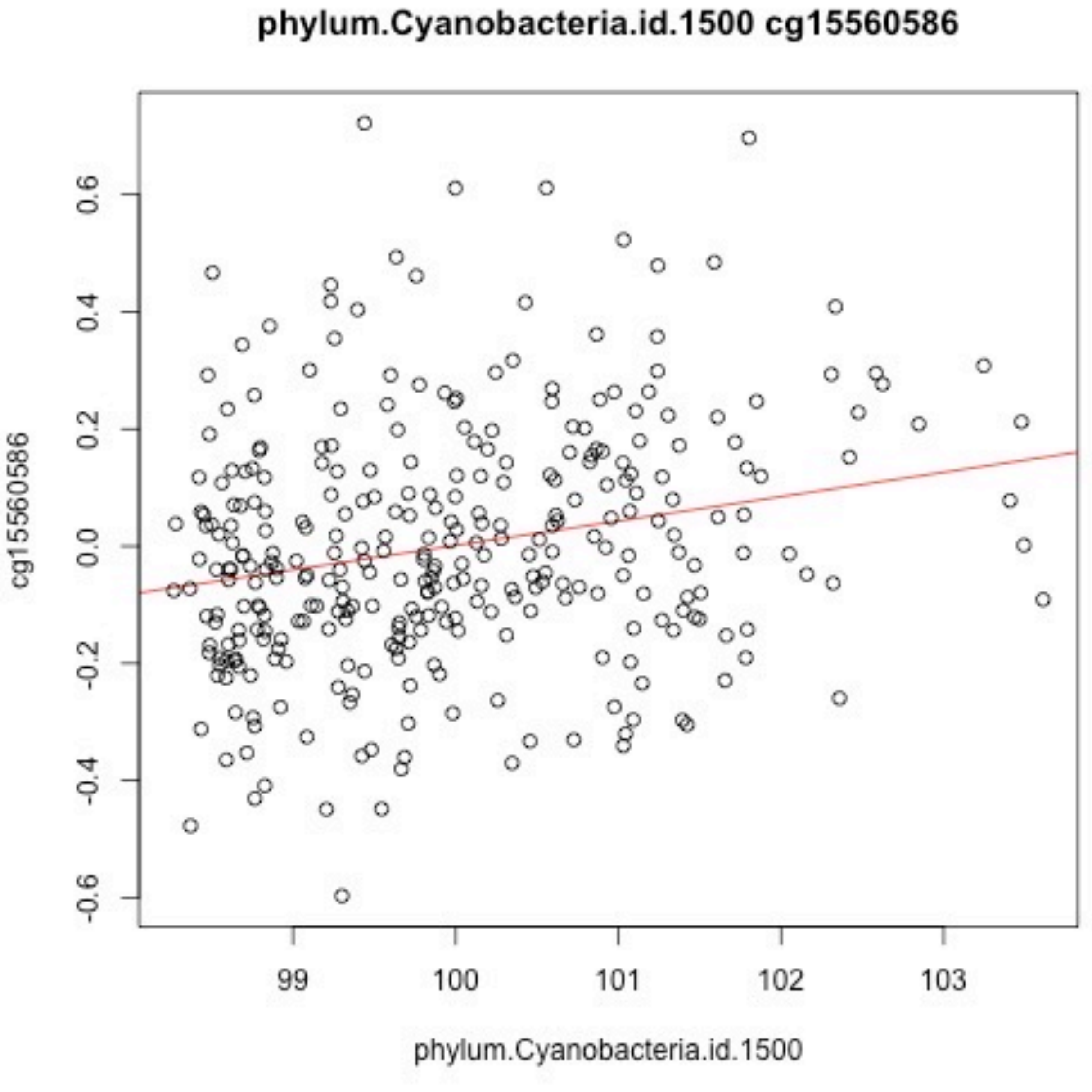
phylum.Cyanobacteria.id.1500 cg15560586
cg15560586
0.6
0.4
0.2
0.0
-0.2
-0.4
-0.6
99
100
101
102
103
phylum.Cyanobacteria.id.1500

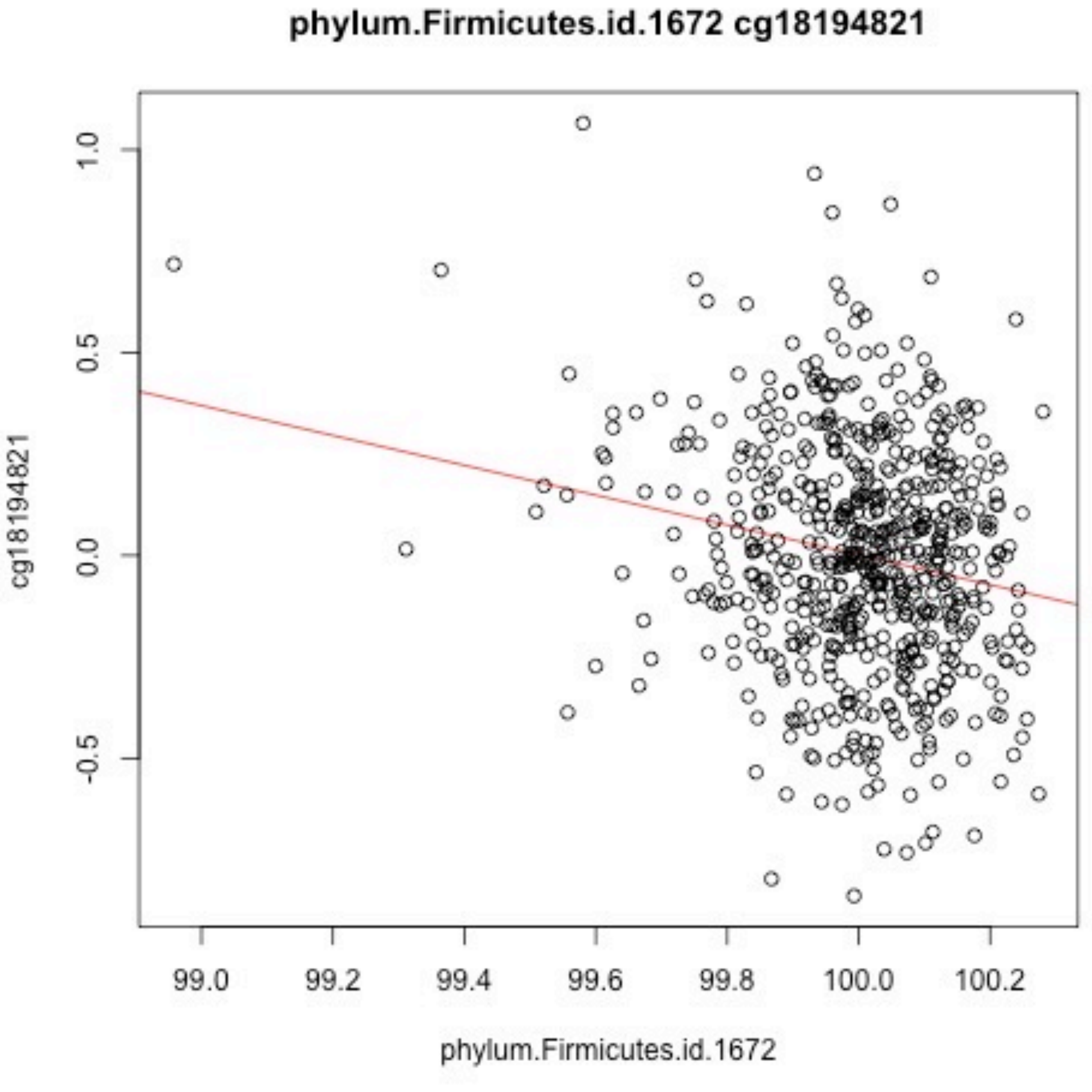
phylum.Firmicutes.id.1672 cg18194821
1.0
0.5
0.0
-0.5
cg18194821
99.0
99.2
99.4
99.6
99.8
100.0
100.2
phylum.Firmicutes.id.1672

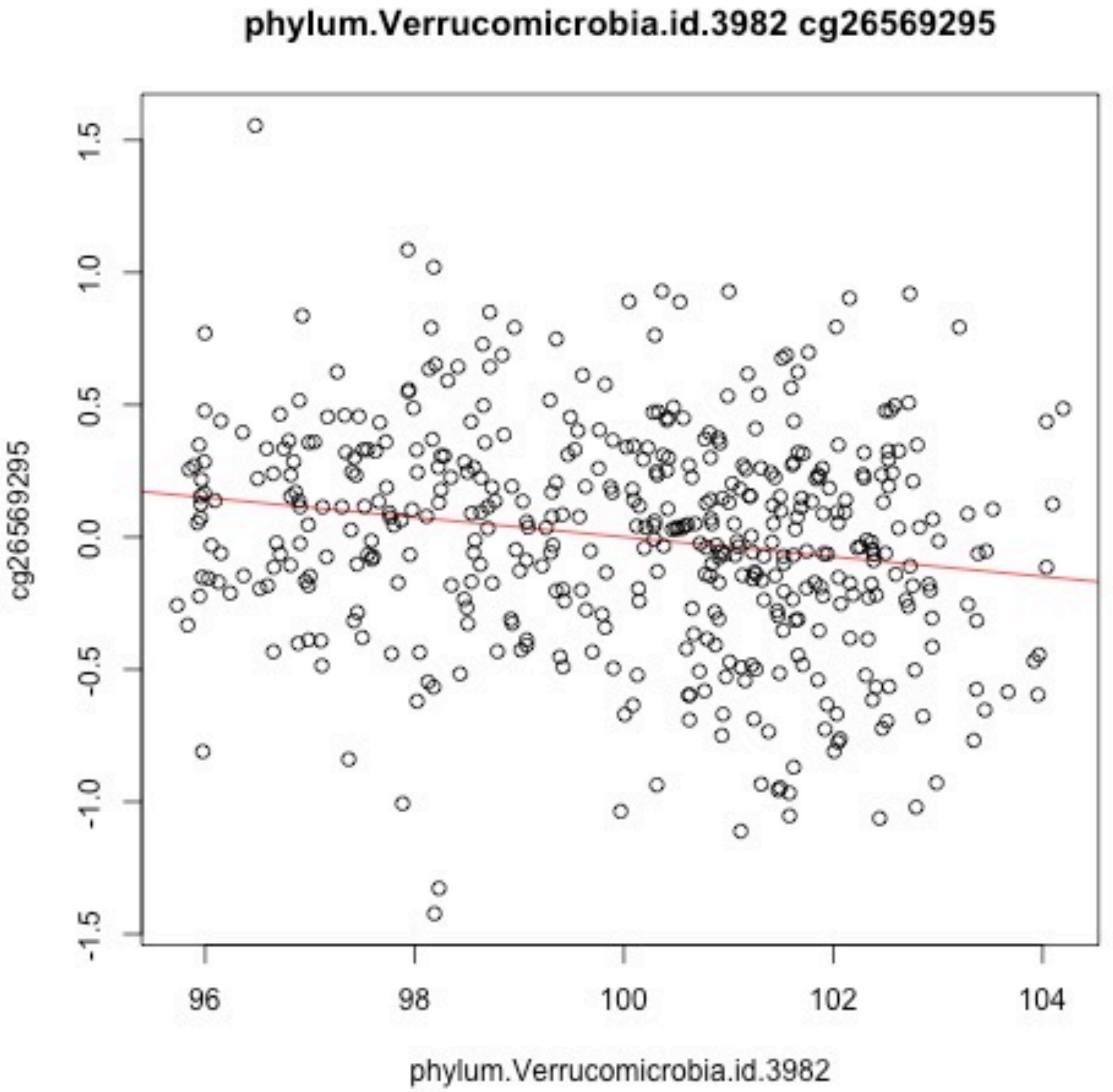

phylum.Verrucomicrobia.id.3982 cg26569295
cg26569295
1.5
1.0
0.5
0.0
-0.5
-1.0
-1.5
96
98
100
102
104
phylum.Verrucomicrobia.id.3982
